# Coevolution-induced stabilizing and destabilizing selection shapes species richness in clade co-diversification

**DOI:** 10.1101/2023.11.29.569146

**Authors:** Yichao Zeng, David H. Hembry

## Abstract

Coevolution can occur as a result of species interactions. However, it remains unclear how coevolutionary processes translate into the accumulation of species richness over macroevolutionary timescales. Assuming speciation occurs in a metacommunity as a result of genetic differentiation across communities due to dispersal limitation, we examine the effects of coevolution-induced stabilizing and destabilizing selection of a single quantitative trait on species diversification. We propose and test two hypotheses. (1) Stabilizing selection within communities enhances species diversification through strengthened dispersal limitation. (2) Destabilizing selection within communities impedes species diversification through weakened dispersal limitation. Here, we simulate clade co-diversification using an individual-based model, considering scenarios where phenotypic evolution is shaped by neutral dynamics, mutualistic coevolution, or antagonistic coevolution, where coevolution operates through trait matching or trait difference, and where the strength of coevolutionary selection is symmetrical or asymmetrical. Our assumption that interactions occur between an independent party (whose individuals can establish or persist in a community independently, e.g. hosts) and a dependent party (whose individuals cannot establish or persist in a community without the independent party, e.g. parasites or obligate mutualists) yields two contrasting results. Stabilizing selection within communities enhances species diversification in the dependent clade but not in the independent clade. Conversely, destabilizing selection within communities impedes species diversification in the independent clade but not in the dependent clade. These results are partially corroborated by empirical dispersal data, suggesting that these mechanisms might explain the diversification of some of the most species-rich clades in the Tree of Life.

## INTRODUCTION

The natural world is filled with interactions between different species. Many such interactions take the form of a bipartite partnerships in which two interaction parties play two distinct roles, e.g. the interactions between herbivores and plants, parasites and hosts, pollinators and angiosperms, and predators and prey. Bipartite interactions can potentially result in coevolution, that is, reciprocal evolutionary change in two or more interacting lineages driven by natural selection (Thompson 2005). Coevolutionary studies have focused on topics including phenotypic evolution across geographic space (Parchman and Benkman 2002), partnership specificity (Cook and Rasplus 2003), and signatures of coevolution in community assembly (Endara et al. 2017). Two coevolving lineages can co-diversify, i.e. diversify simultaneously, when they are allowed enough time. This has been an active area of research since the “escape-and-radiate” hypothesis, which states that rapid diversification of phytophagous insects or their independent plants often follows a significant shift in interaction partners (Ehrlich and Raven 1964; Cogni, Quental & Guimarães 2022). The ways by which macroevolutionary dynamics can be generated by processes at the local scale can be understood in the light of the geographic mosaic theory of coevolution, which emphasizes that coevolutionary dynamics can occur between geographically connected populations (Thompson 2005). It is therefore plausible that speciation could arise out of coevolutionary mosaics (Hembry, Yoder, and Goodman 2014; Thompson, Segraves, and Althoff 2017). Specifically, local coadaptation could potentially decrease the chance of mating between populations, paving the way for reproductive isolation and consequently speciation (Thompson, Segraves, and Althoff 2017). However, whether speciation is more likely to be enhanced or impeded by coevolution remains unclear (Janz 2011; Harmon et al. 2019; Hembry and Weber 2020).

Coevolution is shaped by the fitness outcomes of species interactions, which can vary in at least three ways. First, species interactions may benefit both parties or benefits one party at the cost of the other, affecting differently the fitness and, consequently, the coevolutionary outcome. Second, coevolution can be characterized by whether they are mediated by trait difference or trait matching (Yoder and Nuismer 2010). Under the trait-difference scenario, a species’ fitness is maximized when its trait value is very different from that of its interaction partner. Examples of the trait-difference scenario (in the sense of Yoder and Nuismer 2010) include antagonistic interactions in which predators and prey evolve increasingly strong weaponry and defense against each other (Vermeij 1994) as well as mutualistic arms races in which both partners gain more benefits from their partner when they have a greater trait value than that of their interaction partner, e.g., a longer tongue in moth pollinators and a longer nectary spur in flowers (Whittall and Hodges 2007). Under the trait-matching scenario (in the sense of Yoder and Nuismer 2010), on the other hand, a species’ coevolutionary fitness is maximized when its trait value more closely matches that of its interaction partner. Antagonistic examples of this scenario include brood parasitism (Langmore, Hunt, and Kilner 2003) whereas mutualistic examples include obligate pollination mutualisms (Althoff and Segraves 2022). Third, the strength of natural selection imposed by a species interaction is not always symmetrical for the interacting parties, with the selection often being much stronger on one party than on the other (Brodie and Brodie 1999; Andreazzi, Thompson, and Guimaraes 2017; Endara et al. 2017).

Different selective regimes (i.e. modes of phenotypic evolution) can potentially arise from the different fitness outcomes of species interactions mentioned above. Prior literature has focused on the microevolutionary effects of different selective regimes on coevolution (Yoder and Nuismer 2010; Hembry et al. 2014). However, a theory is lacking on whether and how different modes of phenotypic selection induced by coevolution drive or impede the accumulation of species richness. The geographic mosaic theory of coevolution points to the possibility of coevolutionary diversification of clades while leaving the exact mechanisms largely unexplored (Thompson 2005). Given that certain types of species interactions are central to the biology of many organisms (e.g. herbivory to the biology of phytophagous insects), a better understanding of how coevolution shapes diversification through phenotypic evolution along this line could potentially explain the diversification of many clades. Here we consider a general mode of speciation where dispersal limitation causes genetic differentiation across space, eventually leading to speciation (Moran 1962; Etienne & Alonso 2005; Rosindell, Harmon & Etienne 2015; Manceau, Lambert, Morlon 2015; Hubert et al. 2015; Maliet, Loeuille & Morlon 2020). Based on this mode of speciation, our model considers how within-community coevolutionary dynamics shapes between-community dispersal dynamics and eventually shapes speciation dynamics across an entire metacommunity (Fig. 1).

**Figure 1.**
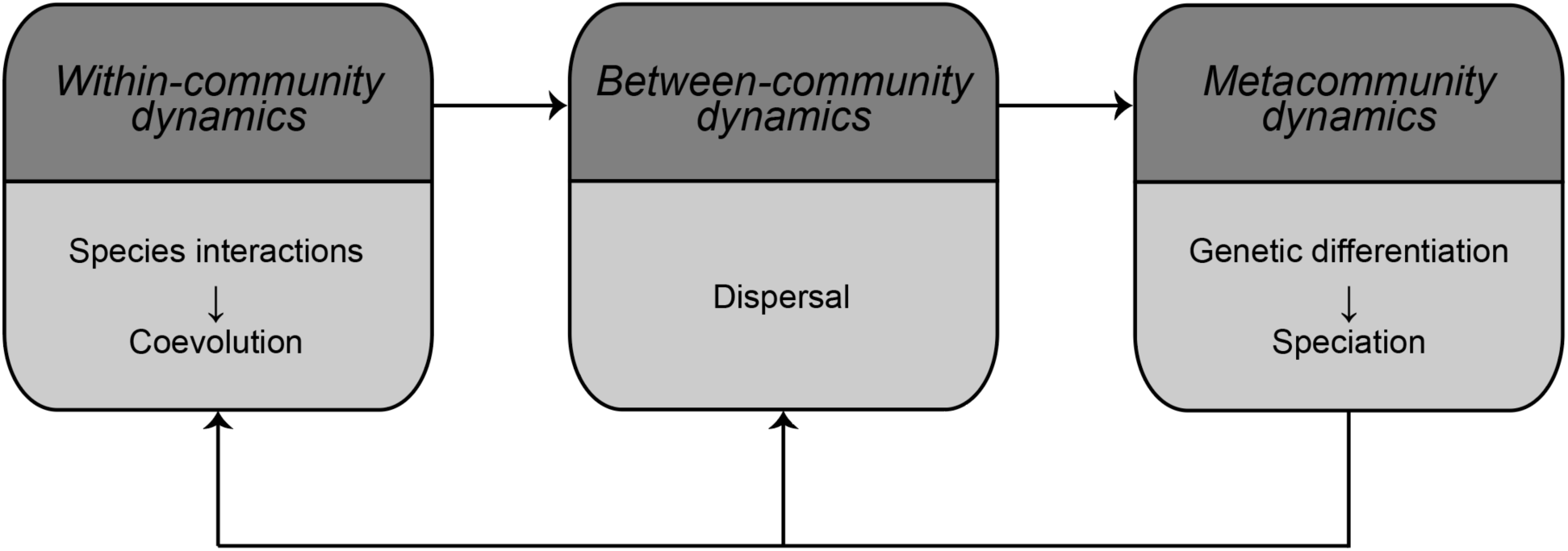
Our model describes the interplay between processes at three scales: within-community, between community, and across a metacommunity. Within community, species interactions can result in coevolution and thus generate selective pressure for individuals of belonging to both interacting parties. Between communities, individuals disperse and exchange genes. Across an entire metacommunity, genetic differentiation occurs between different communities and eventually results in speciation.

We propose two novel hypotheses regarding the different ways by which within-community coevolution shapes between-community dispersal and consequently shapes species across an entire metacommunity (Fig. 2). Over temporal scales, stabilizing selection is characterized by increased temporal trait stability compared to neutral drift, and destabilizing selection can conversely be defined as a selective regime that decreases temporal trait stability (Gingerich 2019a; Gingerich 2019b). The idea that within-population stabilizing selection limits between-population dispersal has been supported both empirically and theoretically (Tufto 2000; Tufto 2001; Lopez el al. 2008; Scheepens, Frei & Stöcklin 2010; Yeaman & Whitlock 2011; Huisman and Tufto 2012; Zacchello, Vinyeta, and Ågren 2020). This leads to the intuitive notion that, the stronger temporal trait stability a selective regime causes, the more strongly dispersal is limited – and a stronger degree of dispersal limitation causes stronger genetic differentiation across space and eventually a higher species richness accumulation. Following this intuition, we propose a first hypothesis termed the *stabilizing selection hypothesis*: stabilizing selection impedes dispersal (and thus increases the degree of dispersal limitation), which results in an increase in genetic differentiation across space, eventually resulting in an increase in the species richness accumulation. The same intuition can lead to another hypothesis, which is termed the *destabilizing selection hypothesis*: destabilizing selection enhances dispersal (and thus reduces the degree of dispersal limitation), which results in a reduction in genetic differentiation across space, eventually resulting in a reduction in the species richness accumulation.

**Figure 2.**
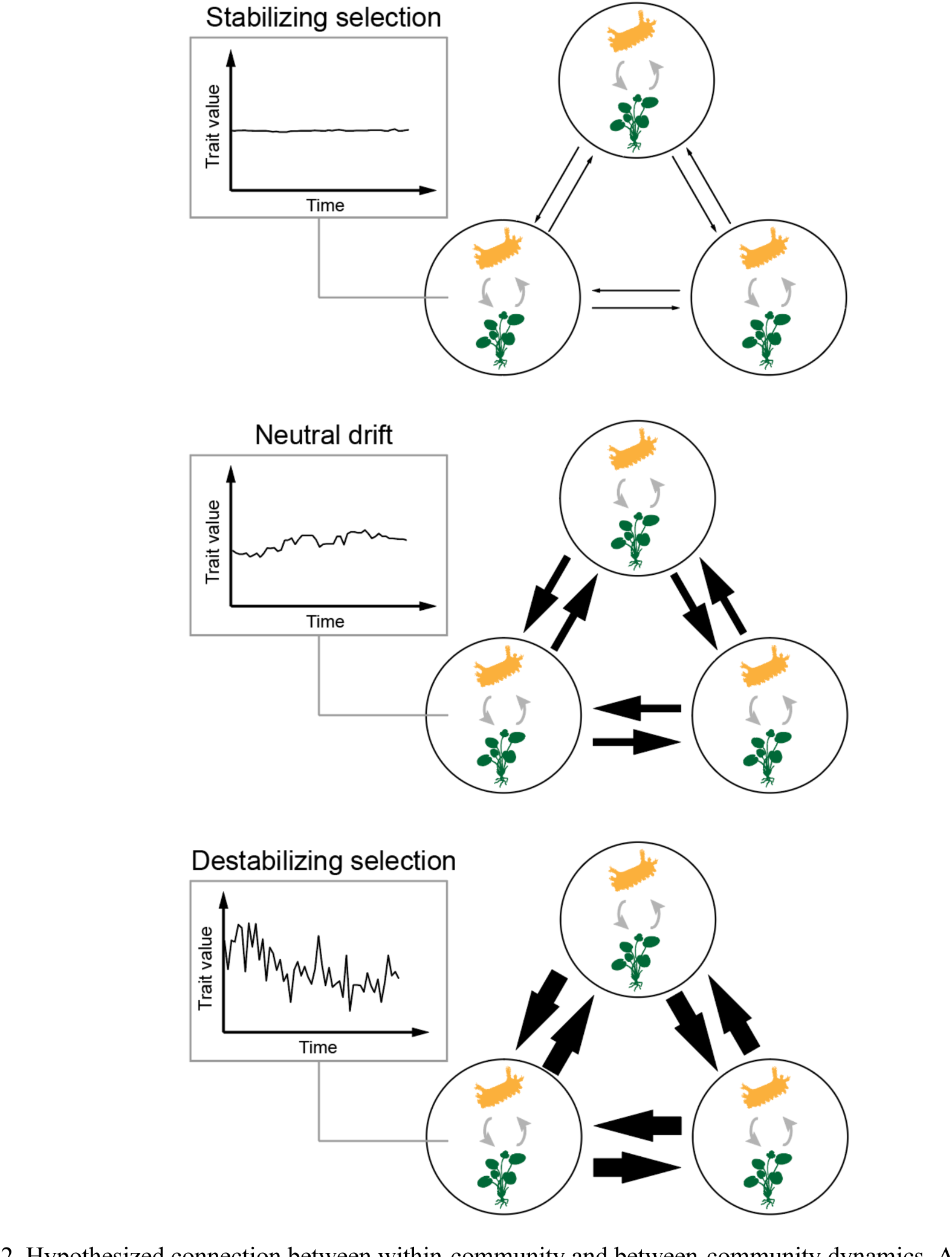
Hypothesized connection between within-community and between-community dynamics. A metacommunity is composed of multiple communities (circles) connected through dispersal (straight arrows). Species interactions (curved arrows) occur within each community. The size of a straight arrow indicates the level of dispersal. Compared to neutral drift as the baseline, stabilizing selection is hypothesized to impede dispersal whereas destabilizing selection is hypothesized to enhance dispersal.

To test these hypotheses in the context of two coevolving clades, we built an individual-based model for the coevolutionary accumulation of species richness across two-dimensional space. We considered different coevolutionary scenarios, including those where the interaction is mutualistic versus antagonistic, where the interaction is mediated by trait differences versus trait matching, and where coevolutionary selection is symmetrically versus asymmetrically strong for the interacting parties. In bipartite interactions such as antagonisms and mutualisms, one partner is often more dependent on the other than the other way around. Specifically, many antagonistic interactions occur between antagonistic symbionts (e.g. parasites, pathogens) and their hosts without which they cannot survive (Schmid-Hempel 2013). Even in mutualistic interactions, the mutual dependence of interaction partners can be highly asymmetric (Bascompte, Jordano, and Olesen 2006; Bronstein 2015). Given this, we built into our model the differences between independence and dependence in partnership, assuming that the independent party’s individuals, e.g. hosts, can establish or persist in a community independently, whereas a dependent party’s individuals cannot establish or persist in a community without the independent party, e.g. parasites or obligate mutualists. Our simulation results suggest mechanisms for the diversification of some of the most species-rich groups in the Tree of Life (Table 1).

**Table 1.** Examples of species interactions that may be described as independent-dependent partnerships and the species richness estimates for the independent and dependent parties.

| Interaction | Independent party<br>(species richness) | Dependent party (species<br>richness) | Reference(s) |
| --- | --- | --- | --- |
| Host plants and<br>herbivorous insects | Plants (ca. 374,000) | Herbivorous insects (over<br>500,000) | Bernays (2009);<br>Christenhusz and Byng<br>(2016) |
| Host and parasites | Hosts (NA) | Parasites (ca. 6,000,000) | Dobson et al. (2008); |
| Host plants and<br>mutualistic mites | Some plants (NA) | Mutualistic mites (NA) | Romero and Benson<br>(2005) |
| Corals and *Symbiodinium* | Corals (ca. 10,661<br>estimated for Anthozoa) | *Symbiodinium* (NA) | Bánki et al. (2023) |
| Insects and gut microbes | Insects (ca. 5,500,000) | Gut microbes (NA) | Stork (2018) |

## METHODS

### Model description

#### Partnership between independent and dependent individuals

In the model, we consider the partnership between independent and dependent individuals. An independent individual can partner with multiple dependent individuals, but a dependent individual can only partner with one independent individual. An independent individual does not need to partner with any dependent individual for survival, but a dependent individual needs the one and only independent individual that it partners with for survival. This is modeled in two separate ways. First, during dispersal, an independent individual could survive where no dependent individuals are present; however, a dependent individual dies where no independent individuals are present. Second, during competition, when an independent individual dies, so do all of its dependent partners; however, when a dependent individual dies, its one and only independent partner does not die. It is important to note that, in nature, the party that interacts with multiple individuals are not necessarily the independent (less dependent) party.

#### Dispersal

The model starts with two populations each belonging to one of the interacting parties (the independent and dependent) placed at the central site of a 𝑛 × 𝑛 grid, where each cell represents a geographic site. Each individual, independent or dependent, has the opportunity to disperse only at birth and moves to one of the four neighboring sites or else remain at the same site (each with an equal probability of 0.2). As previously mentioned, if a dependent individual moves to a site where no independent individuals are present, it dies; however, if it moves to a site where at least one independent individual is present, the dependent individual randomly chooses an independent individual to partner with among the independent individuals available at that site. The entire grid forms a metacommunity composed of 𝑛 × 𝑛 communities (sites) connected through dispersal.

#### Sexual reproduction and speciation

The genetic model largely follows a previous model built for the diversification of a single clade (Aguilée et al. 2018), but with simplifications. Each independent or dependent individual is diploid and has a given number of *L_dist_* loci determining its genetic distance from another individual. A new mutation can arise at any of these loci at a fixed probability 𝜇_*dist*_. To determine whether two individuals are genetically incompatible enough to be considered reproductively isolated, we calculated the genetic distance between two individuals based on how many loci carry completely different alleles between the two individuals. The genetic distance is considered sufficient for reproductive isolation if greater than a fixed genetic distance threshold 𝑇_*dist*_. These features allowed speciation through genetic differentiation across the grid.

Both independent and dependent individuals are hermaphrodites and reproduce sexually. For independent individuals or dependent individuals, each mating attempt occurs within each site following these steps: (1) two individuals with the highest fitness are chosen regardless of whether they have reproduced before (see “Fitness outcomes of species interactions” for how fitness is decided; for simplicity, we did not choose a more complicated reproduction model); (2) the two parents successfully reproduce *n_offspr_* offspring if they are not reproductively isolated. These steps are repeated until *n_mat_* mating attempts are made, regardless of whether mating attempts result in successful reproduction. For each offspring, the genotypes of the genetic distance loci are determined by Mendelian independent assortment of parental alleles. Given that dependent individuals die along with their independent partner but not vice versa, it is necessary for *n_offspr_* and *n_mat_* to be greater for the dependent individuals than for the independent individuals so that the dependent individuals do not die out.

#### Genetics of phenotype

Each independent or dependent individual has one locus determining its ecological phenotype which consists of a single quantitative trait. Mutation occurs at each of these loci, with the mutated allele value drawn from a normal distribution with a mean equal to the parent allele value and a standard deviation equal to 𝜎_eco_. For each offspring, the genotype of the ecological phenotype locus is determined by a Mendelian random segregation of parental alleles.

#### Competitive death

Many types of antagonisms and mutualisms can occur between a consumer and a resource, such as herbivory, parasitism, or pollination (Bronstein 2015). Given that populations cannot grow infinitely in consumer-resource systems due to resource competition (Abrams 2009), we considered there to be a growth rate of zero when population size reached carrying capacity. In the model, resource competition occurs among dependent individuals partnering with the same independent individual (e.g., parasites on the same host) and among independent individuals within the same site. The number of mutually competing individuals 𝑛 cannot grow above carrying capacities *K_independent_* or *K_dependent_*. We ensured this by assigning *n_independent_* − *K_dependent_* or *n_dependent_* individuals to death (where *n_independent_* and *n_dependent_* are the numbers of mutually competing individuals, for independents and dependents, respectively), on a lowest-fitness-first basis (see “Fitness outcomes of species interactions” for how fitness is decided). Again, when an independent individual dies, so do all of the dependent individuals that partner with it; however, when a dependent individual dies, its one and only independent partner does not die.

#### Fitness outcomes of species interactions

We allowed different modes of phenotypic evolution to arise from species interactions with different fitness outcomes (Fig. 3). Antagonisms (interactions that benefit one party at the cost of the other) and mutualisms vary in whether each of the two interacting parties receives a benefit or pays a cost (Bronstein 2015). For either antagonisms or mutualisms, the fitness of phenotypes was modeled as the result of either (i) trait difference: how different the two interacting phenotypes were (the direction of phenotypic difference between interacting parties does matter), or (ii) trait matching: how closely two interacting trait values matched each other (the direction of phenotypic difference between interacting parties does not matter). Following Yoder and Nuismer (2010), the fitness of an individual 𝑖 interacting with an antagonist or mutualist individual 𝑗 can be expressed as

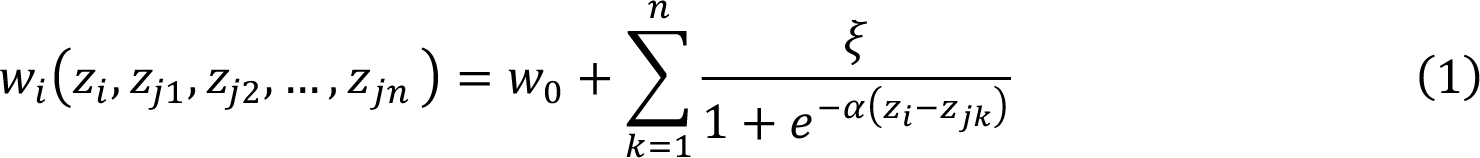

**Figure 3.**
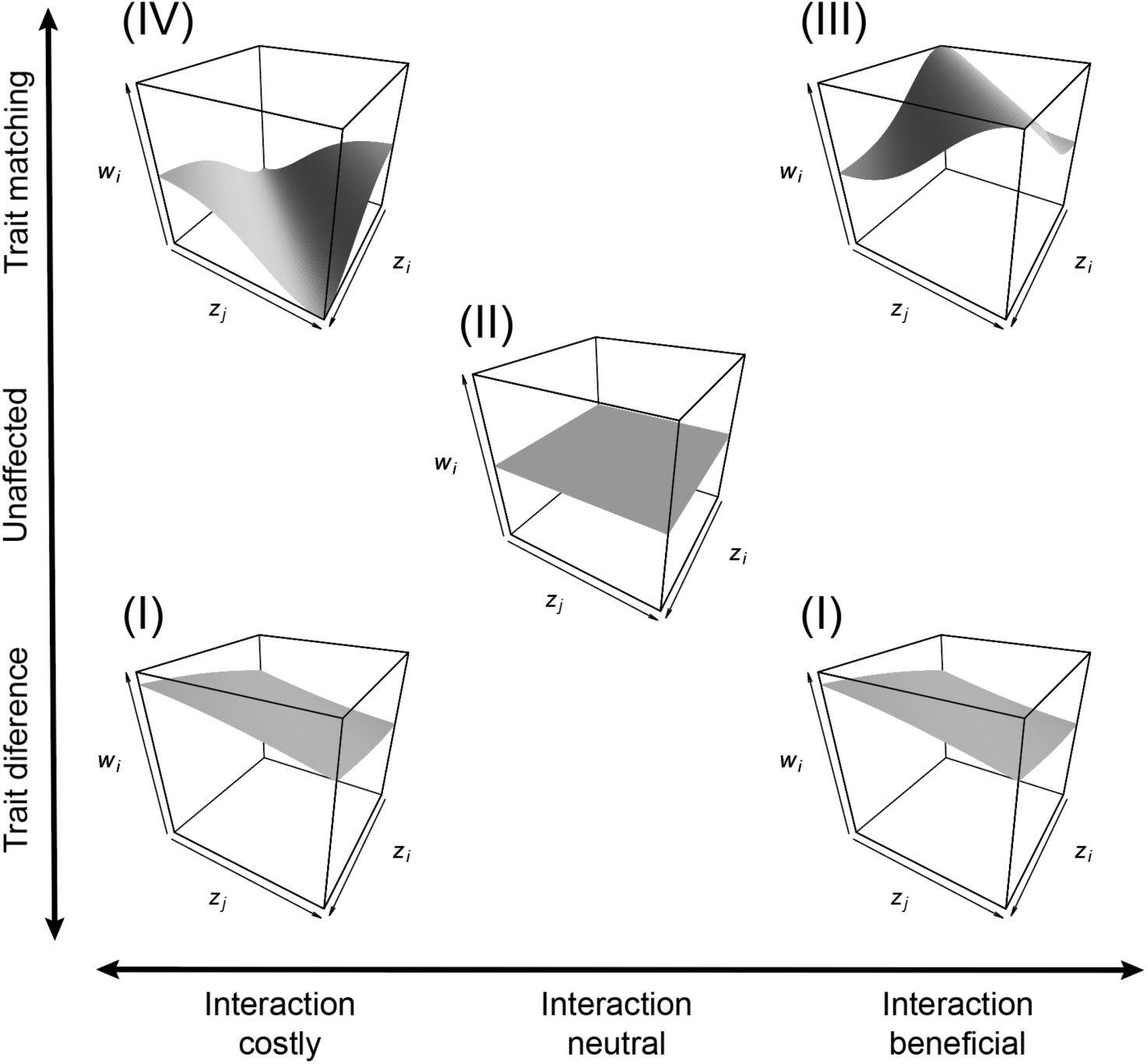
In species interactions, individual fitness is a function of the traits of interacting individuals. Each surface is the fitness of individual 𝑖 (𝑤_*i*_) as a function of 𝑧_*j*_, which denotes the trait value of individual 𝑖 itself, and 𝑧_*j*_, which denotes the trait value for individual 𝑗 (individual 𝑖’s interaction partner). Function I: Equation 1, 𝜉 = 0.1, 𝛼 = 1. Function II: 𝑤_*i*_ = 0. Function III: Equation 2, 𝜉 = 0.1, 𝛼 = 1. Function IV: Equation 2, 𝜉 = −0.1, 𝛼 = 1. For this model, Function II, where individual fitness is held constant and independent from trait values, is equivalent to Equation 1 or 2 when 𝛼 = 0 regardless of the value of 𝜉. Following Yoder and Nuismer (2010), the fitness function does not dependent on whether the interaction is beneficial or costly when coevolution operates through trait difference (Function I). In all panels, the ranges of the 𝑧_*i*_ and 𝑧_*j*_ axes are between 0 and 2 and the range of the 𝑤_*i*_ axis is between -0.1 and 0.1.

under trait difference, or

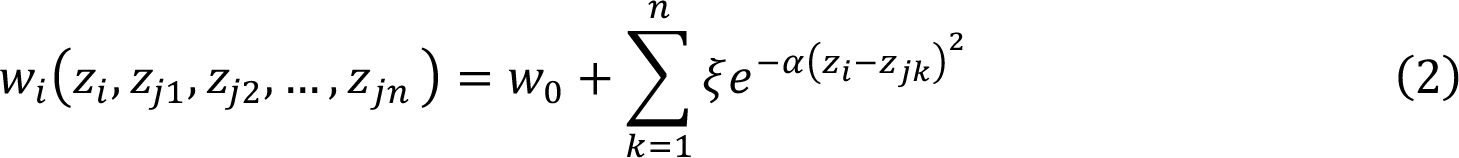

under trait matching, where 𝑧_*i*_ and 𝑧_*j*_ are phenotypic trait values of interacting individuals, 𝑛 is the number of partners that individual 𝑖 has, 𝑤_0_ is the individual 𝑖’s fitness in the absence of the interaction, 𝜉 is the cost or benefit to the individual (positive if beneficial, negative if costly), and 𝛼 determines the sensitivity of the fitness outcome to the difference between interacting trait values (i.e. deviation from fitness neutrality where trait values do not affect fitness). The strength of coevolutionary selection can often be highly asymmetrical, i.e., much weaker on one interacting party than on the other (Brodie and Brodie 1999; Andreazzi, Thompson, and Guimaraes 2017; Endara et al. 2017). Here we considered the extreme case of asymmetrical coevolutionary selection by allowing either the independents’ or dependents’ individual fitness to be unaffected by trait values (Function II in Fig. 3). A fully factorial design would contain 4*4=16 scenarios given the all the combinations of the 4 fitness functions. We simulated a total of 9 possible scenarios (Table 2) to include all possible combinations except 7 combinations of which no or few empirical cases are known, i.e., those where one clade’s fitness follows trait difference while the other’s follows trait matching (Yoder and Nuismer 2010) or where the interaction is costly for the dependent clade.

**Table 2.** The 9 scenarios considered in this study and their corresponding fitness functions (Figure 2). 4 fitness functions (I-IV) necessitate a total of 9 combinations after unrealistic combinations are excluded.

| Scenario | Trait difference or trait matching | Fitness function for the independent | Fitness function for the dependent | Outcome of interaction |
| --- | --- | --- | --- | --- |
| *a* | Difference | I | I | Mutualism or antagonism |
| *b* | Difference | I | II | Mutualism or antagonism |
| *c* | Difference | II | I | Mutualism or antagonism |
| *d* | Matching | III | III | Mutualism |
| *e* | Matching | III | II | Mutualism |
| *f* | Matching | II | III | Mutualism or antagonism |
| *g* | Matching | IV | III | Antagonism |
| *h* | Matching | IV | II | Antagonism |
| *i* | Unaffected | II | II | Mutualism or antagonism |

#### Simulations

The model was implemented and the simulation results were analyzed in the R language (v4.0.0; R Core Team, 2020). All simulations were run on the University of Arizona High-Performance Computing clusters. We ran 96 replicates for each of the 9 scenarios, totaling 864 simulations.

The large number of replicate simulations were made possible by custom code that can record the progress of a simulation and resume where that simulation is terminated by the workload manager. Each simulation was run for 1500 generations. For each of the 9 simulated scenarios, t-tests comparing the distributions of mean species richness during the 1401^th^-1450^th^ generations versus the 1451^th^-1500^th^ generations found no statistically significant difference (*P* > 0.05, *n* = 96 replicates for each of the two distributions being compared). This confirmed that the duration of simulation (1500 generations) was enough for simulations to reach a stationary state in terms of species richness, despite the ongoing fluctuations in species richness near the end of simulation (Fig. S1-S18). The large number of replicate simulations made the analyses computationally intense. This computational challenge was overcome though parallel computation using the R package doParallel (v1.0.17; Microsoft Corporation and Weston). All custom code used in this work is available as a supplementary file (File S1) and on GitHub (https://github.com/Dragonfly4412/Macro_Coevolution). All the constants used in the simulations are provided in Table S1.

#### Identifying selective regimes based on temporal trait (in)stability

To identify the modes of phenotypic evolution for each scenario, we recorded the phenotypic values of all independent and dependent individuals at the central site of the 𝑛 × 𝑛 grid during the entire duration of each simulation (1500 generations). We then quantified the change in mean trait value during each generation, Δ𝑧. Δ𝑧 is conventionally referred to as step difference and is measured in haldanes, i.e., standard deviations per generation on a timescale of one generation (Gingerich 2019a; Gingerich 2019b). For example, a Δ𝑧 value of 2 haldanes means that the change in mean trait value from Generation 1 to Generation 2 is two times the standard deviation of the trait distribution at Generation 1. We further took the absolute value of Δ𝑧 to get |Δ𝑧|. We then averaged |Δ𝑧| across the entire duration of simulation to get the mean step difference |Δ𝑧̅| as a measure of temporal trait instability. We determined the minimum and maximum for the |Δ𝑧̅| of the entirely neutral scenario (Scenario *i*, where both the independents’ and dependents’ fitness are held constant and do not depend on trait values). We treated simulations in which |Δ𝑧̅| is greater than the neutral maximum as destabilizing selection and those in which |Δ𝑧̅| is less than the neutral minimum as stabilizing selection.

#### Degree of dispersal limitation, genetic distance between sites, and species richness

We quantified the degree of dispersal limitation as the proportion of native individuals among all individuals, with higher proportions of native individuals indicating stronger degrees of dispersal limitation. We define a native individual as an individual that inhabits the same site at which it was born. Specifically, what we measured was realized dispersal rather than dispersal *per se*, because we are interested in the contribution of dispersal to local gene pools, which is contingent on successful establishment of the dispersers following dispersal. The degree of dispersal limitation we refer to is the degree to which dispersal success is limited. Then, we calculated genetic distance between any two sites (i.e., between the two populations inhabiting the two sites) by taking the mean genetic distance between any two individuals from the two sites (i.e., the two populations). The genetic distance between all sites was then quantified as the mean genetic distance between any two sites. To quantify the species richness that each clade had accumulated at the end of the simulation, we considered populations to belong to a single species if their genetic distance did not exceed the genetic distance threshold for speciation *T_dist_* (for details, see “Sexual reproduction and speciation”).

For the degree of dispersal limitation (the proportion of native individuals), we took the mean over the entire 1500 generations of simulation because it is the cumulative effect of dispersal that is of interest. For genetic distance between sites and species richness accumulation, we took the mean over the last 10 generations of simulation because we were interested in them as the eventual results of multiple generations of dispersal and speciation.

## RESULTS

Each of the 9 simulated scenarios generated a unique coevolutionary trajectory within communities (Fig. 4). The 9 result coevolutionary trajectories were each characterized by rapid and continuous reciprocal escalation of trait values in the two clades characteristic of an arms race (Scenario *a*; Fig. 4), trait escalation in the independent clade and an initial decrease in trait value followed by a random walk (in the sense that it does not substantially differ from a trajectory typical of the neutral scenario) in the dependent clade (Scenario *b*; Fig. 4), trait escalation followed by a random walk in the dependent clade and random walk in the independent clade (Scenario *c*; Fig. 4), varying degrees of temporal trait stationarity in the two clades (Scenarios *d*-*f*; Fig. 4), dramatic fluctuation in trait value in both clades characteristic of coevolutionary cycling (Scenario *g*; Fig. 4), a tendency for the host trait value to not overlap with that of the dependent (Scenario *h*; Fig. 4), and random walks underlain by neutral dynamics in both clades (Scenario *i*; Fig. 4).

**Figure 4.**
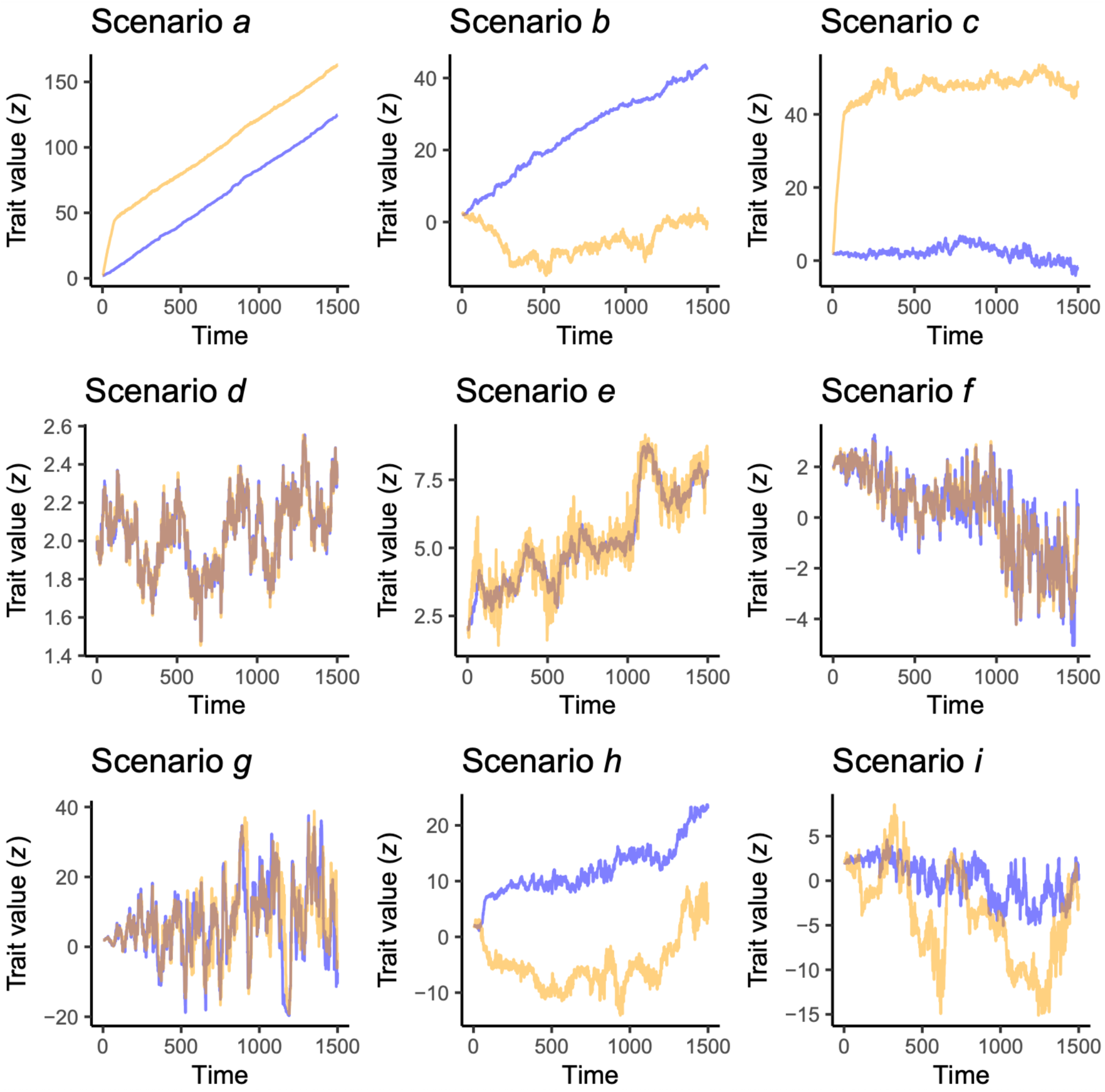
Coevolutionary trajectories within communities in each of the nine simulated scenarios (*a*-*i*). Solid lines indicate the mean trait value calculated for individuals at the central community (site) of the 𝑛 × 𝑛 grid in one of the 96 replicates for each scenario. Blue and yellow are used to indicate the independent and dependent trait values, respectively.

Analyses of the effects of selective regime on the degree of dispersal limitation, genetic distance between sites, and species richness accumulation revealed the mechanisms by which coevolution shaped species richness (Fig. 5). Quantification of the mean step difference |Δ𝑧| as a measure of temporal trait instability showed that the 9 scenarios generated all three modes of phenotypic evolution, i.e. neutral drift, stabilizing selection, and destabilizing selection in the two clades. For each of the relationships of interest (Fig. 5, A-C), we performed a loess regression to visualize non-linearities, but also two localized simple linear regressions to quantify the average effects of stabilizing and destabilizing selection (Fig. S19).

**Figure 5.**
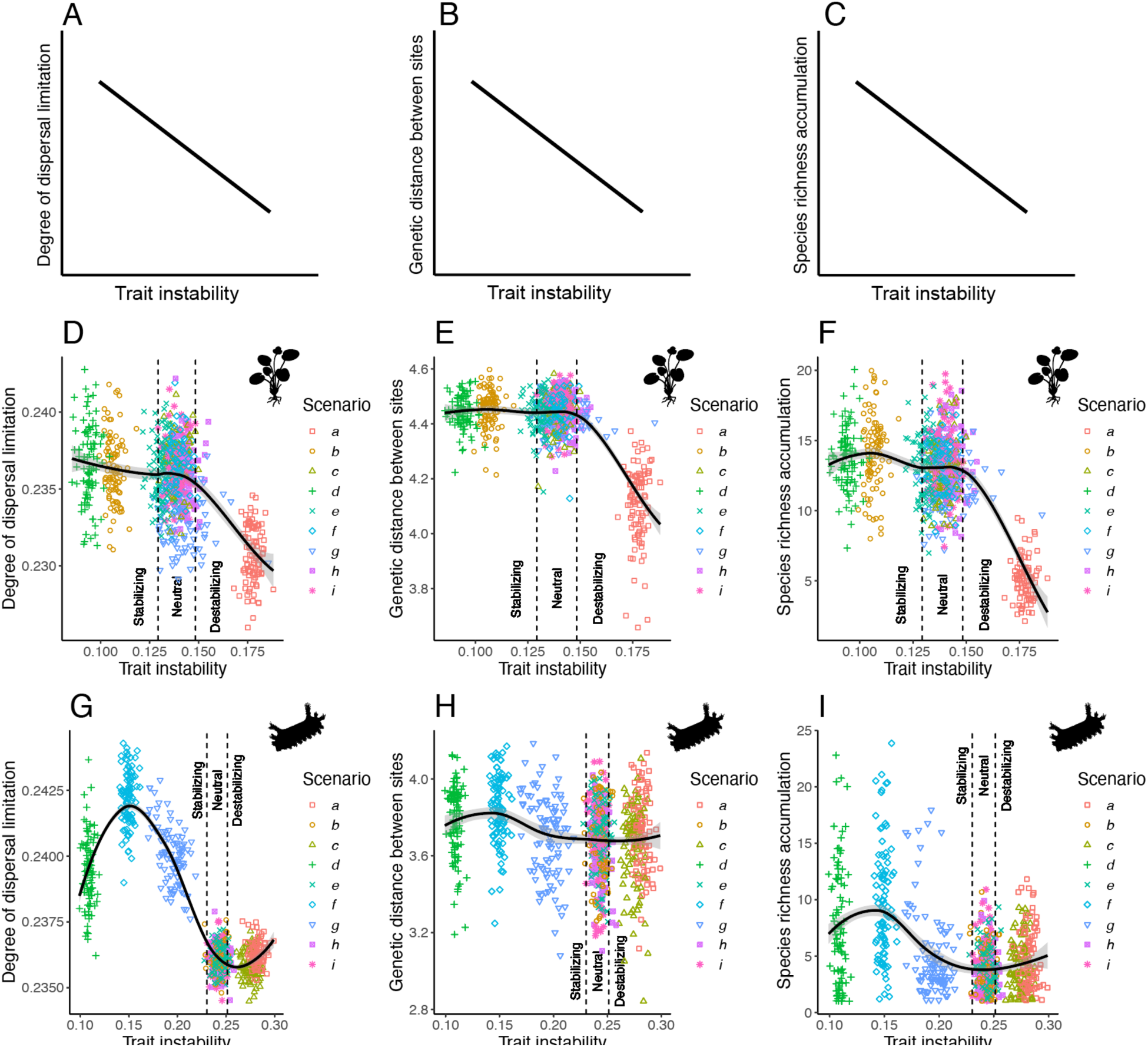
Observed relationships between with-community dynamics (trait instability), between-community dynamics (degree of dispersal limitation), and metacommunity dynamics (genetic distance between sites and species richness accumulation). Selective regimes (neutral drift, stabilizing selection, or destabilizing selection) are determined by trait instability measured as the mean step difference ^1^|Δ^11^𝑧^11^|. (A)-(C): Expected relationships – a significantly negative slope is predicted based on the stabilizing and destabilizing selection hypotheses. (D)-(F): Observed differences between selection regimes in the independent clade. (G)-(I): Observed differences between selection regimes in the dependent clade. In (D)-(I), black lines with gray ribbons show loess regressions with 95% confidence intervals.

In the independent clade (Fig. 5, D-F; Fig. S19, D-F), compared to neutral drift, stabilizing selection resulted in relatively weak increases in the degree of dispersal limitation (slope = -1.47 percentage/haldane, *P* = 0.0013, left line in Fig. S19D), genetic distance between sites (slope = - 0.1664 loci/haldane, *P* = 0.208, left line in Fig. S19E), and species richness accumulation (slope = -16.4918 species/haldane, *P* = 0.00117, left line in Fig. S19F). However, compared to neutral drift, destabilizing selection resulted in relatively strong reductions in degree of dispersal limitation (slope = -12.11 percentage/haldane, *P* < 2e-16, right line in Fig. S19D), genetic distance between sites (slope = -7.60415 loci/haldane, *P* < 2e-16, right line in Fig. S19E), and species richness accumulation (slope = -181.4629 species/haldane, *P* < 2e-16, right line in Fig. S19F). These results suggest that the predominant mechanism shaping species richness in the independent clade is one where destabilizing selection reduces the degree of dispersal limitation, which reduces genetic distance between sites, eventually resulting in reduced species richness accumulation.

In the dependent clade (Fig. 5, G-I; Fig. S19, G-I), compared to neutral drift, stabilizing selection resulted in relatively strong increases in the degree of dispersal limitation (slope = -3.755 percentage/haldane, *P* < 2e-16, left line in Fig. S19G), genetic distance between sites (slope = - 1.0355 loci/haldane, *P* = 2.87e-12, left line in Fig. S19H), and species richness accumulation (slope = -35.7923 species/haldane, *P* < 2e-16, left line in Fig. S19I). The effect of stabilizing selection showed strong non-linearity, with the highest degree of dispersal limitation, genetic distance between sites, and species richness accumulation being achieved by intermediately rather than extremely strong stabilizing selection (Fig. 5, G-I). However, compared to neutral drift, destabilizing selection resulted in relatively week reductions or even a slight increase in the degree of dispersal limitation (slope = -0.00114 percentage/haldane, *P* = 0.993, right line in Fig. S19G), genetic distance between sites (slope = -0.08698 loci/haldane, *P* = 0.853, right line in Fig. S19H), and species richness accumulation (slope = 6.4800 species/haldane, *P* = 0.375, right line in Fig. S19I). These results suggest that the predominant mechanism shaping species richness in the dependent clade is one where stabilizing selection increased the degree of dispersal limitation, which increases genetic distance between sites, eventually resulting in increased species richness accumulation.

Specifically, the destabilizing selection hypothesis explained how species richness accumulation was reduced for the independent clade in the coevolutionary scenarios of mutualistic or antagonistic arms races where selective pressure is comparable between the independent and dependent individuals, i.e., a classical arms race, and in the scenario of antagonistic trait matching where selective pressure is comparable between the independent and dependent individuals (Fig. 5, D-F; Fig. S19, D-F; Scenarios *a* & *g* in Table 2). The variable |Δ𝑧|, as a measure of temporal trait instability, is turned out to be key in shaping species richness accumulation. This is because trait instability is a direct result of coevolution and in turn shapes dispersal and consequently speciation. The stabilizing selection hypothesis explained how species richness was increased for the dependent clade in the coevolutionary scenario of mutualistic trait matching where selective pressure is comparable between the independent and dependent individuals, the scenario of mutualistic or antagonistic trait matching where selective pressure is weak for independent individuals, and the scenario of antagonistic trait matching where selective pressure is comparable between the independent and dependent individuals (Fig. 5, G-I; Fig. S19, G-I; Scenarios *d*, *f* & *g* in Table 2).

## DISCUSSION

Here we have shown that coevolution shapes species richness through two different mechanisms depending on whether the clade of interest is independent or dependent. In the independent clade, destabilizing selection enhances dispersal, which results in a reduction in the genetic distance across space, eventually resulting in a reduction in the species richness accumulation (i.e. the destabilizing selection hypothesis). In the dependent clade, stabilizing selection impedes dispersal, which results in an increase in the genetic distance cross space, eventually resulting in an increase in the species richness accumulation (i.e. the stabilizing selection hypothesis). These results arose through two simple rules governing the dynamics of independent-dependent interactions: one independent individual can pair with multiple dependent individuals but not vice versa; a dependent individual dies in the absence of an independent partner but not vice versa. These rules should be applicable to a wide variety of systems including most host-phytophagous-insect interactions, most host-parasite interactions, most host-pathogen interactions, and some obligate interactions between mutualists and their hosts, although exceptions inevitably exist because the biology of species interactions is highly diverse..

Empirical data generally corroborate our results that coevolution-induced stabilizing and destabilizing selection differentially affect the dispersal of independent and dependent individuals, suggesting that our model provides a potential mechanistic explanation for these patterns of dispersal. Previous models have explored the conditions for coevolutionary stable states or strategies to be achieved, yet these model did not consider dispersal or speciation across space (Vasco, Nazarea & Richardson 1987; Gilchrist & Sasaki 2002; Day & Burns 2003).

Empirical examples of coevolution-induced destabilizing selection include classical arms race where there is reciprocal escalation of trait values (Dawkins & Krebs 1979; Vermeij 1994; Whittall & Hodges 2007) and coevolutionary cycling where there is persistent alternation between different trait states such between high and low trait values (Prado et al. 2009; Ashby & Gupta 2014; Ashby & Boots 2015). Whether coevolution causes stabilizing or destabilizing selection is usually unknown except in a limited number of empirical systems, but the results of our model tend to match several empirical observations pertaining to ant-plant, ant-bacterium, and plant-fungus mutualisms. First, our model shows that coevolution-induced stabilizing selection impedes the dispersal of dependent individuals only. Although this conclusion is based upon correlation, the direction of causation is inambiguous because the model took a bottom-up approach where dispersal, a between-community process, arose as a result of species interaction and coevlution within communities. This matches large-scale observations that only specialized mutualism (high dependence on mutualistic partners) is associated with a reduced likelihood of successful establishment beyond native ranges (Nathan et al. 2023). Second, our model shows that coevolution-induced destabilizing selection enhances the dispersal of independent individuals only. This matches large-scale observations that only generalized mutualism (low dependence on mutualistic partners) is associated with an increased likelihood of successful establishment beyond native ranges (Nathan et al. 2023). We show that coevolution-induced stabilizing and destabilizing selection can generate these widely observable patterns. This suggests that coevolution-induced stabilizing and destabilizing selection, as well as their macroevolutionary consequences as shown in our model, may be prevalent in nature.

It is useful to understand these results in the light of environmental selection and trait multi-dimensionality. Some traits under coevolutionary selection may be simultaneously under selection from other factors in the environment. Selection imposed by the environment can be either stabilizing or destabilizing – environment-induced stabilizing selection can operate through selection against phenotypes that deviate from a fixed environmental optimum (Kopp & Matuszewski 2014), whereas fluctuating environment can cause potentially destabilizing selection (De Villemereuil 2020). Disruptive selection along an environmental gradient has itself been shown as a mechanism driving speciation (Doebeli & Dieckmann 2003). It would be interesting for future research to explore the interactive effects of coevolutionary selection and environmental selection on co-diversification, as has been pointed out in cases where biotic interactions drive speciation in tandem with spatial isolation (Kay and Sargent 2009, Hembry et al. 2014). In addition, species interactions are sometime better modeled using multi-dimensional traits (Eklöf et al. 2013). The trait difference mechanism can have similar effects on phenotypic evolution to those of the trait matching mechanism when the trait of interest is multidimensional (Yamamichi, Lyberger & Patel 2019). Therefore, destabilizing selection underlain by the trait difference mechanism might be less common in systems with higher trait dimensionalities.

These results may provide mechanistic insights into the diversification of some extremely diverse clades that fit the description of a dependent clade. Specifically, explaining the origins of the diversity of phytophagous insects and parasites has long been an active area of research (Ehrlich and Raven 1964; Poulin and Morand 2000; Hardy and Otto 2014; Weinstein and Kuris 2016; Kawahara et al. 2023), and our model provided a potential explanation for their staggering diversity: stabilizing selection induced by coevolution selects against immigrants, reducing dispersal success and consequently enhancing genetic differentiation across space that is necessary for speciation under our assumption. This is conceptually similar to the notion that local coadaptation can reduce the chance of mating between populations and this reduction in gene flow can promote reproductive isolation and consequently speciation (Thompson, Segraves & Althoff 2017). However, we furthered this notion by showing that this is likely to be the case only for clades that fit our assumption about dependence such as phytophagous insects, parasites, pathogens, or obligate mutualists that depend on facultative hosts, but is unlikely to be the case for clades that fit our assumption about independence such as various clades of hosts as mutualists or victims of phytophagous insects, parasites, and pathogens. Overall, the mechanisms from this model provide potential mechanisms by which coevolution may have had a profound impact on the diversification of phytophagous insects and parasites, although the prevalence of coevolution in insect herbivory and parasitism has yet to be confirmed by empirical evidence.

Our results suggest that the diversity of clades that fit our description of an independent clade (e.g., plants, hosts of parasites and pathogens, or the facultative hosts of obligate mutualists) could also be explained in the light of coevolution. The effect of coevolution on an independent clade’s diversity, under our model assumptions, is likely negative for diversification as shown in the destabilizing selection hypothesis. It is interesting in this light that independent clades, e.g., clades of angiosperms are often more species-poor than their dependent clades, e.g., clades of phytophagous insects (Bernays 2009; Christenhusz & Byng 2016), although many other mechanisms likely contribute to this asymmetry in richness. This decelerating effect of coevolution on diversification that we show here has not been explored before, as classic coevolutionary hypotheses such as the escape-and-radiate hypothesis tended to focus on the accelerating effects of coevolution on diversification (Ehrlich and Raven 1964). The escape-and-radiate hypothesis is less interested in speciation events per se than in bursts of diversification in lineages that escape herbivory and is not specifically accounted for in our model.

Comparing our model to previous models provides some additional implications. We show that a classical arms race impedes the accumulation of species richness in the independent clade but has no effect in the dependent clade. Our results focusing on species richness is in contrast with the results of a previous study showing that the classical arms race neither promotes nor inhibits phenotypic diversification (Yoder and Nuismer 2010). We also show that destabilizing selection also predicts a decrease in species richness in the independent clade in the scenario of antagonistic trait matching where selective pressure is comparable between the independent and dependent individuals. This also contrasts with the previous model which showed that coevolution promotes phenotypic diversification when trait matching is costly, e.g., as in competition or antagonisms. These contrasts suggest that phenotypic diversification can be decoupled from species diversification during coevolution. The idea that having multiple partners poses a constraint on diversification when the interaction is mutualistic and based on trait matching has been shown in a non-spatially explicit model previously (Raimundo et al. 2014). This agrees with the results of our spatially explicit model and suggests that our results could potentially arise through similar mechanisms. Overall, the new model contributes to an ongoing effort to integrate metacommunity ecology with macroevolution in general (McPeek 2008; Reijenga, Murrell & Pigot 2021) and for independent-dependent systems in particular (Forister & Jenkins 2017).

Our model provides an alternative or supportive explanation for some long-standing hypotheses. In the seminal escape-and-radiate hypothesis (Ehrlich & Raven 1964), a lineage, independent or dependent, that acquires a new defense or counter-defense (i.e. key innovations) may then rapidly radiate into a new adaptive zone. We show a different picture here – coevolutionary spatial dynamics can continuously enhance or impede speciation without involving sporadic key innovations. There are also long-standing hypotheses that dependents should be more specialized the more intimate their interactions with their independents are (Ollerton 2006; Thompson 1994). These hypotheses generally predict that dependents should be more species-rich than independents, which is consistent with our results suggesting that coevolution generally impedes diversification for an independent clade and enhances diversification for a dependent clade. However, the intimacy hypotheses are agnostic to the geography of speciation. It is also clear that a single independent can contain a multitude of niches for different dependents (Farrell & Sequeira 2004), so the asymmetry in richness is less surprising regardless of the specific mechanisms of diversification.

In conclusion, here we show that there are two general mechanisms of coevolutionary diversification: coevolution-induced stabilizing selection enhances the accumulation of species richness in dependent clades, whereas coevolution-induced destabilizing selection impedes the accumulation of species richness in independent clades. The model provides a new line of thinking in bridging symbiotic biology, coevolution, metacommunity ecology, and macroevolution. Given that symbiotic relationships between a more dependent party and a less dependent party permeate the natural world (Margulis 1998), these general mechanisms of coevolutionary diversification can potentially explain the diversification of many clades in the Tree of Life.

## Authorship statement

**Yichao Zeng**: Conceptualization, Methodology, Software, Validation, Formal analysis, Investigation, Data cutation, Writing – Original draft, Review & Editing, Visualization. **David H. Hembry**: Writing – Original draft, Review & Editing.

## Acknowledgements

We thank Judie Bronstein, Régis Ferrière, Paulo R. Guimarães Jr., and John Wiens for discussing and reviewing early versions of this manuscript. John Wiens edited early versions of this manuscript. High-performance computing resources were provided by the University of Arizona. These high-performance computing resources were funded by central funding and were free to use for all university affiliates (https://public.confluence.arizona.edu/display/UAHPC/Compute+ Resources). We thank Sara Willis for technical support with cluster computing.

