## Supplementary material for "Coevolution-induced stabilizing and destabilizing selection shapes species richness in clade co-diversification": Figures S1-S19

Table of contents

Figures S1-S9 – Change in independent species richness over time, for Scenarios *a*-*i*

Figures S10-S18 – Change in dependent species richness over time, for Scenarios *a*-*i*

Figure S19 – The linear regressions of the relationships between selective regime, degree of dispersal limitation, genetic distance among sites, and species richness accumulated.

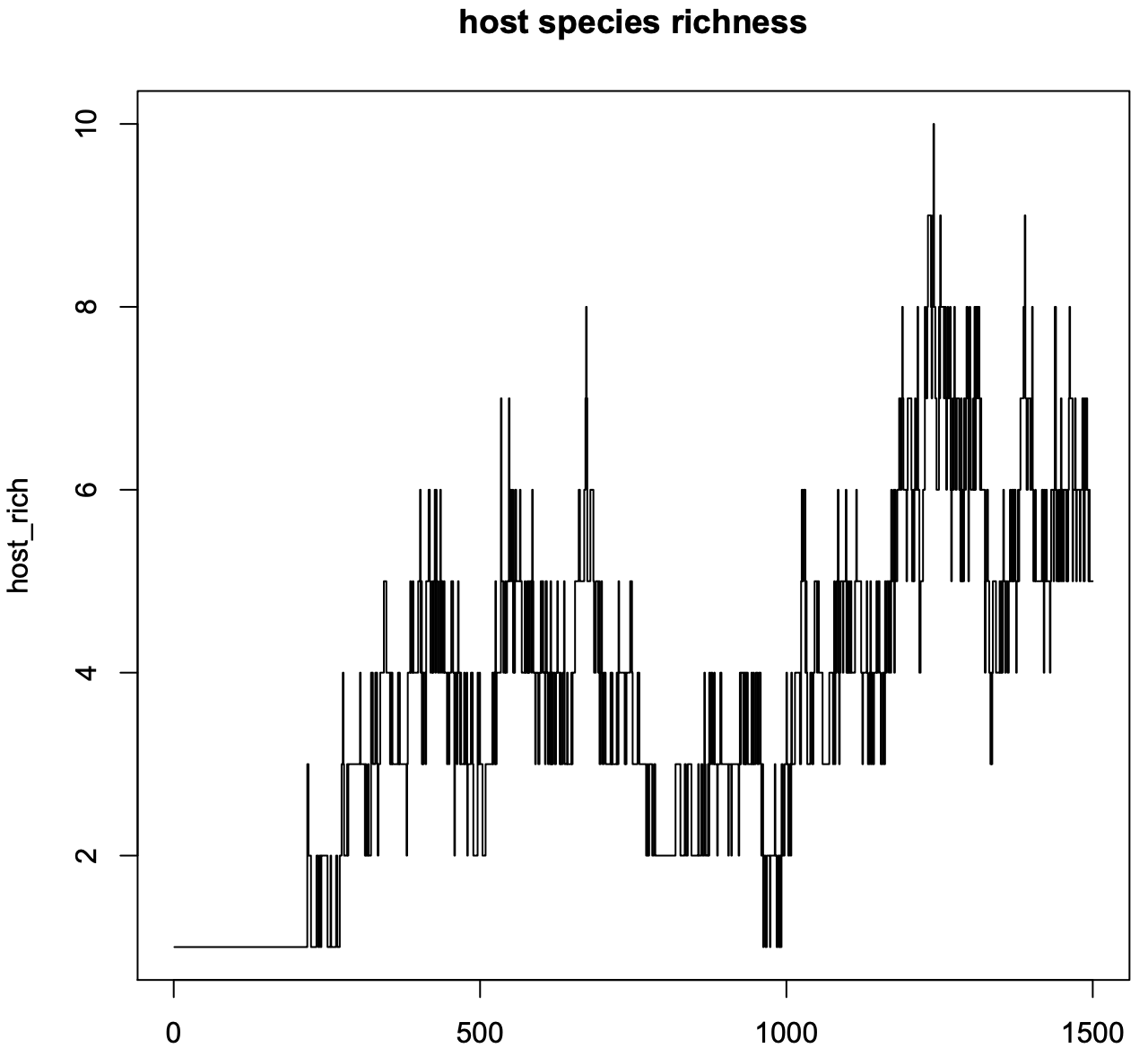

Figure S1. Change in independent species richness over time, for Scenarios *a*. Shown here is one of the 96 replicates that were run for this scenario.

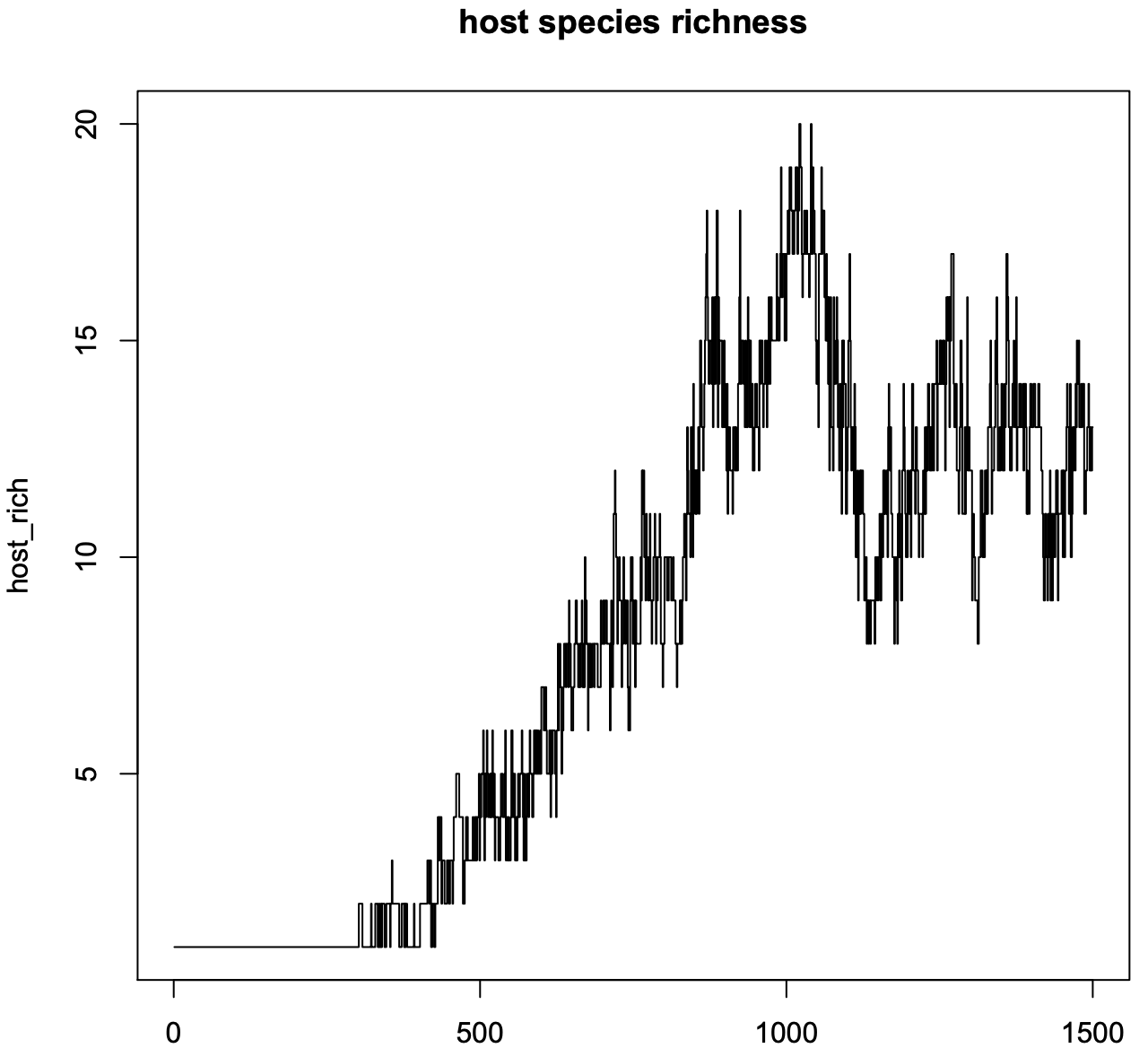
Figure S2. Change in independent species richness over time, for Scenarios *b*. Shown here is one of the 96 replicates that were run for this scenario.

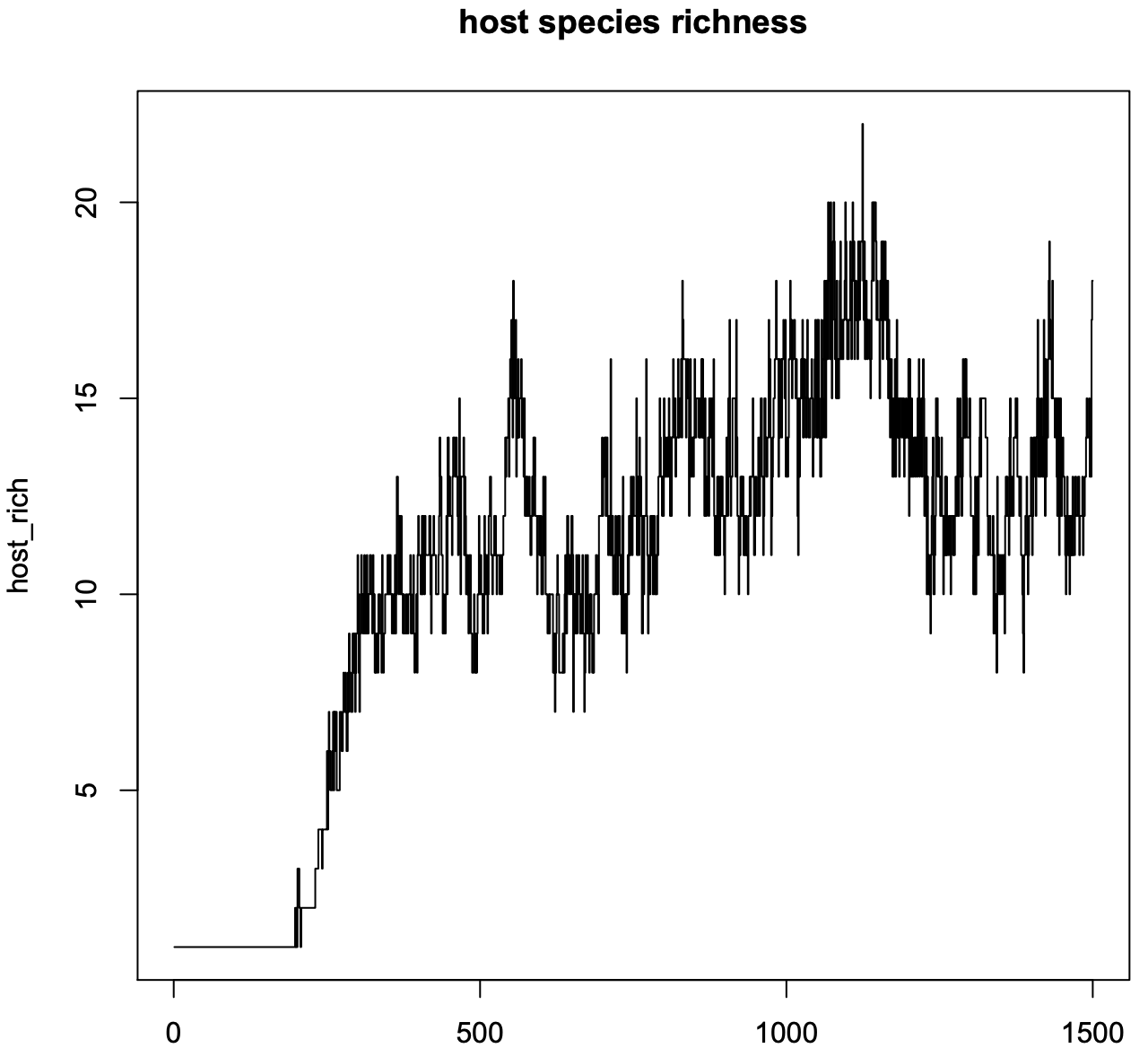
Figure S3. Change in independent species richness over time, for Scenarios *c*. Shown here is one of the 96 replicates that were run for this scenario.

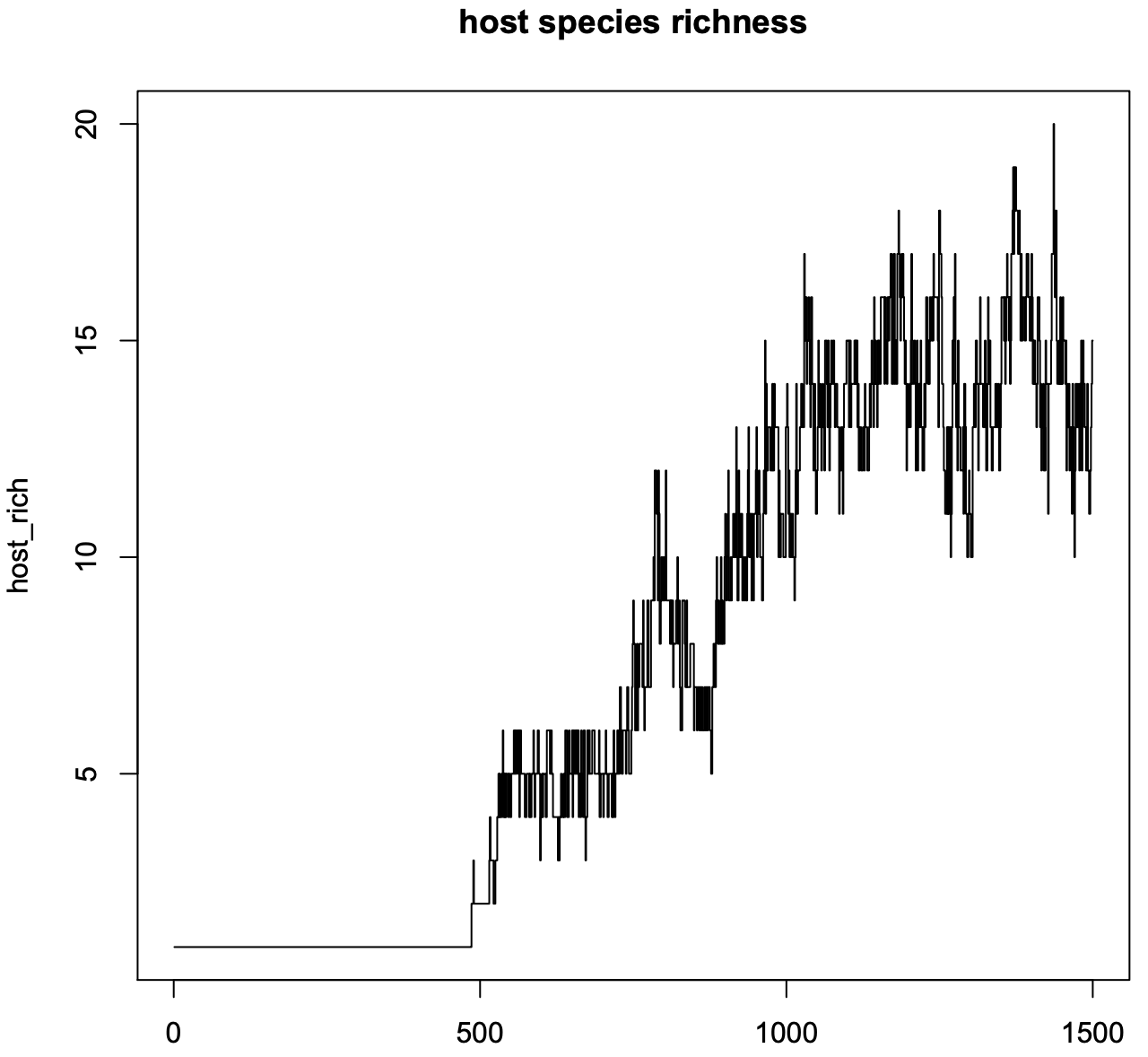
Figure S4. Change in independent species richness over time, for Scenarios *d*. Shown here is one of the 96 replicates that were run for this scenario.

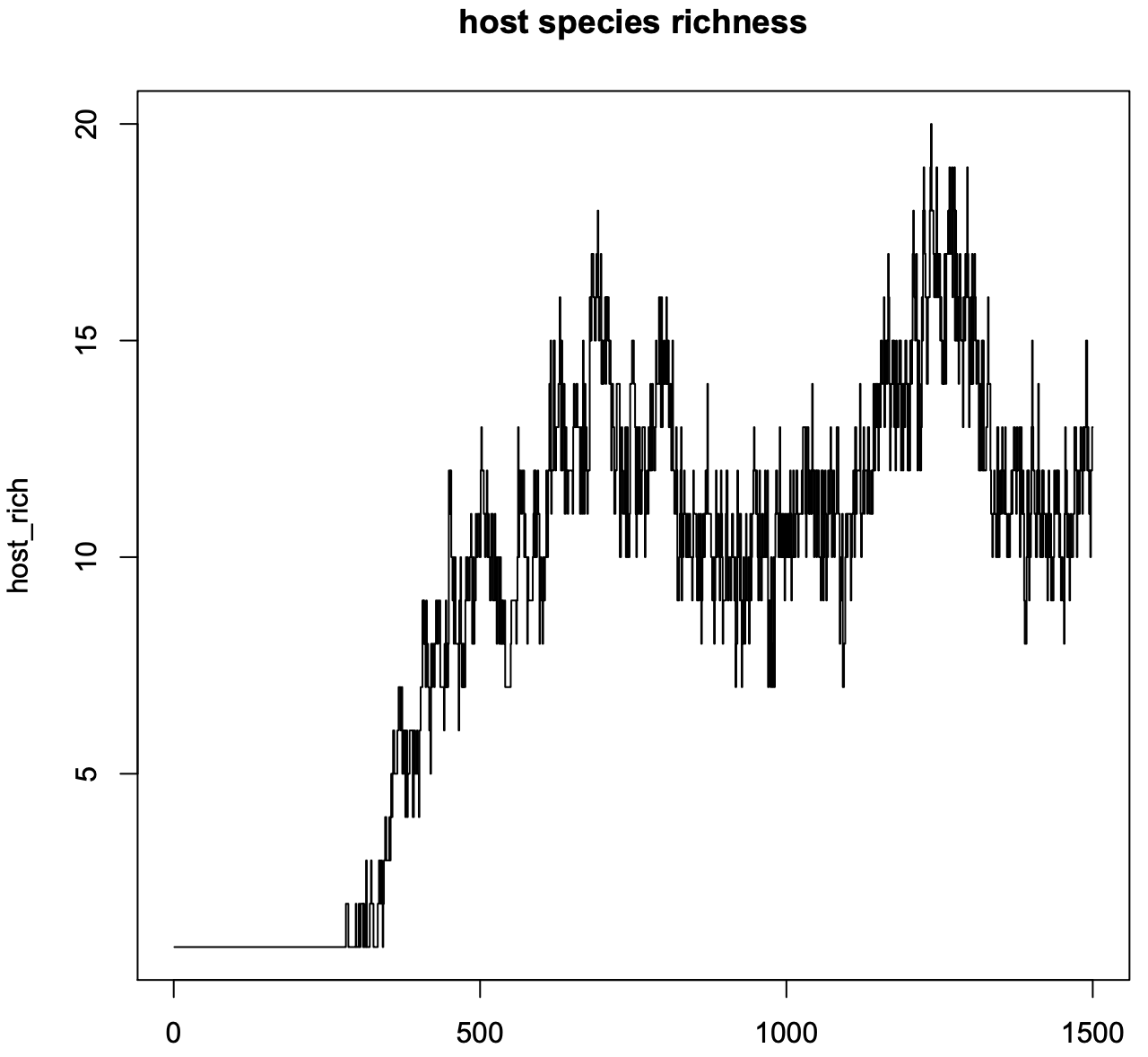
Figure S5. Change in independent species richness over time, for Scenarios *e*. Shown here is one of the 96 replicates that were run for this scenario.

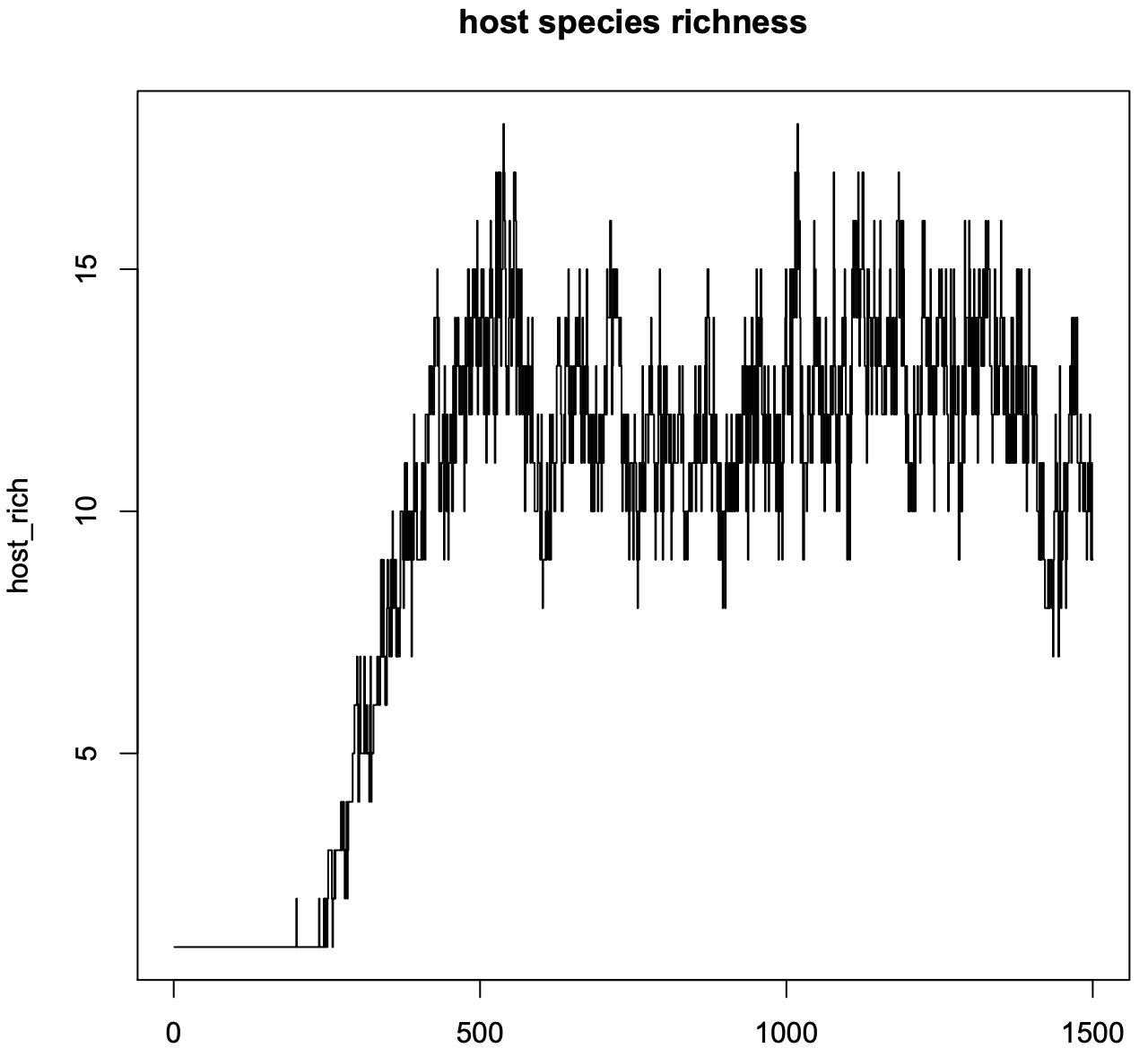
Figure S6. Change in independent species richness over time, for Scenarios *f*. Shown here is one of the 96 replicates that were run for this scenario.

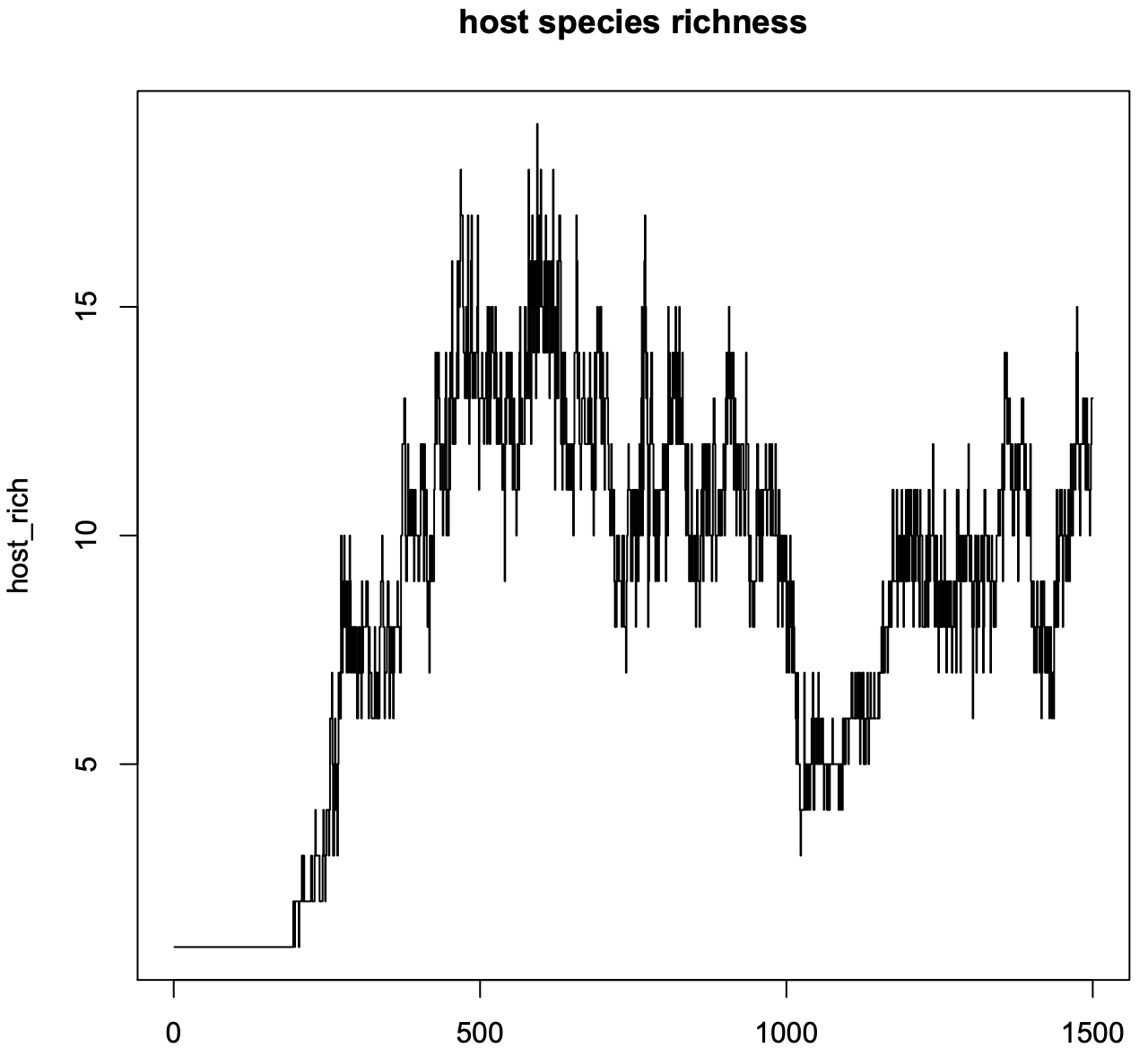
Figure S7. Change in independent species richness over time, for Scenarios *g*. Shown here is one of the 96 replicates that were run for this scenario.

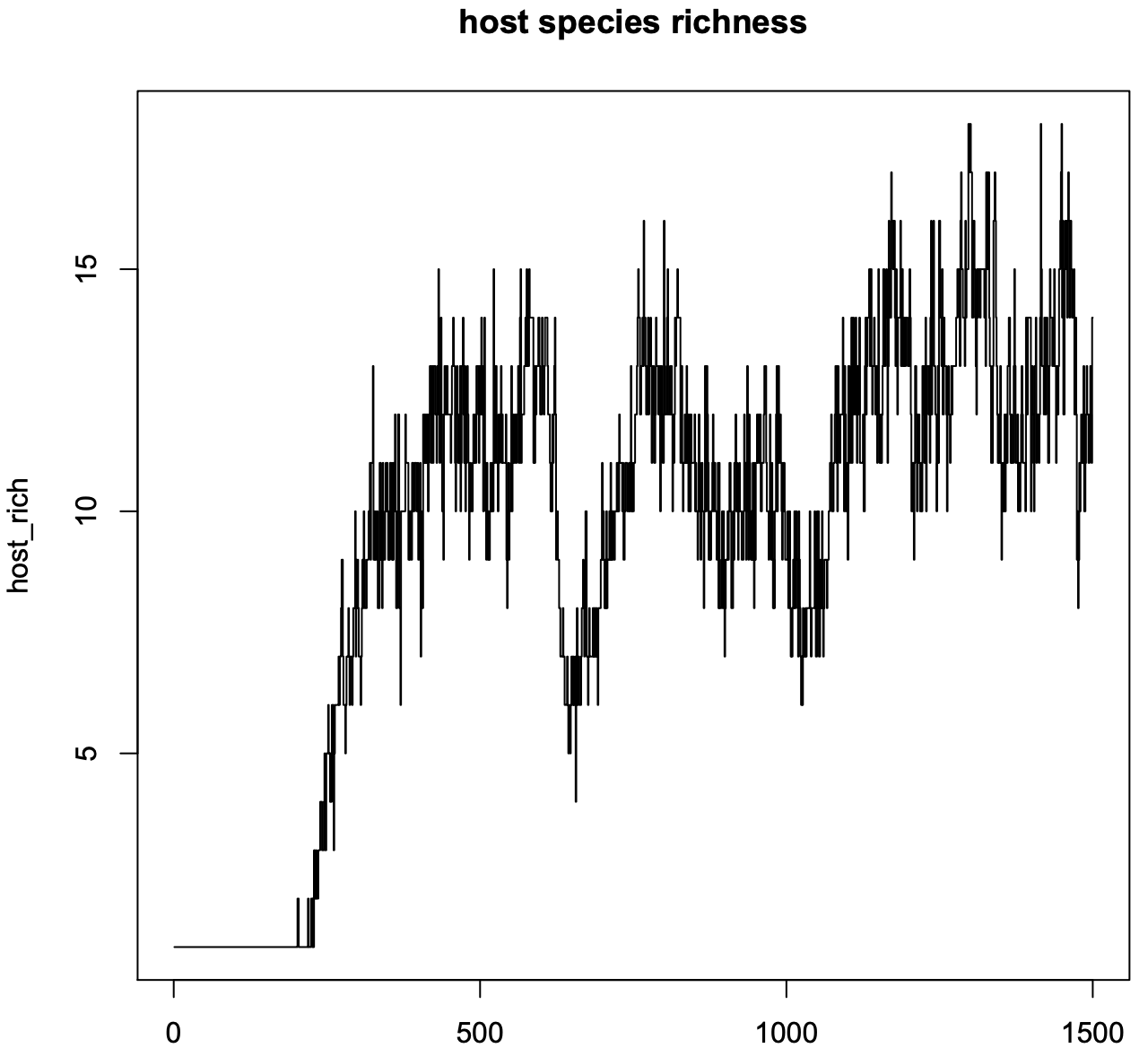
Figure S8. Change in independent species richness over time, for Scenarios *h*. Shown here is one of the 96 replicates that were run for this scenario.

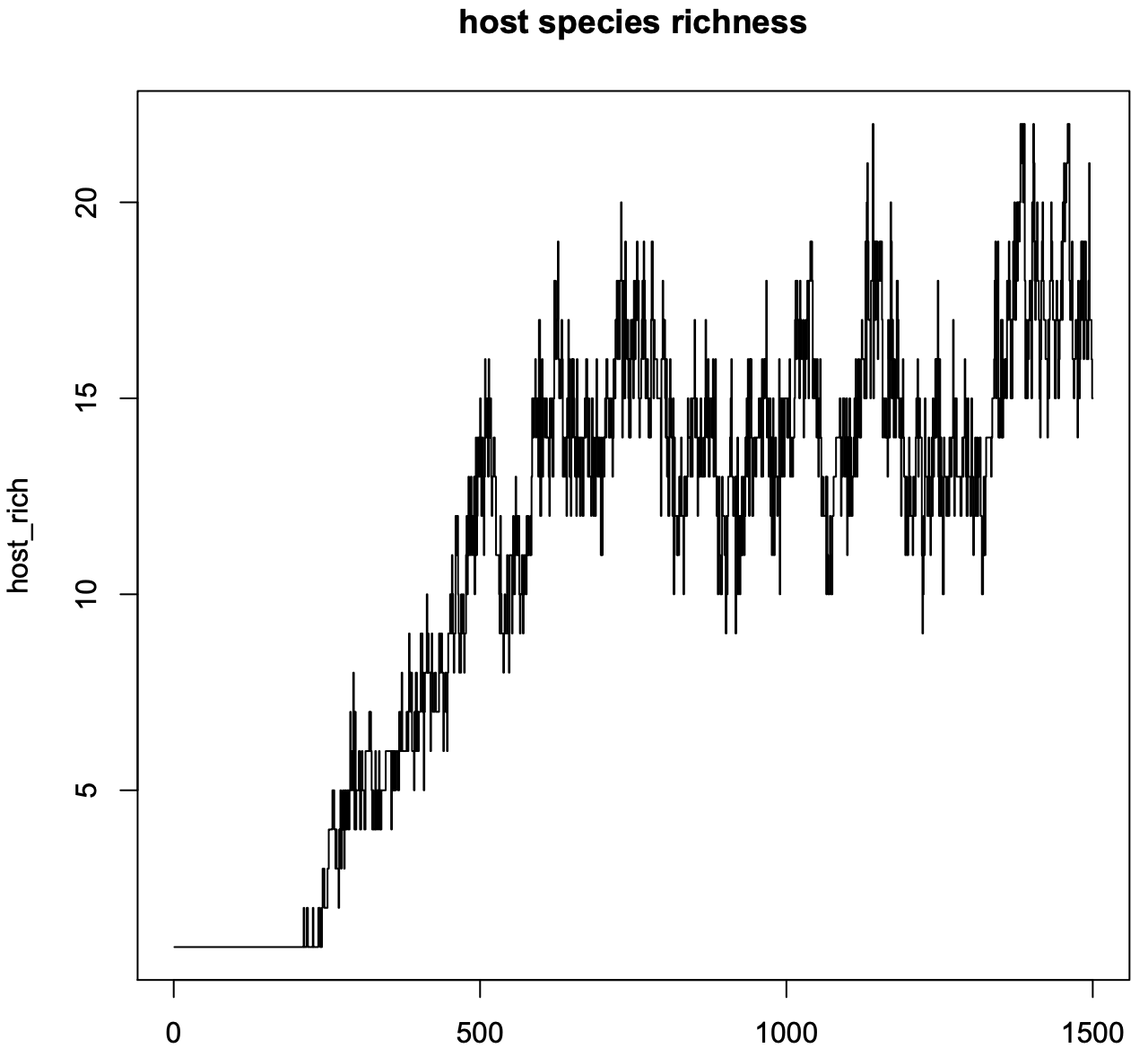
Figure S9. Change in independent species richness over time, for Scenarios *i*. Shown here is one of the 96 replicates that were run for this scenario.

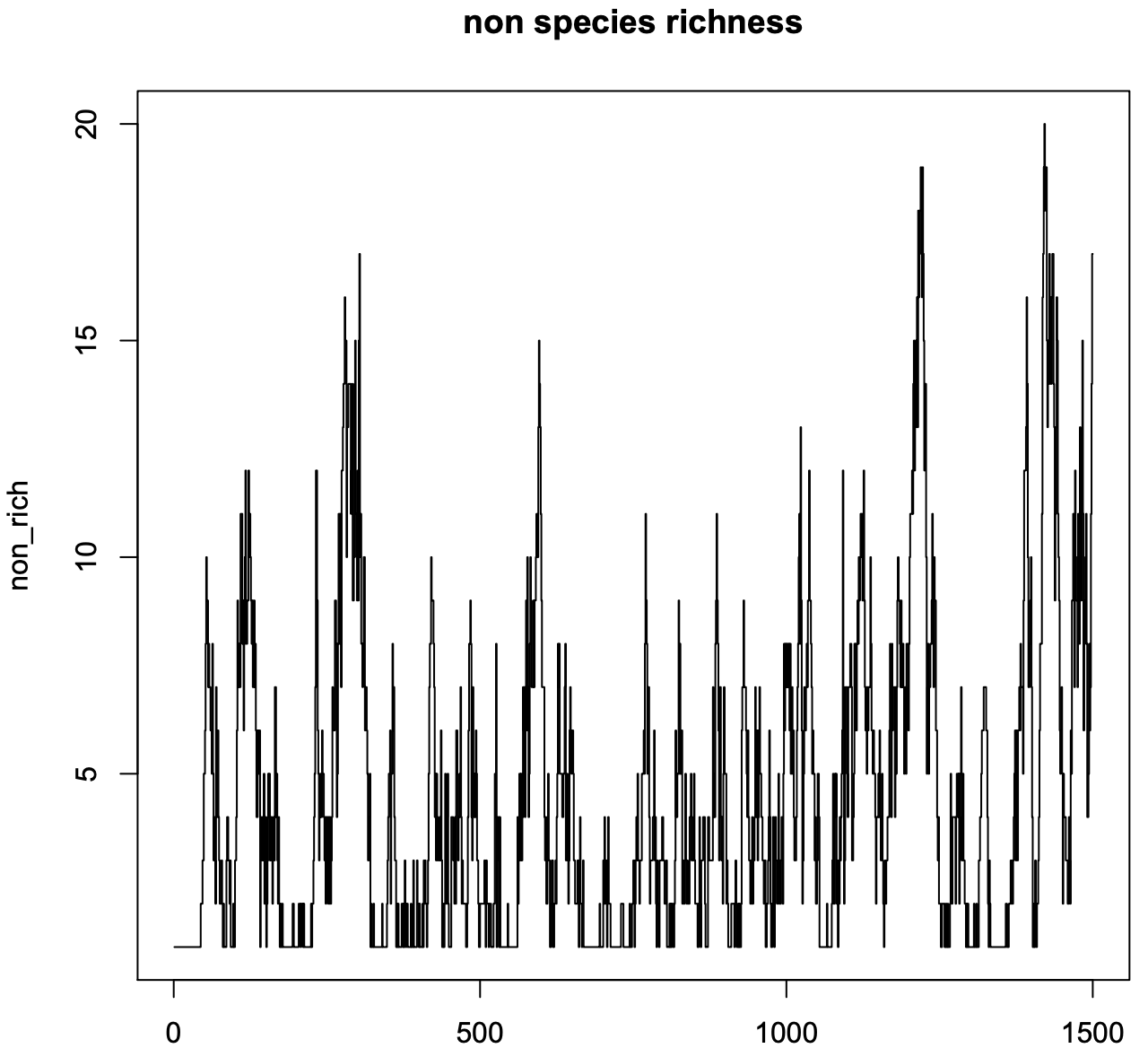
Figure S10. Change in dependent species richness over time, for Scenarios *a*. Shown here is one of the 96 replicates that were run for this scenario.

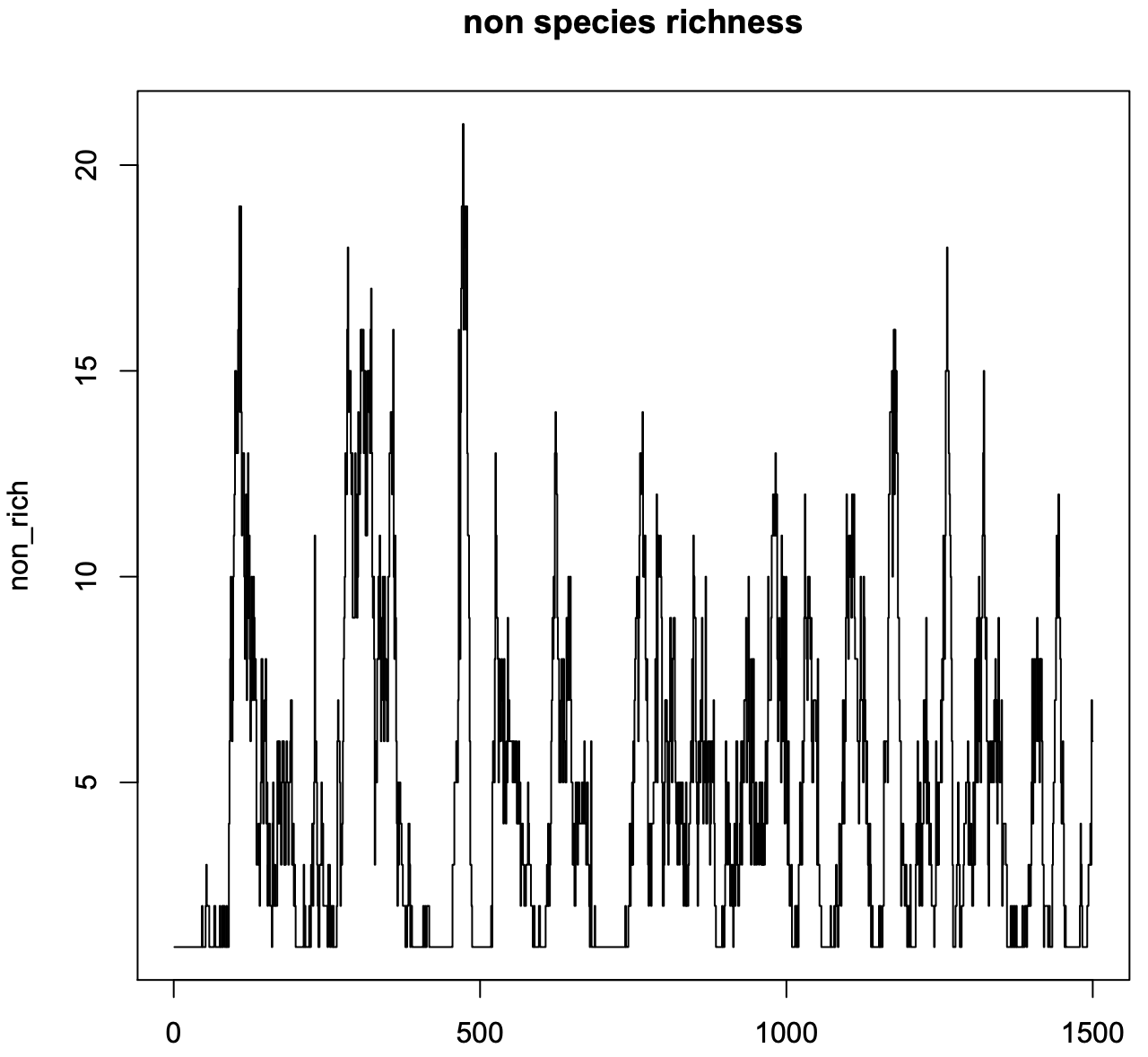
Figure S11. Change in dependent species richness over time, for Scenarios *b*. Shown here is one of the 96 replicates that were run for this scenario.

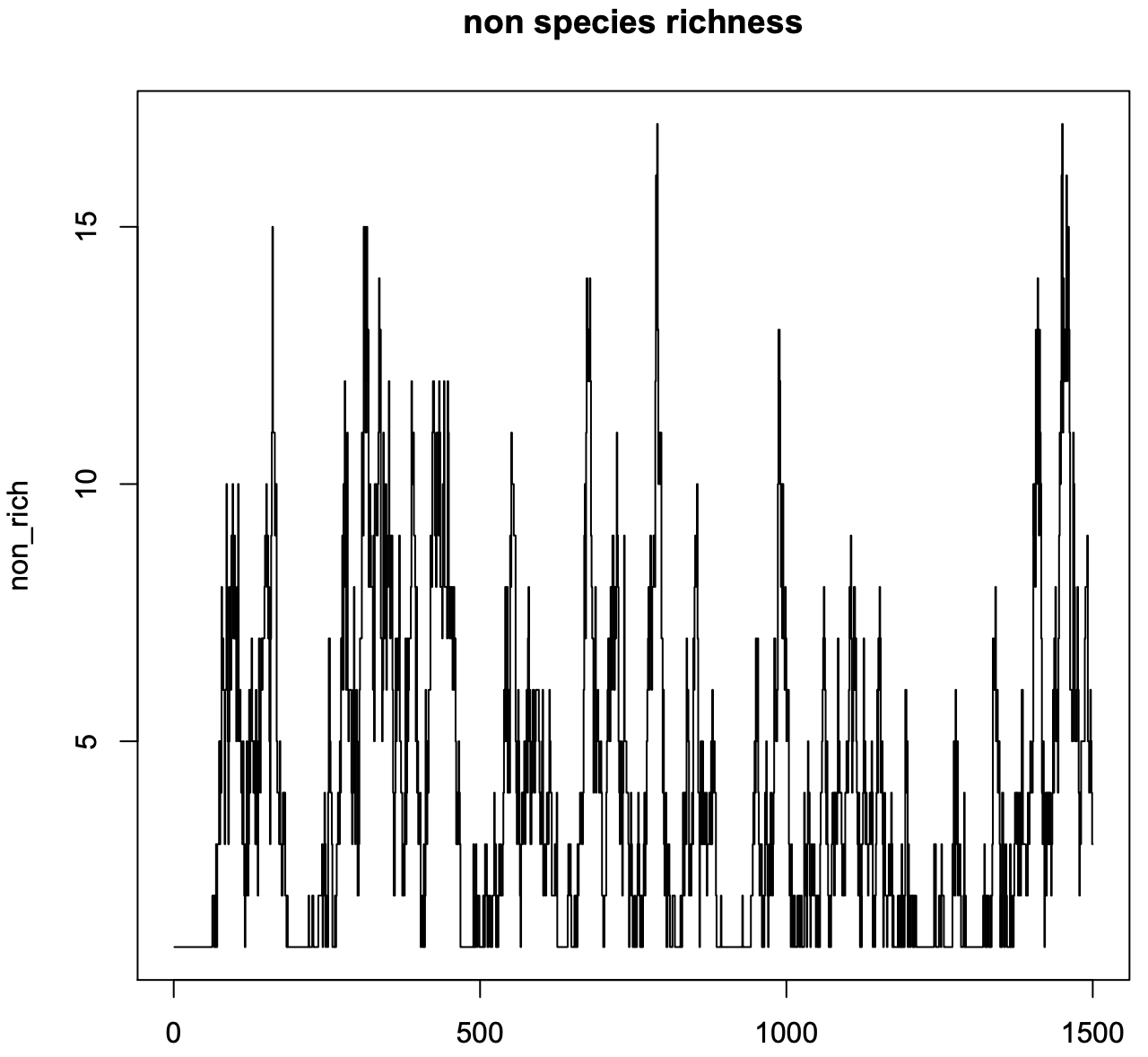
Figure S12. Change in dependent species richness over time, for Scenarios *c*. Shown here is one of the 96 replicates that were run for this scenario.

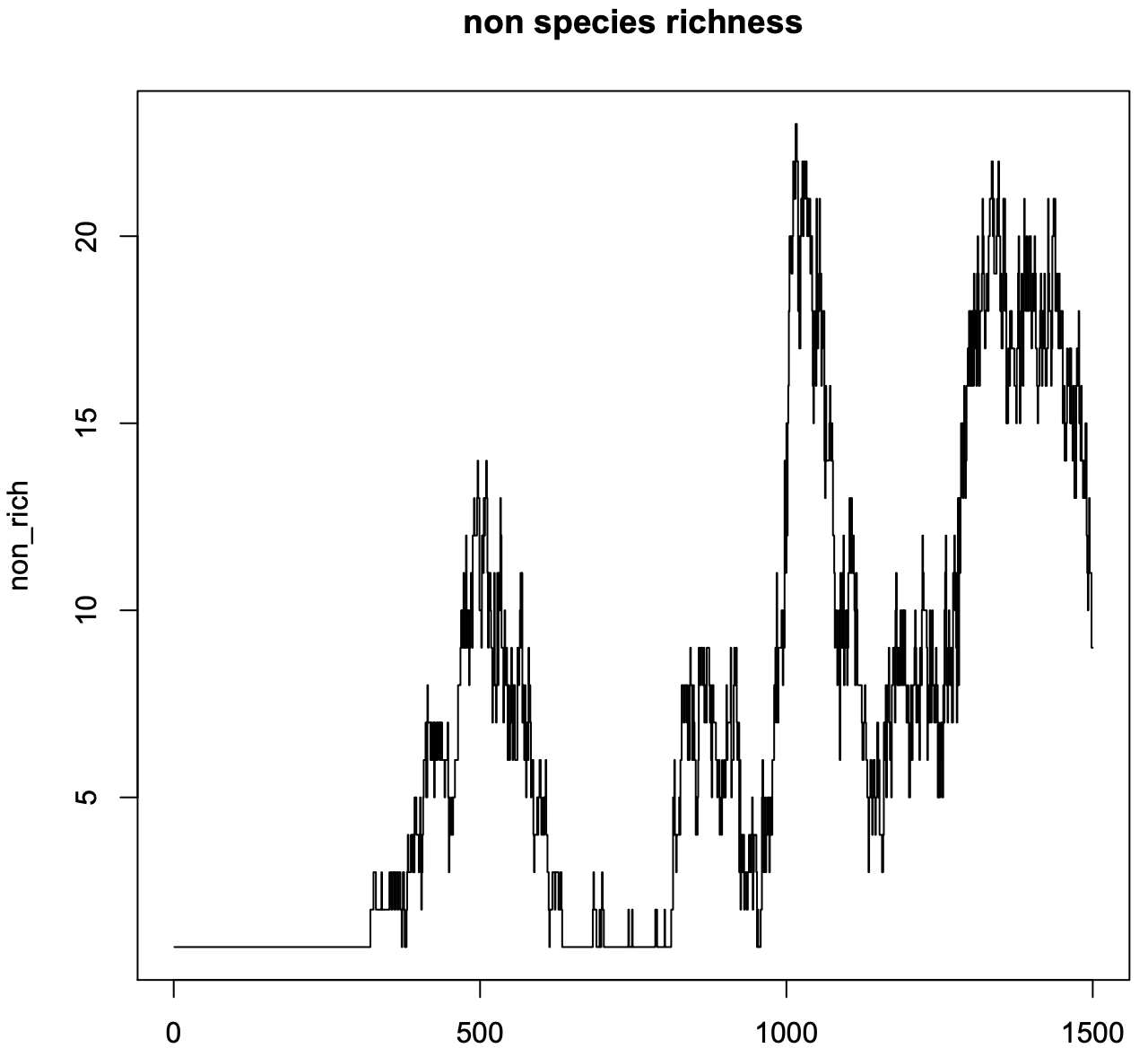
Figure S13. Change in dependent species richness over time, for Scenarios *d*. Shown here is one of the 96 replicates that were run for this scenario.

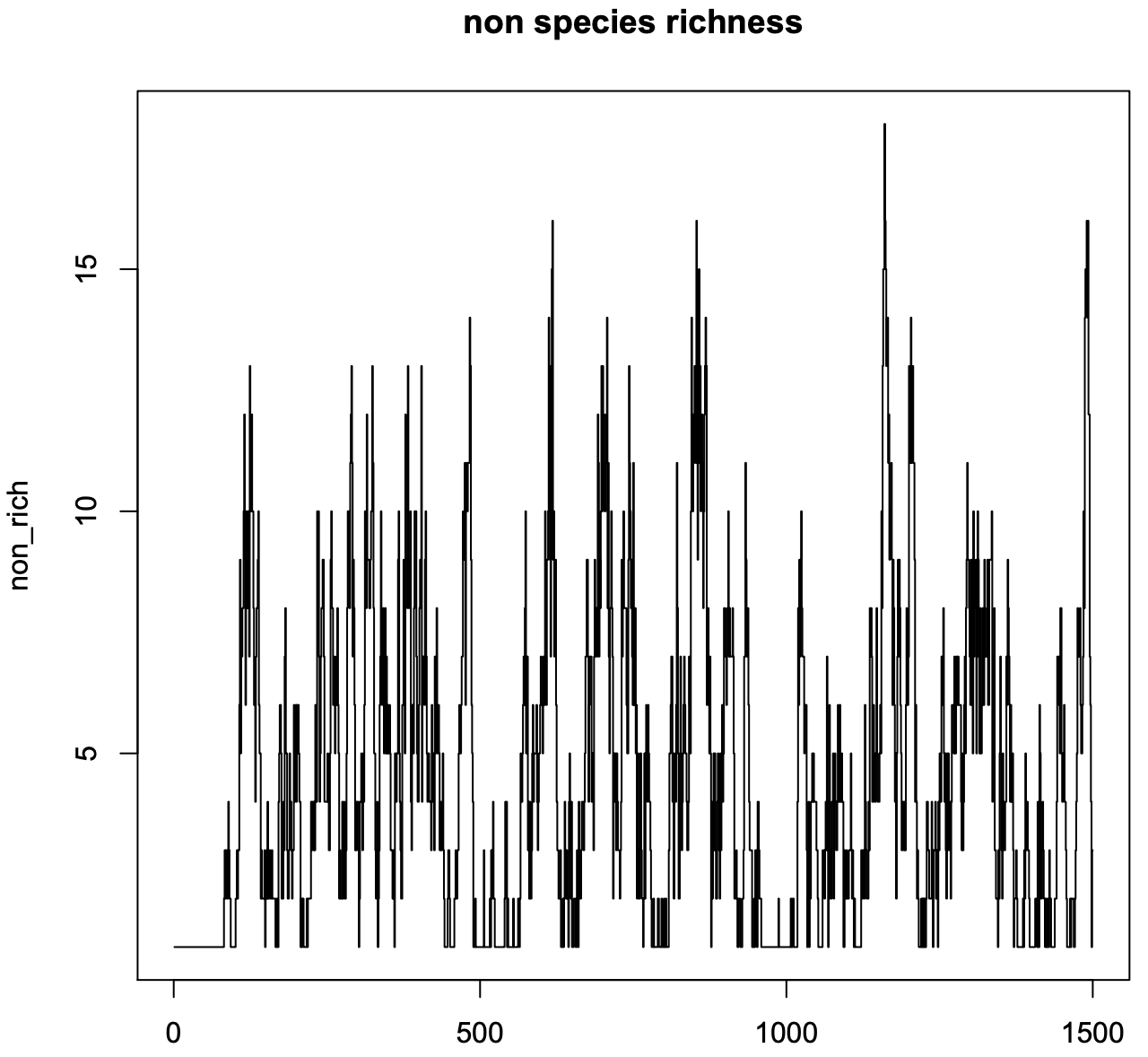
Figure S14. Change in dependent species richness over time, for Scenarios *e*. Shown here is one of the 96 replicates that were run for this scenario.

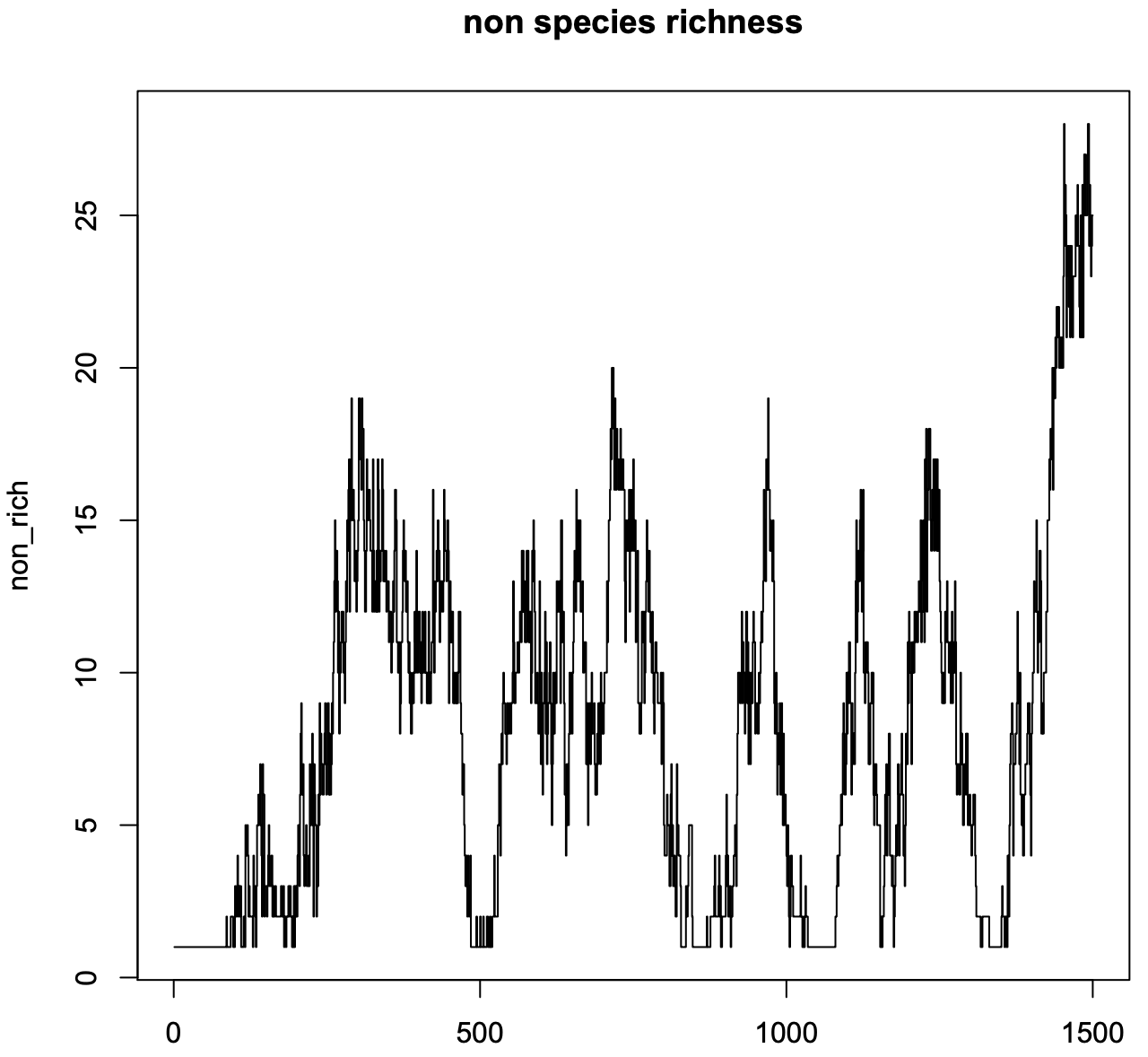
Figure S15. Change in dependent species richness over time, for Scenarios *f*. Shown here is one of the 96 replicates that were run for this scenario.

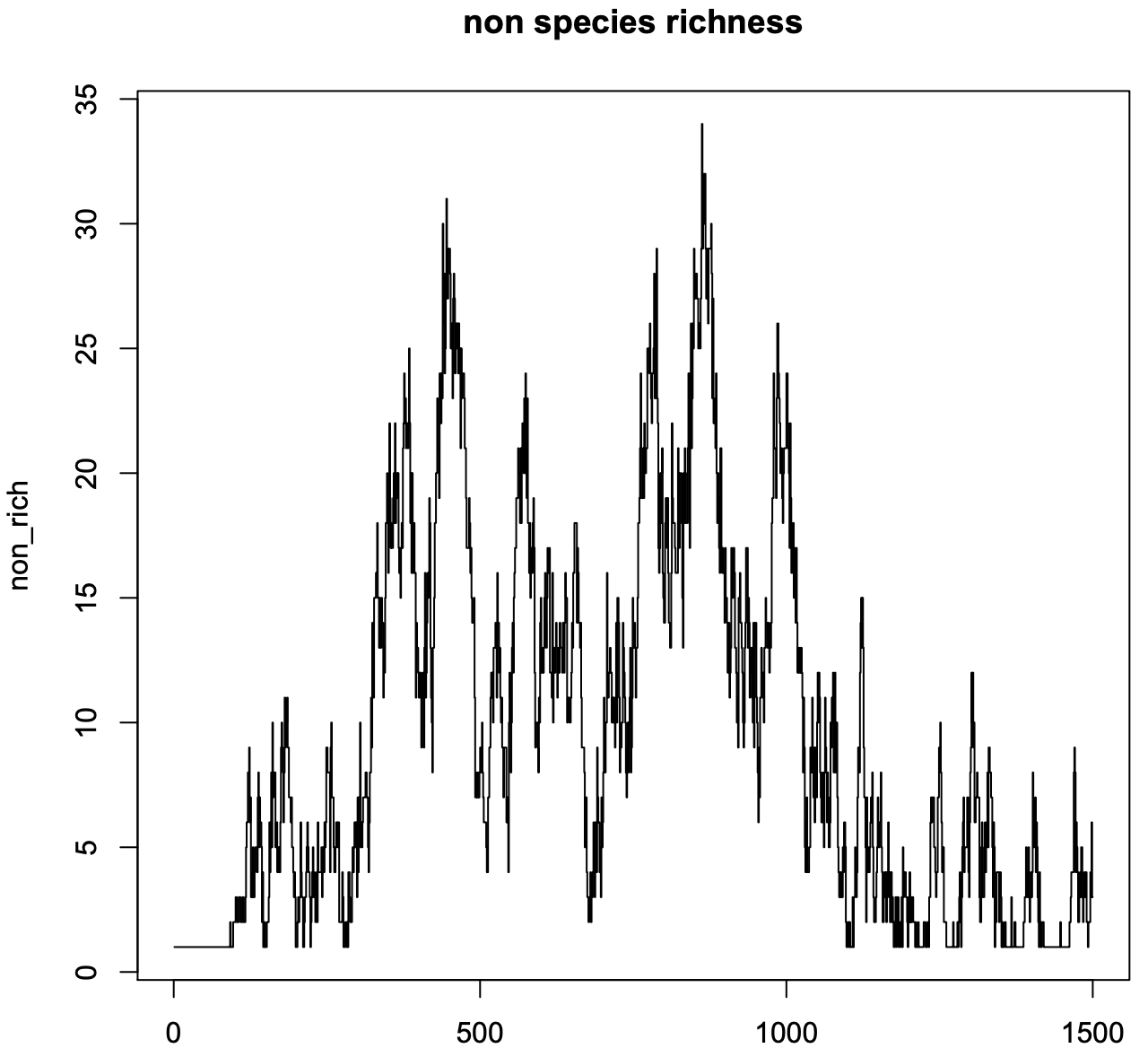
Figure S16. Change in dependent species richness over time, for Scenarios *g*. Shown here is one of the 96 replicates that were run for this scenario.

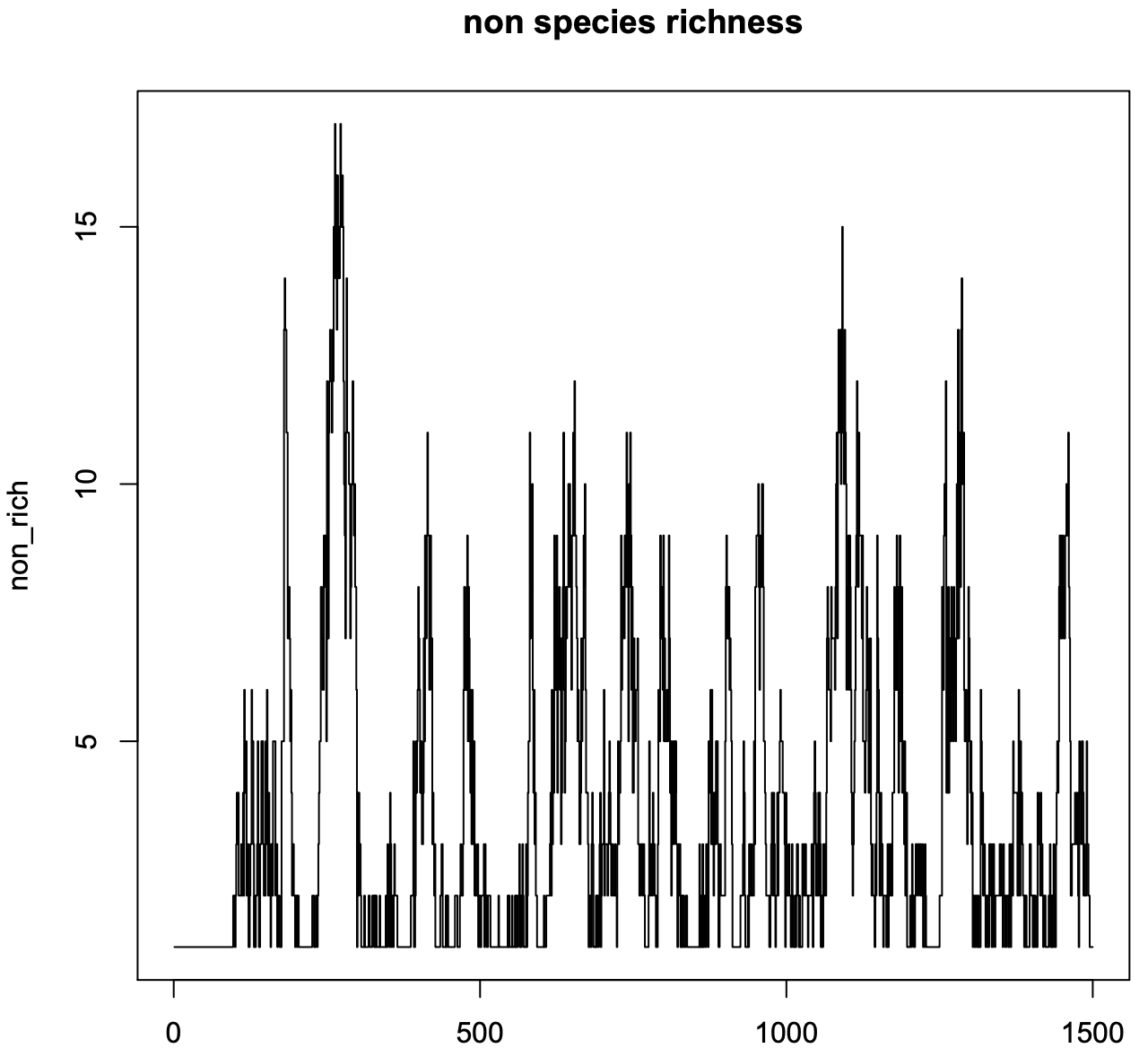
Figure S17. Change in dependent species richness over time, for Scenarios *h*. Shown here is one of the 96 replicates that were run for this scenario.

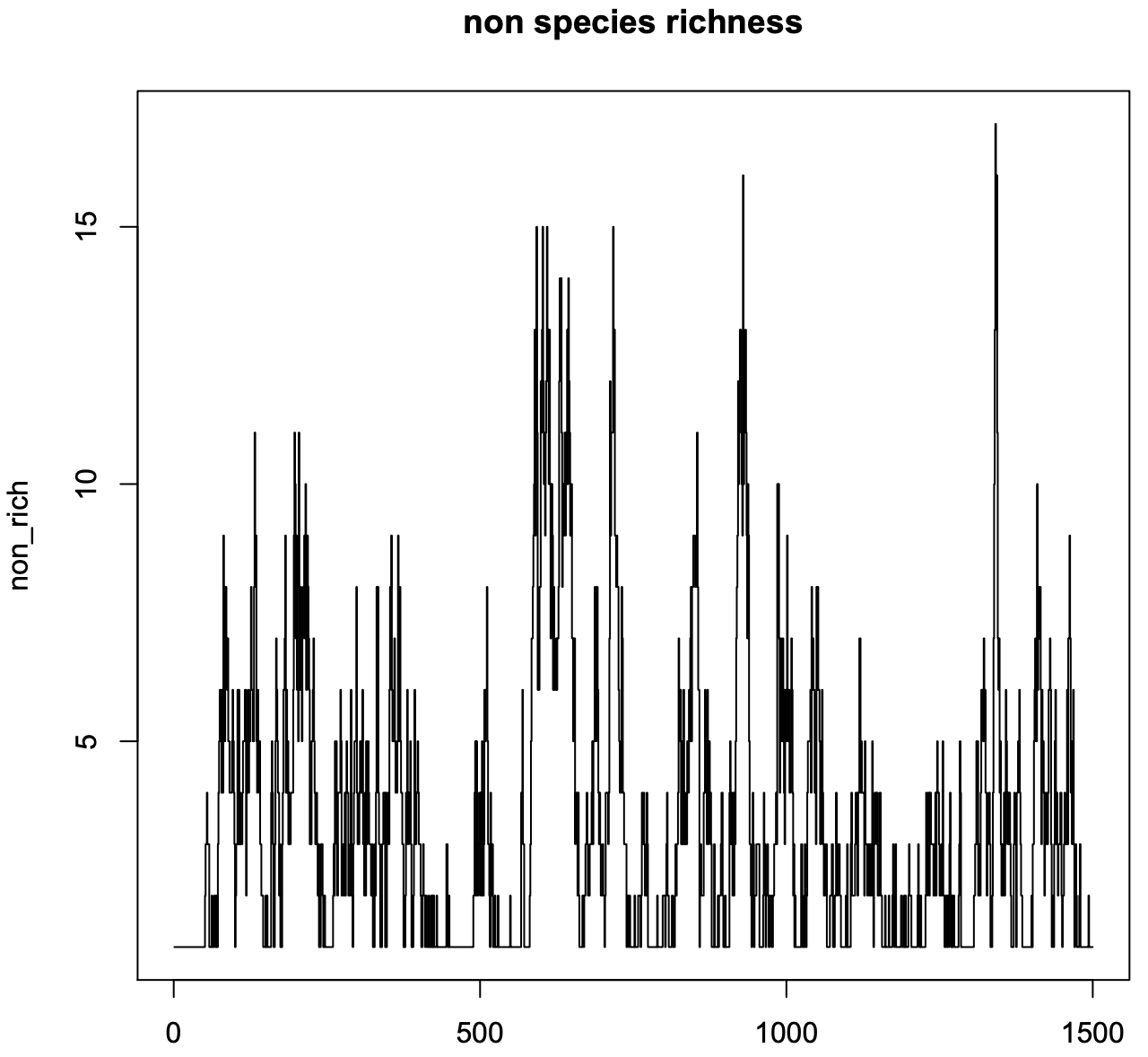
Figure S18. Change in dependent species richness over time, for Scenarios *i*. Shown here is one of the 96 replicates that were run for this scenario.

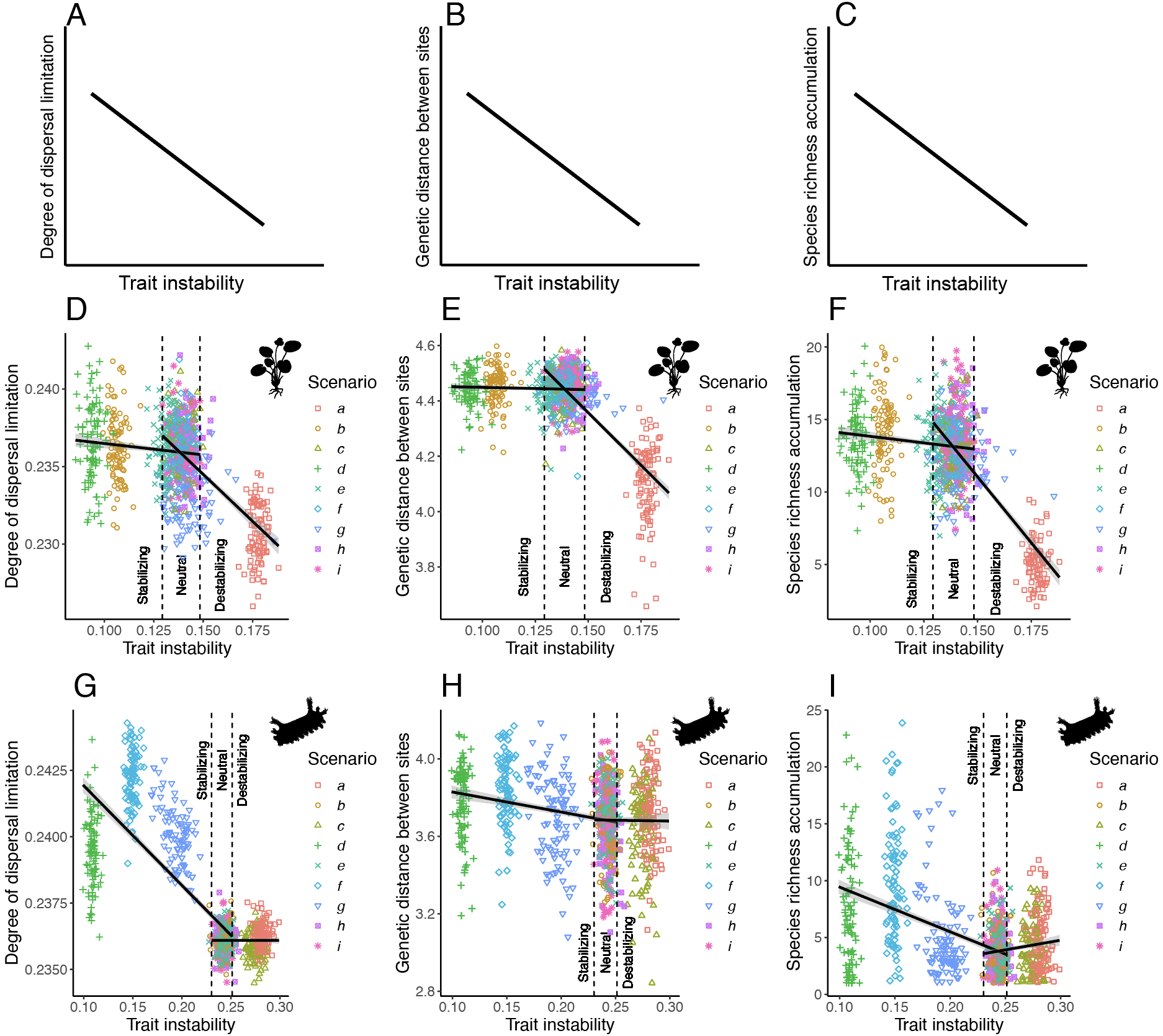

Figure S19. The linear regressions of the relationships between selective regime, degree of dispersal limitation, genetic distance among sites, and species richness accumulated. Selective regimes (neutral drift, stabilizing selection, or destabilizing selection) are determined by temporal trait stability measured as mean step difference $\bar{|\Delta z|}$. (A)-(C): Expected relationships – a significantly negative slope is predicted based on the stabilizing and destabilizing selection hypotheses. (D)-(F): Observed differences among selection regimes in the independent clade. (G)-(I): Observed differences among selection regimes in the dependent clade. In (D)-(I), black lines with gray ribbons show linear regressions with 95% confidence intervals. Two separate regressions were performed in each of these panels to visualize the differences between stabilizing selection and neutral drift and between destabilizing selection and neutral drift.
