## Supplementary material for "Coevolution-induced stabilizing and destabilizing selection shapes species richness in clade co-diversification": Table S1

Table S1. A list of all constants used in the simulations.

For the landscape:
*n* = 7

For both the independent and dependent clades:
*L_dist_* = 5
*T_dist_* = *L_dist_* – 1 = 4
*μ_dist_* = 0.03
*σ_phen_* = 0.1
*w_0_* = 0.01

For the independent clade:
*K_independent_* = 10
*n_mat_* = 3
*n_offspr_* = 2

For the dependent clade:
*K_dependent_* = 2
*n_mat_* = 9
*n_offspr_* = 6
