## Supplementary material for "Coevolution-induced stabilizing and destabilizing selection shapes species richness in clade co-diversification": File S1: all_in_one.R

```
###################################################
# function for getting ecological trait value
get_eco_trait_val <- function(ind){
  
  sum <-  0
  for (i in 1:length(ind$eco)){
    sum <- sum + ind$eco[[i]][1] + ind$eco[[i]][2]
  }
  
  return(sum)
}

###################################################
# function for getting choosiness trait value
get_choos_trait_val <- function(ind){
  
  sum <-  0
  for (i in 1:length(ind$pre)){
    sum <- sum + ind$pre[[i]][1] + ind$pre[[i]][2]
  }
  
  return(sum)
}

#######################################################
# function for calculating mating probability
mat_prob <- function(ind1, ind2, c_am){
  
  #get ecological trait value for both parents
  z1 <- get_eco_trait_val(ind1)
  z2 <- get_eco_trait_val(ind2)
  
  #get choosiness  trait value for both parents
  a1 <- get_choos_trait_val(ind1)
  a2 <- get_choos_trait_val(ind2)
  
  #use the choosier parent for probability calculation
  a <- a1
  if(abs(a1) < abs(a2)){
    a <- a2
  }

  temp1 <- 1-0.5*exp(-a^2) #1-0.5*exp(-a^2)
  temp2 <- exp( - (z1 -z2)^2 / (2/(a^2 * c_am))^2 )  #exp( - (z1 -z2)^2 / (2/(a^2 * c_am))^2 )
  
  temp <- temp1 * temp2
  
  if(a < 0){
    mat_prob <- 1 - temp
  }
  else if(a ==0){
    mat_prob <- 0.5
  }
  else if(a > 0){
    mat_prob <- temp
  }
  
  return(mat_prob)
  
}


#######################################################
# function for counting the number of loci with different compatibility alleles
incomp_count <- function(ind1, ind2){
  
  count <- 0
#  for (i in 1:L_b) {
#    if(ind1$post[[i]][1]==ind1$post[[i]][2] && ind2$post[[i]][1]==ind2$post[[i]][2] && ind1$post[[i]][1]==ind2$post[[i]][1]){
#      count <- count
#    }
#    else
#      count <- count + 1
#  }

  for (i in 1:L_b) {
    if(ind1$post[[i]][1]!=ind2$post[[i]][1] && ind1$post[[i]][1]!=ind2$post[[i]][2] && ind1$post[[i]][2]!=ind2$post[[i]][1] && ind1$post[[i]][2]!=ind2$post[[i]][2]){
      count <- count + 1
    }
    else
      count <- count
  }
  
  return(count)
}
```
