## Supplementary material for "Coevolution-induced stabilizing and destabilizing selection shapes species richness in clade co-diversification": File S1: basic_functions.R

```
#library(plot.matrix)
library(Rlab)
library(Rcpp)
library(stringr)
library(brio)
library(rlist)
library(foreach)
library(doParallel)
library(ape)
library(igraph)
library(wesanderson)

source("all_in_one.R")
source("coevolutionary_fitness_function.R")

# Functions that update the individual lists & the gird

#Update number of hosts in each cell for the grid (without barriers)
count_host_grid <- function(host_list, n_dimension){

  grid_host = matrix(0, nrow = n_dimension, ncol = n_dimension)

  for (i in 1:length(host_list))
  {
    # check if the individual is dead
    if (length(host_list[[i]]) == 8)
    {
      site.id <- host_list[[i]]$site.id
      grid_host[site.id[1],site.id[2]] <- grid_host[site.id[1],site.id[2]] + 1
    }
  }

  return(grid_host)

}

## Update number of non-hosts in each cell for the grid (without barriers)
count_non_grid <- function(non_list, n_dimension){

  grid_non = matrix(0, nrow = n_dimension, ncol = n_dimension)

  for (i in 1:length(non_list))
  {
    # check if the individual is dead
    if (length(non_list[[i]]) == 8)
    {
      site.id <- non_list[[i]]$site.id
      grid_non[site.id[1],site.id[2]] <- grid_non[site.id[1],site.id[2]] + 1
    }
  }

  return(grid_non)

}

## Update number of non-hosts on each host for the host list
count_non_host <- function(host_list, non_list){

  for (i in 1:length(host_list)){
    if(length(host_list[[i]]) == 8){ # check if the host still is alive and contains n_non
      host_list[[i]]$n_non <- 0
    }
  }

  for (i in 1:length(non_list)){
    if (length(non_list[[i]]) == 8) # check if the individual is dead
    {
      host.id <- non_list[[i]]$host.id
      host_list[[host.id]]$n_non <- host_list[[host.id]]$n_non + 1
    }
  }

  return(host_list)

}

## For the i-th host in a site, find its position in the host list
find_host <- function(host_list, pos, site.id){

  count <- 0 # count
  for (i in 1:length(host_list)){

    if(length(host_list[[i]]) == 8){
      if(identical(as.numeric(host_list[[i]]$site.id), as.numeric(site.id)) == T){
        count <- count + 1
      }
    }

    if(as.numeric(count) == as.numeric(pos)){
      break
    }
  }

  return(i)

}

## For the i-th non-host on a host, find its position in the non-host list
find_non <- function(non_list, pos, host.id){

  count <- 0 # count
  for (i in 1:length(non_list)){

    if(length(non_list[[i]]) == 8){
      if(non_list[[i]]$host.id == host.id){
        count <- count + 1
      }
    }

    if(count == as.numeric(pos)){
      break
    }
  }
  return(i)

}


### Host movement
### 1 = up, 2 = right, 3 = down, 4 = left, 5 = stay
host_ind_mov <- function(host_ind, grid_bar, n_dimension){
  if (length(host_ind) == 8){

    # initialization
    passable <- "NO"

    site.row <- host_ind$site.id[1]
    site.col <- host_ind$site.id[2]

    options <- 1:5
    probs <- c(p_dis_host/4, p_dis_host/4, p_dis_host/4, p_dis_host/4, (1-p_dis_host))

    while (passable == "NO") {

      decision <- options
      if(length(decision) > 1){
        decision <- sample(x = options, prob = probs, size = 1)
      }

      if (decision==1){
        if (site.row==1 || identical(grid_bar[2*site.row-2, 2*site.col-1], 99.9) == T){
          passable <- "NO"
        }
        else passable <- "YES"
      }

      else if (decision==2){
        if (site.col==n_dimension || identical(grid_bar[2*site.row-1, 2*site.col], 99.9) == T){
          passable <- "NO"
        }
        else passable <- "YES"
      }

      else if (decision==3){
        if (site.row==n_dimension || identical(grid_bar[2*site.row, 2*site.col-1], 99.9) == T){
          passable <- "NO"
        }
        else passable <- "YES"
      }

      else if (decision==4){
        if (site.col==1 || identical(grid_bar[2*site.row-1, 2*site.col-2], 99.9) == T){
          passable <- "NO"
        }
        else passable <- "YES"
      }

      else if (decision==5){
        passable <- "YES"
      }

      if(passable=="NO"){
          options_old <- options
          options <- options_old[options_old!=decision] #refining options
          probs <- probs[options_old!=decision] #refining options
      }


    }

    if (decision==1) site.row <- site.row - 1
    else if (decision==2) site.col <- site.col + 1
    else if (decision==3) site.row <- site.row + 1
    else if (decision==4) site.col <- site.col - 1

    host_ind$site.id[1] <- site.row
    host_ind$site.id[2] <- site.col
    
    # if the individuals moves rather than stays, label it as a "colonist"
    if (decision!=5) host_ind$origin <- "colonist"

  }

  return(host_ind)

}


#########################################################################3
### Non-host movement
non_ind_mov <- function(non_ind, grid_bar, n_dimension, grid_host, host_list){

  if(length(non_ind) == 8){

    # initialization
    passable <- "NO"

    site.row <- non_ind$site.id[1]
    site.col <- non_ind$site.id[2]

    host.id <- non_ind$host.id

    options <- 1:5
    probs <- c(p_dis_non/4, p_dis_non/4, p_dis_non/4, p_dis_non/4, (1-p_dis_non))

    while (passable == "NO") {

      decision <- options
      if(length(decision) > 1){
        decision <- sample(x = options, prob = probs, size = 1)
      }

      if (decision==1){
        if (site.row==1 || identical(grid_bar[2*site.row-2, 2*site.col-1], 99.9) == T){
          passable <- "NO"
        }
        else passable <- "YES"
      }

      else if (decision==2){
        if (site.col==n_dimension || identical(grid_bar[2*site.row-1, 2*site.col], 99.9) == T){
          passable <- "NO"
        }
        else passable <- "YES"
      }

      else if (decision==3){
        if (site.row==n_dimension || identical(grid_bar[2*site.row, 2*site.col-1], 99.9) == T){
          passable <- "NO"
        }
        else passable <- "YES"
      }

      else if (decision==4){
        if (site.col==1 || identical(grid_bar[2*site.row-1, 2*site.col-2], 99.9) == T){
          passable <- "NO"
        }
        else passable <- "YES"
      }

      else if (decision ==5){
        passable <- "YES"
      }

      if(passable=="NO"){
        options_old <- options
        options <- options_old[options_old!=decision] #refining options
        probs <- probs[options_old!=decision] #refining options
      }

    }

    if (decision==1) site.row <- site.row - 1
    else if (decision==2) site.col <- site.col + 1
    else if (decision==3) site.row <- site.row + 1
    else if (decision==4) site.col <- site.col - 1

    non_ind$site.id[1] <- site.row
    non_ind$site.id[2] <- site.col

    ## choose a host in the new site
    n_host <- grid_host[site.row,site.col]
    if(n_host==0){
      non_ind <- as.list("D_no_host") # the non-host-dies if the new site does not have any hosts
      #print("Die!")
    }
    else{
      host_pos <- sample(1:n_host, 1)
      non_ind$host.id <- find_host(host_list, host_pos, non_ind$site.id)
    }
    
    # if the individuals moves rather than stays, label it as a "colonist"
    if (decision!=5) non_ind$origin <- "colonist"

  }

  return(non_ind)

}
```
