## Supplementary material for "Coevolution-induced stabilizing and destabilizing selection shapes species richness in clade co-diversification": File S1: calc_div_turn.R

```
library(rlist)

# specify the name of the output folder
output_dir <- "Output_extinction"

# specify the number of replicates
n_rep <- 3

# generate a vector of the names of the simulations to be plotted
sim_names <- c( # all simulations
  "aa0010AMF",
  "aa0011AMF",
  "aa1010AMF",
  "aa1011AMF",
  "as0010AMF",
  "as0011AMF",
  "as1010AMF",
  "as1011AMF",
  "sa0010AMF",
  "sa0011AMF",
  "sa1010AMF",
  "sa1011AMF",
  "ss0010AMF",
  "ss0011AMF",
  "ss1010AMF",
  "ss1011AMF",
  "aa0110AMM",
  "aa0111AMM",
  "aa1110AMM",
  "aa1111AMM",
  "as0110AMM",
  "as0111AMM",
  "as1110AMM",
  "as1111AMM",
  "sa0110AMM",
  "sa0111AMM",
  "sa1110AMM",
  "sa1111AMM",
  "ss0110AMM",
  "ss0111AMM",
  "ss1110AMM",
  "ss1111AMM",
  "aa0100FFD",
  "aa0101FFD",
  "aa1100FFD",
  "aa1101FFD",
  "as0100FFD",
  "as0101FFD",
  "as1100FFD",
  "as1101FFD",
  "sa0100FFD",
  "sa0101FFD",
  "sa1100FFD",
  "sa1101FFD",
  "ss0100FFD",
  "ss0101FFD",
  "ss1100FFD",
  "ss1101FFD",
  "aa0000FFF",
  "aa0001FFF",
  "aa1000FFF",
  "aa1001FFF",
  "as0000FFF",
  "as0001FFF",
  "as1000FFF",
  "as1001FFF",
  "sa0000FFF",
  "sa0001FFF",
  "sa1000FFF",
  "sa1001FFF",
  "ss0000FFF",
  "ss0001FFF",
  "ss1000FFF",
  "ss1001FFF",
  "aa0100FFM",
  "aa0101FFM",
  "aa1100FFM",
  "aa1101FFM",
  "as0100FFM",
  "as0101FFM",
  "as1100FFM",
  "as1101FFM",
  "sa0100FFM",
  "sa0101FFM",
  "sa1100FFM",
  "sa1101FFM",
  "ss0100FFM",
  "ss0101FFM",
  "ss1100FFM",
  "ss1101FFM",
  "aa0110MDD",
  "aa0111MDD",
  "aa1110MDD",
  "aa1111MDD",
  "as0110MDD",
  "as0111MDD",
  "as1110MDD",
  "as1111MDD",
  "sa0110MDD",
  "sa0111MDD",
  "sa1110MDD",
  "sa1111MDD",
  "ss0110MDD",
  "ss0111MDD",
  "ss1110MDD",
  "ss1111MDD",
  "aa0010MDF",
  "aa0011MDF",
  "aa1010MDF",
  "aa1011MDF",
  "as0010MDF",
  "as0011MDF",
  "as1010MDF",
  "as1011MDF",
  "sa0010MDF",
  "sa0011MDF",
  "sa1010MDF",
  "sa1011MDF",
  "ss0010MDF",
  "ss0011MDF",
  "ss1010MDF",
  "ss1011MDF",
  "aa0010MMF",
  "aa0011MMF",
  "aa1010MMF",
  "aa1011MMF",
  "as0010MMF",
  "as0011MMF",
  "as1010MMF",
  "as1011MMF",
  "sa0010MMF",
  "sa0011MMF",
  "sa1010MMF",
  "sa1011MMF",
  "ss0010MMF",
  "ss0011MMF",
  "ss1010MMF",
  "ss1011MMF",
  "aa0110MMM",
  "aa0111MMM",
  "aa1110MMM",
  "aa1111MMM",
  "as0110MMM",
  "as0111MMM",
  "as1110MMM",
  "as1111MMM",
  "sa0110MMM",
  "sa0111MMM",
  "sa1110MMM",
  "sa1111MMM",
  "ss0110MMM",
  "ss0111MMM",
  "ss1110MMM",
  "ss1111MMM"
)


# calculate host and non-host diversity (species richness) and turnover rates
sp_host <- rep(NA, n_rep * length(sim_names))
ex_host <- rep(NA, n_rep * length(sim_names))

sp_non <- rep(NA, n_rep * length(sim_names))
ex_non <- rep(NA, n_rep * length(sim_names))

div_host <- rep(NA, n_rep * length(sim_names))
turn_host <- rep(NA, n_rep * length(sim_names))

div_non <- rep(NA, n_rep * length(sim_names))
turn_non <- rep(NA, n_rep * length(sim_names))

for (sim_pos in 1:length(sim_names)){
  for (rep_id in 1:n_rep) {
    print(sim_names[sim_pos])
    
    # read in the needed file
    file_name <- paste(output_dir, "/", as.character(sim_names[sim_pos]), "/", as.character(rep_id), "/pop_spec_host.rds", sep = '')
    pop_spec_host <- list.load(file_name)
    file_name <- paste(output_dir, "/", as.character(sim_names[sim_pos]), "/", as.character(rep_id), "/pop_spec_non.rds", sep = '')
    pop_spec_non <- list.load(file_name)
    
    # file_name <- paste(output_dir, "/", as.character(sim_names[sim_pos]), "/", as.character(rep_id), "/anc_spec_host.rds", sep = '')
    # anc_spec_host <- list.load(file_name)
    # file_name <- paste(output_dir, "/", as.character(sim_names[sim_pos]), "/", as.character(rep_id), "/anc_spec_non.rds", sep = '')
    # anc_spec_non <- list.load(file_name)
    
    # for host
    host_spec_rich <- rep(NA, length(pop_spec_host))
    for (g in 2:length(pop_spec_host)) {
      host_spec_rich[g] <- length(pop_spec_host[[g]]$host_spec_list)
    }
    # extinction <- rep(NA, length(pop_spec_host))
    # for (g in 2:length(pop_spec_host)) {
    #   extinction[g] <- length(pop_spec_host[[g - 1]]$host_spec_list) - length(unique(anc_spec_host[[g]])) # the number of lineages from the previous generation minus the number of lineages from the previous generation with decsendants from the current generation
    # }
    # speciation <- rep(NA, length(pop_spec_host))
    # for (g in 2:length(pop_spec_host)) {
    #   speciation[g] <- length(pop_spec_host[[g]]$host_spec_list) - length(unique(anc_spec_host[[g]])) # the number of current lineages minus the number of ancestral lineages from the previous generation
    # }
    # net_div <- speciation - extinction
    # turnover <- speciation + extinction
    div_host[(sim_pos - 1) * n_rep + rep_id] <- mean(host_spec_rich[(length(host_spec_rich) - 10) : (length(host_spec_rich) - 0)]) # only use the last 100 generations to ensure macroevolutioanry equilibrium
    # turn_host[(sim_pos - 1) * n_rep + rep_id] <- mean(turnover[(length(turnover) - 10) : length(turnover)]) # only use the last 100 generations to ensure macroevolutioanry equilibrium
    # 
    # sp_host[(sim_pos - 1) * n_rep + rep_id] <- mean(speciation[(length(speciation) - 10) : length(speciation)])
    # ex_host[(sim_pos - 1) * n_rep + rep_id] <- mean(extinction[(length(extinction) - 10) : length(extinction)])
    
    # for non
    non_spec_rich <- rep(NA, length(pop_spec_non))
    for (g in 2:length(pop_spec_non)) {
      non_spec_rich[g] <- length(pop_spec_non[[g]]$non_spec_list)
    }
    # extinction <- rep(NA, length(pop_spec_non))
    # for (g in 2:length(pop_spec_non)) {
    #   extinction[g] <- length(pop_spec_non[[g - 1]]$non_spec_list) - length(unique(anc_spec_non[[g]])) # the number of lineages from the previous generation minus the number of lineages from the previous generation with decsendants from the current generation
    # }
    # speciation <- rep(NA, length(pop_spec_non))
    # for (g in 2:length(pop_spec_non)) {
    #   speciation[g] <- length(pop_spec_non[[g]]$non_spec_list) - length(unique(anc_spec_non[[g]])) # the number of current lineages minus the number of ancestral lineages from the previous generation
    # }
    # net_div <- speciation - extinction
    # turnover <- speciation + extinction
    div_non[(sim_pos - 1) * n_rep + rep_id] <- mean(non_spec_rich[(length(non_spec_rich) - 10) : (length(non_spec_rich) - 0)]) # only use the last 100 generations to ensure macroevolutioanry equilibrium
    # turn_non[(sim_pos - 1) * n_rep + rep_id] <- mean(turnover[(length(turnover) - 10) : length(turnover)]) # only use the last 100 generations to ensure macroevolutioanry equilibrium
    # 
    # sp_non[(sim_pos - 1) * n_rep + rep_id] <- mean(speciation[(length(speciation) - 10) : length(speciation)])
    # ex_non[(sim_pos - 1) * n_rep + rep_id] <- mean(extinction[(length(extinction) - 10) : length(extinction)])
  }
}


# create a vector recording which treatment each value belongs to
nam <- rep(NA, length(sim_names) * n_rep)
for ( rep_id in 1:n_rep) {
  nam[(1:length(sim_names)) * n_rep - (n_rep - rep_id)] <- sim_names
}

# create a data frame
div_turn <- data.frame(treatment=nam, div_host=div_host, div_non=div_non, turn_host=turn_host, turn_non=turn_non, sp_host = sp_host, ex_host = ex_host, sp_non = sp_non, ex_non = ex_non)

# (do not do this for formal analysis) make pseudo-replicates to get a sense of the effects of sample size
# nest_mod <- rbind(nest_mod, nest_mod, nest_mod, nest_mod, nest_mod, nest_mod, nest_mod, nest_mod, nest_mod, nest_mod, nest_mod, nest_mod, nest_mod, nest_mod, nest_mod, nest_mod, nest_mod, nest_mod, nest_mod, nest_mod, nest_mod)

# save the data frame to a spreadsheet
write.csv(div_turn, paste(output_dir, "/div_turn.csv", sep = ''))
```
