## Supplementary material for "Coevolution-induced stabilizing and destabilizing selection shapes species richness in clade co-diversification": File S1: calc_hybr_lik.R

```
library(rlist)
library(Microsoft365R)
library(scales)

library(future)
library(doParallel)
library(parallel)
n.cores <- future::availableCores()
registerDoParallel(cores=n.cores - 8)

# print the number of slaves
print("The number of slaves is:")
print(n.cores - 8)

sim_nam <- "aa0000FFF" # this can be anything
source('basic_functions.R')
source('all_in_one.R')
source('reproduction_pop.R')
source('geographic_dynamics.R')
source('env_optima.R')
source('initialization/common.R')

# specify the number of replicates and the total number of generations simulated
n_rep <- 3
n_gen <- 1500

# specify the size the of landscape
n_dimension <- 7

# generate a vector of the names of the simulations to be plotted
sim_names <- c( # all simulations
  "aa0010AMF",
  "aa0011AMF",
  "aa1010AMF",
  "aa1011AMF",
  "as0010AMF",
  "as0011AMF",
  "as1010AMF",
  "as1011AMF",
  "sa0010AMF",
  "sa0011AMF",
  "sa1010AMF",
  "sa1011AMF",
  "ss0010AMF",
  "ss0011AMF",
  "ss1010AMF",
  "ss1011AMF",
  "aa0110AMM",
  "aa0111AMM",
  "aa1110AMM",
  "aa1111AMM",
  "as0110AMM",
  "as0111AMM",
  "as1110AMM",
  "as1111AMM",
  "sa0110AMM",
  "sa0111AMM",
  "sa1110AMM",
  "sa1111AMM",
  "ss0110AMM",
  "ss0111AMM",
  "ss1110AMM",
  "ss1111AMM",
  "aa0100FFD",
  "aa0101FFD",
  "aa1100FFD",
  "aa1101FFD",
  "as0100FFD",
  "as0101FFD",
  "as1100FFD",
  "as1101FFD",
  "sa0100FFD",
  "sa0101FFD",
  "sa1100FFD",
  "sa1101FFD",
  "ss0100FFD",
  "ss0101FFD",
  "ss1100FFD",
  "ss1101FFD",
  "aa0000FFF",
  "aa0001FFF",
  "aa1000FFF",
  "aa1001FFF",
  "as0000FFF",
  "as0001FFF",
  "as1000FFF",
  "as1001FFF",
  "sa0000FFF",
  "sa0001FFF",
  "sa1000FFF",
  "sa1001FFF",
  "ss0000FFF",
  "ss0001FFF",
  "ss1000FFF",
  "ss1001FFF",
  "aa0100FFM",
  "aa0101FFM",
  "aa1100FFM",
  "aa1101FFM",
  "as0100FFM",
  "as0101FFM",
  "as1100FFM",
  "as1101FFM",
  "sa0100FFM",
  "sa0101FFM",
  "sa1100FFM",
  "sa1101FFM",
  "ss0100FFM",
  "ss0101FFM",
  "ss1100FFM",
  "ss1101FFM",
  "aa0110MDD",
  "aa0111MDD",
  "aa1110MDD",
  "aa1111MDD",
  "as0110MDD",
  "as0111MDD",
  "as1110MDD",
  "as1111MDD",
  "sa0110MDD",
  "sa0111MDD",
  "sa1110MDD",
  "sa1111MDD",
  "ss0110MDD",
  "ss0111MDD",
  "ss1110MDD",
  "ss1111MDD",
  "aa0010MDF",
  "aa0011MDF",
  "aa1010MDF",
  "aa1011MDF",
  "as0010MDF",
  "as0011MDF",
  "as1010MDF",
  "as1011MDF",
  "sa0010MDF",
  "sa0011MDF",
  "sa1010MDF",
  "sa1011MDF",
  "ss0010MDF",
  "ss0011MDF",
  "ss1010MDF",
  "ss1011MDF",
  "aa0010MMF",
  "aa0011MMF",
  "aa1010MMF",
  "aa1011MMF",
  "as0010MMF",
  "as0011MMF",
  "as1010MMF",
  "as1011MMF",
  "sa0010MMF",
  "sa0011MMF",
  "sa1010MMF",
  "sa1011MMF",
  "ss0010MMF",
  "ss0011MMF",
  "ss1010MMF",
  "ss1011MMF",
  "aa0110MMM",
  "aa0111MMM",
  "aa1110MMM",
  "aa1111MMM",
  "as0110MMM",
  "as0111MMM",
  "as1110MMM",
  "as1111MMM",
  "sa0110MMM",
  "sa0111MMM",
  "sa1110MMM",
  "sa1111MMM",
  "ss0110MMM",
  "ss0111MMM",
  "ss1110MMM",
  "ss1111MMM"
)

# specify the generations to look at (the last 100 generations to ensure equilibrium)
gens <- (n_gen - 10 + 1):n_gen

# # initializae the vectors for recording calculated values
# hybr_lik_host <- rep(NA, n_rep * length(sim_names))
# hybr_lik_non <- rep(NA, n_rep * length(sim_names))
# 
# # initializae the vectors for recording the type of calculated value (mat_prob vs. gen_incomp)
# hybr_lik_host_mode <- rep(NA, n_rep * length(sim_names))
# hybr_lik_non_mode <- rep(NA, n_rep * length(sim_names))
# 
# # start calculation
# od <- get_business_onedrive()
# for (sim_pos in 1:length(sim_names)){
#   sim_id <- sim_names[sim_pos]
#   print(sim_names[sim_pos])
#   
#   for (rep_id in 1:n_rep) {
#     
#     print(rep_id)
#     
#     # initialize a vector recording the mean hybridization_likelihood for the last 10% of all generations
#     hybr_lik_host_gens <- rep(NA, n_gen)
#     hybr_lik_non_gens <- rep(NA, n_gen)
#     
#     # read in the generations file
#     file_name <- paste(output_dir, '/', sim_id, "/", rep_id, "/generations.rds", sep = '')
#     od$download_file(paste("model_rcpp_hpc_OneDrive/", file_name, sep = ''), file_name, overwrite = T)
#     generations <- list.load(file_name)
#     unlink(file_name)
#     
#     # read in the population and species info
#     pop_spec_host <- list.load( paste(output_dir, '/', sim_id, "/", rep_id, "/pop_spec_host.rds", sep = '') )
#     pop_spec_non <- list.load( paste(output_dir, '/', sim_id, "/", rep_id, "/pop_spec_non.rds", sep = '') )
#     
#     # for each generation
#     for(gen_id in gens){
#       
#       host_list <- generations[[gen_id]]$host_list
#       non_list <- generations[[gen_id]]$non_list
#       
#       host_spec_list <- pop_spec_host[[gen_id]]$host_spec_list
#       non_spec_list <- pop_spec_non[[gen_id]]$non_spec_list
#       
#       host_pop_list <- pop_spec_host[[gen_id]]$host_pop_list
#       non_pop_list <- pop_spec_non[[gen_id]]$non_pop_list
#       
#       # decide whther the mating probabilty or genetic compatibility should be calculated
#       if(substr(sim_id, 1, 1) == 'a'){ # if geographic speciation
#         host_mode <- "gen_incomp"
#       }else if(substr(sim_id, 1, 1) == 's'){ # if ecological speciation
#         host_mode <- "mat_prob"
#       }
#       hybr_lik_host_mode[(sim_pos - 1) * n_rep + rep_id] <- host_mode
#       
#       if(substr(sim_id, 2, 2) == 'a'){ # if geographic speciation
#         non_mode <- "gen_incomp"
#       }else if(substr(sim_id, 2, 2) == 's'){ # if ecological speciation
#         non_mode <- "mat_prob"
#       }
#       hybr_lik_non_mode[(sim_pos - 1) * n_rep + rep_id] <- non_mode
#       
#       
#       
#       ########################################################################################
#       # for hosts
#       if(length(host_spec_list) < 2){ # hybridization cannot occur if there is only one species
#         hybr_lik_host_gens[gen_id] <- NA
#       }
#       else{
#         
#         # initialzie matrices recording the hybridization likelihood for each pair of species
#         hybr_lik_host_spec <- matrix(NA, nrow = length(host_spec_list), ncol = length(host_spec_list))
#         
#         # fill in the matrix
#         for (spec_id1 in 2:length(host_spec_list)) { # only calculate the bottom left half of the matrix
#           for (spec_id2 in 1:(spec_id1 - 1)) {
#             
#             # find all the individual IDs of the two species
#             inds1 <- unlist(host_pop_list[host_spec_list[[spec_id1]]])
#             inds2 <- unlist(host_pop_list[host_spec_list[[spec_id2]]])
#             
#             # calculate the save the mating probability or genetic compatibility
#             if(host_mode == "mat_prob"){
#               hybr_lik_host_spec[spec_id1, spec_id2] <- mat_prob_pop(inds1, inds2, host_list, host_list, c_am)
#             }else if(host_mode == "gen_incomp"){
#               hybr_lik_host_spec[spec_id1, spec_id2] <- incomp_count_pop(inds1, inds2, host_list, host_list)
#             }
#             
#           }
#         }
#         
#         hybr_lik_host_gens[gen_id] <- mean(hybr_lik_host_spec, na.rm = T) # take the mean across all species pairs  
#         
#       }
#       
# 
#       
#       ########################################################################################
#       # for nons
#       if(length(non_spec_list) < 2){ # hybridization cannot occur if there is only one species
#         hybr_lik_non_gens[gen_id] <- NA
#       }
#       else{
#         
#         # initialzie matrices recording the hybridization likelihood for each pair of species
#         hybr_lik_non_spec <- matrix(NA, nrow = length(non_spec_list), ncol = length(non_spec_list))
#         
#         # fill in the matrix
#         for (spec_id1 in 2:length(non_spec_list)) { # only calculate the bottom left half of the matrix
#           for (spec_id2 in 1:(spec_id1 - 1)) {
#             
#             # find all the individual IDs of the two species
#             inds1 <- unlist(non_pop_list[non_spec_list[[spec_id1]]])
#             inds2 <- unlist(non_pop_list[non_spec_list[[spec_id2]]])
#             
#             # calculate the save the mating probability or genetic compatibility
#             if(non_mode == "mat_prob"){
#               hybr_lik_non_spec[spec_id1, spec_id2] <- mat_prob_pop(inds1, inds2, non_list, non_list, c_am)
#             }else if(non_mode == "gen_incomp"){
#               hybr_lik_non_spec[spec_id1, spec_id2] <- incomp_count_pop(inds1, inds2, non_list, non_list)
#             }
#             
#           }
#         }
#         
#         hybr_lik_non_gens[gen_id] <- mean(hybr_lik_non_spec, na.rm = T) # take the mean across all species pairs    
#         
#       }
#       
# 
#       
#       
#       
#       
#       
#     }
#     
#     # take the mean across all generations
#     hybr_lik_host[(sim_pos - 1) * n_rep + rep_id] <- mean(hybr_lik_host_gens, na.rm = T)
#     hybr_lik_non[(sim_pos - 1) * n_rep + rep_id] <- mean(hybr_lik_non_gens, na.rm = T)
#     
#   }
# }
# 
# # save the data to a data frame
# hybr_lik <- data.frame(hybr_lik_host, hybr_lik_non, hybr_lik_host_mode, hybr_lik_non_mode)
# write.csv(hybr_lik, paste(output_dir, '/hybr_lik.csv', sep = ''))


# create a vector for parallelization
para_vec <- rep(NA, length(sim_names) * n_rep)
for (i in 1:length(sim_names)) {
  for (j in 1:n_rep) {
    para_vec[(i-1) * n_rep + j] <- paste(sim_names[i], as.character(j), sep = '_')
  }
}

# start calculation
od <- NULL
while (length(od) == 0) {
  try( # try again if it fails
    od <- get_business_onedrive(),
    TRUE
  )  
}

hybr_lik <- foreach (para_id = para_vec) %dopar% {
  
  sim_id <- substr(para_id, 1, 9)
  rep_id <- substr(para_id, 11, nchar(para_id))
  print(percent((which(para_vec == para_id)-1)/length(para_vec), accuracy = 1))
  
  # initialize a vector recording the mean hybridization_likelihood for the last 10% of all generations
  hybr_lik_host_gens <- rep(NA, n_gen)
  hybr_lik_non_gens <- rep(NA, n_gen)
  
  # read in the generations file
  generations <- NULL
  file_name <- paste(output_dir, '/', sim_id, "/", rep_id, "/generations.rds", sep = '')
  while (length(generations) == 0) {
    try( # try again if it fails
      od$download_file(paste("model_rcpp_hpc_OneDrive/", file_name, sep = ''), file_name, overwrite = T),
      TRUE
    )
    try( # try again if it fails
      generations <- list.load(file_name),
      TRUE
    )    
  }
  unlink(file_name)
  
  # read in the population and species info
  pop_spec_host <- list.load( paste(output_dir, '/', sim_id, "/", rep_id, "/pop_spec_host.rds", sep = '') )
  pop_spec_non <- list.load( paste(output_dir, '/', sim_id, "/", rep_id, "/pop_spec_non.rds", sep = '') )
  
  # for each generation
  for(gen_id in gens){
    
    host_list <- generations[[gen_id]]$host_list
    non_list <- generations[[gen_id]]$non_list
    
    host_spec_list <- pop_spec_host[[gen_id]]$host_spec_list
    non_spec_list <- pop_spec_non[[gen_id]]$non_spec_list
    
    host_pop_list <- pop_spec_host[[gen_id]]$host_pop_list
    non_pop_list <- pop_spec_non[[gen_id]]$non_pop_list
    
    # decide whther the mating probabilty or genetic compatibility should be calculated
    #if(substr(sim_id, 1, 1) == 'a'){ # if geographic speciation
      host_mode <- "gen_incomp"
    #}else if(substr(sim_id, 1, 1) == 's'){ # if ecological speciation
    #  host_mode <- "mat_prob"
    #}
    #hybr_lik_host_mode[(sim_pos - 1) * n_rep + rep_id] <- host_mode
    
    #if(substr(sim_id, 2, 2) == 'a'){ # if geographic speciation
      non_mode <- "gen_incomp"
    #}else if(substr(sim_id, 2, 2) == 's'){ # if ecological speciation
    #  non_mode <- "mat_prob"
    #}
    #hybr_lik_non_mode[(sim_pos - 1) * n_rep + rep_id] <- non_mode
    
    
    
    ########################################################################################
    # for hosts
    if(length(host_spec_list) < 2){ # hybridization cannot occur if there is only one species
      hybr_lik_host_gens[gen_id] <- NA
    }
    else{
      
      # initialzie matrices recording the hybridization likelihood for each pair of species
      hybr_lik_host_spec <- matrix(NA, nrow = length(host_spec_list), ncol = length(host_spec_list))
      
      # fill in the matrix
      for (spec_id1 in 2:length(host_spec_list)) { # only calculate the bottom left half of the matrix
        for (spec_id2 in 1:(spec_id1 - 1)) {
          
          # find all the individual IDs of the two species
          inds1 <- unlist(host_pop_list[host_spec_list[[spec_id1]]])
          inds2 <- unlist(host_pop_list[host_spec_list[[spec_id2]]])
          
          # calculate the save the mating probability or genetic compatibility
          if(host_mode == "mat_prob"){
            hybr_lik_host_spec[spec_id1, spec_id2] <- mat_prob_pop(inds1, inds2, host_list, host_list, c_am)
          }else if(host_mode == "gen_incomp"){
            hybr_lik_host_spec[spec_id1, spec_id2] <- incomp_count_pop(inds1, inds2, host_list, host_list)
          }
          
        }
      }
      
      hybr_lik_host_gens[gen_id] <- mean(hybr_lik_host_spec, na.rm = T) # take the mean across all species pairs  
      
    }
    
    
    
    ########################################################################################
    # for nons
    if(length(non_spec_list) < 2){ # hybridization cannot occur if there is only one species
      hybr_lik_non_gens[gen_id] <- NA
    }
    else{
      
      # initialzie matrices recording the hybridization likelihood for each pair of species
      hybr_lik_non_spec <- matrix(NA, nrow = length(non_spec_list), ncol = length(non_spec_list))
      
      # fill in the matrix
      for (spec_id1 in 2:length(non_spec_list)) { # only calculate the bottom left half of the matrix
        for (spec_id2 in 1:(spec_id1 - 1)) {
          
          # find all the individual IDs of the two species
          inds1 <- unlist(non_pop_list[non_spec_list[[spec_id1]]])
          inds2 <- unlist(non_pop_list[non_spec_list[[spec_id2]]])
          
          # calculate the save the mating probability or genetic compatibility
          if(non_mode == "mat_prob"){
            hybr_lik_non_spec[spec_id1, spec_id2] <- mat_prob_pop(inds1, inds2, non_list, non_list, c_am)
          }else if(non_mode == "gen_incomp"){
            hybr_lik_non_spec[spec_id1, spec_id2] <- incomp_count_pop(inds1, inds2, non_list, non_list)
          }
          
        }
      }
      
      hybr_lik_non_gens[gen_id] <- mean(hybr_lik_non_spec, na.rm = T) # take the mean across all species pairs    
      
    }
    
    
    
    
    
    
    
  }
  
  c(
    as.character(mean(hybr_lik_host_gens, na.rm = T)),
    as.character(mean(hybr_lik_non_gens, na.rm = T)),
    host_mode,
    non_mode
  )
  
}


try( # try again if it fails
  list.save(hybr_lik, paste(output_dir, '/hybr_lik.rds', sep = '')),
  TRUE
)

# # load the list file
# try( # try again if it fails
#   hybr_lik <- list.load(paste(output_dir, '/hybr_lik.rds', sep = '')),
#   TRUE
# )
# 
# # save the data to a data frame
# hybr_lik <- unlist(hybr_lik)
# hybr_lik <- matrix(hybr_lik, ncol = length(sim_names) * n_rep, nrow = 4)
# hybr_lik <- t(hybr_lik)
# 
# hybr_lik <- data.frame(hybr_lik_host=hybr_lik[,1], hybr_lik_non=hybr_lik[,2], hybr_lik_host_mode=hybr_lik[,3], hybr_lik_non_mode=hybr_lik[,4])
# try( # try again if it fails
#   write.csv(hybr_lik, paste(output_dir, '/hybr_lik.csv', sep = '')),
#   TRUE
# )
```
