## Supplementary material for "Coevolution-induced stabilizing and destabilizing selection shapes species richness in clade co-diversification": File S1: calc_prop_col.R

```
library(rlist)
library(Microsoft365R)
library(scales)

library(future)
library(doParallel)
library(parallel)
n.cores <- future::availableCores()
registerDoParallel(cores=n.cores - 3)

# print the number of slaves
print("The number of slaves is:")
print(n.cores - 3)

sim_nam <- "aa0000FFF" # this can be anything
source('basic_functions.R')

# specify the number of replicates and the total number of generations simulated
n_rep <- 3
n_gen <- 1500

# specify the ages to look at
ages <- c(0:9, 9999)

# specify the size the of landscape
n_dimension <- 7

# generate a vector of the names of the simulations to be plotted
sim_names <- c( # all simulations
  "aa0010AMF",
  "aa0011AMF",
  "aa1010AMF",
  "aa1011AMF",
  "as0010AMF",
  "as0011AMF",
  "as1010AMF",
  "as1011AMF",
  "sa0010AMF",
  "sa0011AMF",
  "sa1010AMF",
  "sa1011AMF",
  "ss0010AMF",
  "ss0011AMF",
  "ss1010AMF",
  "ss1011AMF",
  "aa0110AMM",
  "aa0111AMM",
  "aa1110AMM",
  "aa1111AMM",
  "as0110AMM",
  "as0111AMM",
  "as1110AMM",
  "as1111AMM",
  "sa0110AMM",
  "sa0111AMM",
  "sa1110AMM",
  "sa1111AMM",
  "ss0110AMM",
  "ss0111AMM",
  "ss1110AMM",
  "ss1111AMM",
  "aa0100FFD",
  "aa0101FFD",
  "aa1100FFD",
  "aa1101FFD",
  "as0100FFD",
  "as0101FFD",
  "as1100FFD",
  "as1101FFD",
  "sa0100FFD",
  "sa0101FFD",
  "sa1100FFD",
  "sa1101FFD",
  "ss0100FFD",
  "ss0101FFD",
  "ss1100FFD",
  "ss1101FFD",
  "aa0000FFF",
  "aa0001FFF",
  "aa1000FFF",
  "aa1001FFF",
  "as0000FFF",
  "as0001FFF",
  "as1000FFF",
  "as1001FFF",
  "sa0000FFF",
  "sa0001FFF",
  "sa1000FFF",
  "sa1001FFF",
  "ss0000FFF",
  "ss0001FFF",
  "ss1000FFF",
  "ss1001FFF",
  "aa0100FFM",
  "aa0101FFM",
  "aa1100FFM",
  "aa1101FFM",
  "as0100FFM",
  "as0101FFM",
  "as1100FFM",
  "as1101FFM",
  "sa0100FFM",
  "sa0101FFM",
  "sa1100FFM",
  "sa1101FFM",
  "ss0100FFM",
  "ss0101FFM",
  "ss1100FFM",
  "ss1101FFM",
  "aa0110MDD",
  "aa0111MDD",
  "aa1110MDD",
  "aa1111MDD",
  "as0110MDD",
  "as0111MDD",
  "as1110MDD",
  "as1111MDD",
  "sa0110MDD",
  "sa0111MDD",
  "sa1110MDD",
  "sa1111MDD",
  "ss0110MDD",
  "ss0111MDD",
  "ss1110MDD",
  "ss1111MDD",
  "aa0010MDF",
  "aa0011MDF",
  "aa1010MDF",
  "aa1011MDF",
  "as0010MDF",
  "as0011MDF",
  "as1010MDF",
  "as1011MDF",
  "sa0010MDF",
  "sa0011MDF",
  "sa1010MDF",
  "sa1011MDF",
  "ss0010MDF",
  "ss0011MDF",
  "ss1010MDF",
  "ss1011MDF",
  "aa0010MMF",
  "aa0011MMF",
  "aa1010MMF",
  "aa1011MMF",
  "as0010MMF",
  "as0011MMF",
  "as1010MMF",
  "as1011MMF",
  "sa0010MMF",
  "sa0011MMF",
  "sa1010MMF",
  "sa1011MMF",
  "ss0010MMF",
  "ss0011MMF",
  "ss1010MMF",
  "ss1011MMF",
  "aa0110MMM",
  "aa0111MMM",
  "aa1110MMM",
  "aa1111MMM",
  "as0110MMM",
  "as0111MMM",
  "as1110MMM",
  "as1111MMM",
  "sa0110MMM",
  "sa0111MMM",
  "sa1110MMM",
  "sa1111MMM",
  "ss0110MMM",
  "ss0111MMM",
  "ss1110MMM",
  "ss1111MMM"
)

# specify the generations to look at (the last 100 generations to ensure equilibrium)
#gens <- (n_gen - 10 + 1):n_gen
gens <- 1:n_gen

# create a vector for parallelization
para_vec <- rep(NA, length(sim_names) * n_rep)
for (i in 1:length(sim_names)) {
  for (j in 1:n_rep) {
    para_vec[(i-1) * n_rep + j] <- paste(sim_names[i], as.character(j), sep = '_')
  }
}


# start calculation
od <- NULL
while (length(od) == 0) {
  try( # try again if it fails
    od <- get_business_onedrive(),
    TRUE
  )  
}

prop_col <- foreach (para_id = para_vec) %dopar% {

  sim_id <- substr(para_id, 1, 9)
  rep_id <- substr(para_id, 11, nchar(para_id))
  print(percent((which(para_vec == para_id)-1)/length(para_vec), accuracy = 1))


  # initialize a vector recording the mean range size for the last 10% of all generations
  prop_col_host_gens <- rep(NA, n_gen)
  prop_col_non_gens <- rep(NA, n_gen)
  
  # create a list for storing vectors
  prop_col_host_gens_list <- list(prop_col_host_gens, prop_col_host_gens, prop_col_host_gens, prop_col_host_gens, prop_col_host_gens, prop_col_host_gens, prop_col_host_gens, prop_col_host_gens, prop_col_host_gens, prop_col_host_gens, prop_col_non_gens)
  prop_col_non_gens_list <- list(prop_col_non_gens, prop_col_non_gens, prop_col_non_gens, prop_col_non_gens, prop_col_non_gens, prop_col_non_gens, prop_col_non_gens, prop_col_non_gens, prop_col_non_gens, prop_col_non_gens, prop_col_non_gens)
  
  # read in the generations file
  generations <- NULL
  file_name <- paste(output_dir, '/', sim_id, "/", rep_id, "/generations.rds", sep = '')
  while (length(generations) == 0) {
    try( # try again if it fails
      od$download_file(paste("model_rcpp_hpc_OneDrive/", file_name, sep = ''), file_name, overwrite = T),
      TRUE
    )
    try( # try again if it fails
      generations <- list.load(file_name),
      TRUE
    )    
  }
  unlink(file_name)

  # for each generation
  for(gen_id in gens){

    host_list <- generations[[gen_id]]$host_list
    non_list <- generations[[gen_id]]$non_list
    
    #host_unlist <- unlist(host_list)
    #non_unlist <- unlist(non_list)
    
    for (age in ages) {
      # initialize the colonist and total newborn counts
      sum_col <- 0
      sum_total <- 0
      
      if(age == 9999){
        for (host_id in 1:length(host_list)) {
          #if(host_list[[host_id]]$n_life == age){ # if the individual is a not new born
            sum_total <- sum_total + 1
            if(host_list[[host_id]]$origin == "colonist"){ # if the individuals is also a colonist
              sum_col <- sum_col + 1
            }
          #}
        }
      }else{
        for (host_id in 1:length(host_list)) {
          if(host_list[[host_id]]$n_life == age){ # if the individual is a not new born
            sum_total <- sum_total + 1
            if(host_list[[host_id]]$origin == "colonist"){ # if the individuals is also a colonist
              sum_col <- sum_col + 1
            }
          }
        }
      }
      

      
      prop_col_host_gens_list[[which(ages == age)]][gen_id] <- sum_col / sum_total
      
      # initialize the colonist and total newborn counts
      sum_col <- 0
      sum_total <- 0
      
      if(age == 9999){
        for (non_id in 1:length(non_list)) {
          #if(non_list[[non_id]]$n_life == age){ # if the individual is not a new born
            sum_total <- sum_total + 1
            if(non_list[[non_id]]$origin == "colonist"){ # if the individuals is also a colonist
              sum_col <- sum_col + 1
            }
          #}
        }
      }else{
        for (non_id in 1:length(non_list)) {
          if(non_list[[non_id]]$n_life == age){ # if the individual is not a new born
            sum_total <- sum_total + 1
            if(non_list[[non_id]]$origin == "colonist"){ # if the individuals is also a colonist
              sum_col <- sum_col + 1
            }
          }
        }
      }
      

      
      prop_col_non_gens_list[[which(ages == age)]][gen_id] <- sum_col / sum_total
    }
    
  }
  
  # take the mean across generations for each age to look at
  prop_col_host <- NULL
  prop_col_non <- NULL
  
  for (age in ages) {
    prop_col_host <- c(prop_col_host, mean(prop_col_host_gens_list[[which(ages == age)]], na.rm = T))
    prop_col_non <- c(prop_col_non, mean(prop_col_non_gens_list[[which(ages == age)]], na.rm = T))
  }

  list(
    prop_col_host,
    prop_col_non
  )

}

try( # try again if it fails
  list.save(prop_col, paste(output_dir, '/prop_col.rds', sep = '')),
  TRUE
)
```
