## Supplementary material for "Coevolution-induced stabilizing and destabilizing selection shapes species richness in clade co-diversification": File S1: calc_trait_fluctuation.R

```
library(rlist)
library(Microsoft365R)
library(scales)

library(future)
library(doParallel)
library(parallel)
n.cores <- future::availableCores()
registerDoParallel(cores=n.cores - 8)

# print the number of slaves
print("The number of slaves is:")
print(n.cores - 8)

sim_nam <- "aa0000FFF" # this can be anything
source('basic_functions.R')
source('all_in_one.R')
source('reproduction_pop.R')
source('geographic_dynamics.R')
source('env_optima.R')
source('initialization/common.R')

# specify the number of replicates and the total number of generations simulated
n_rep <- 3
n_gen <- 1500

# specify the size the of landscape
n_dimension <- 7

# generate a vector of the names of the simulations to be plotted
sim_names <- c( # all simulations
  "aa0010AMF",
  "aa0011AMF",
  "aa1010AMF",
  "aa1011AMF",
  "as0010AMF",
  "as0011AMF",
  "as1010AMF",
  "as1011AMF",
  "sa0010AMF",
  "sa0011AMF",
  "sa1010AMF",
  "sa1011AMF",
  "ss0010AMF",
  "ss0011AMF",
  "ss1010AMF",
  "ss1011AMF",
  "aa0110AMM",
  "aa0111AMM",
  "aa1110AMM",
  "aa1111AMM",
  "as0110AMM",
  "as0111AMM",
  "as1110AMM",
  "as1111AMM",
  "sa0110AMM",
  "sa0111AMM",
  "sa1110AMM",
  "sa1111AMM",
  "ss0110AMM",
  "ss0111AMM",
  "ss1110AMM",
  "ss1111AMM",
  "aa0100FFD",
  "aa0101FFD",
  "aa1100FFD",
  "aa1101FFD",
  "as0100FFD",
  "as0101FFD",
  "as1100FFD",
  "as1101FFD",
  "sa0100FFD",
  "sa0101FFD",
  "sa1100FFD",
  "sa1101FFD",
  "ss0100FFD",
  "ss0101FFD",
  "ss1100FFD",
  "ss1101FFD",
  "aa0000FFF",
  "aa0001FFF",
  "aa1000FFF",
  "aa1001FFF",
  "as0000FFF",
  "as0001FFF",
  "as1000FFF",
  "as1001FFF",
  "sa0000FFF",
  "sa0001FFF",
  "sa1000FFF",
  "sa1001FFF",
  "ss0000FFF",
  "ss0001FFF",
  "ss1000FFF",
  "ss1001FFF",
  "aa0100FFM",
  "aa0101FFM",
  "aa1100FFM",
  "aa1101FFM",
  "as0100FFM",
  "as0101FFM",
  "as1100FFM",
  "as1101FFM",
  "sa0100FFM",
  "sa0101FFM",
  "sa1100FFM",
  "sa1101FFM",
  "ss0100FFM",
  "ss0101FFM",
  "ss1100FFM",
  "ss1101FFM",
  "aa0110MDD",
  "aa0111MDD",
  "aa1110MDD",
  "aa1111MDD",
  "as0110MDD",
  "as0111MDD",
  "as1110MDD",
  "as1111MDD",
  "sa0110MDD",
  "sa0111MDD",
  "sa1110MDD",
  "sa1111MDD",
  "ss0110MDD",
  "ss0111MDD",
  "ss1110MDD",
  "ss1111MDD",
  "aa0010MDF",
  "aa0011MDF",
  "aa1010MDF",
  "aa1011MDF",
  "as0010MDF",
  "as0011MDF",
  "as1010MDF",
  "as1011MDF",
  "sa0010MDF",
  "sa0011MDF",
  "sa1010MDF",
  "sa1011MDF",
  "ss0010MDF",
  "ss0011MDF",
  "ss1010MDF",
  "ss1011MDF",
  "aa0010MMF",
  "aa0011MMF",
  "aa1010MMF",
  "aa1011MMF",
  "as0010MMF",
  "as0011MMF",
  "as1010MMF",
  "as1011MMF",
  "sa0010MMF",
  "sa0011MMF",
  "sa1010MMF",
  "sa1011MMF",
  "ss0010MMF",
  "ss0011MMF",
  "ss1010MMF",
  "ss1011MMF",
  "aa0110MMM",
  "aa0111MMM",
  "aa1110MMM",
  "aa1111MMM",
  "as0110MMM",
  "as0111MMM",
  "as1110MMM",
  "as1111MMM",
  "sa0110MMM",
  "sa0111MMM",
  "sa1110MMM",
  "sa1111MMM",
  "ss0110MMM",
  "ss0111MMM",
  "ss1110MMM",
  "ss1111MMM"
)

# create a vector for parallelization
para_vec <- rep(NA, length(sim_names) * n_rep)
for (i in 1:length(sim_names)) {
  for (j in 1:n_rep) {
    para_vec[(i-1) * n_rep + j] <- paste(sim_names[i], as.character(j), sep = '_')
  }
}

od <- NULL
while (length(od) == 0) {
  try( # try again if it fails
    od <- get_business_onedrive(),
    TRUE
  )  
}

delta <- foreach (para_id = para_vec) %dopar% {
  
  sim_id <- substr(para_id, 1, 9)
  rep_id <- substr(para_id, 11, nchar(para_id))
  print(percent((which(para_vec == para_id)-1)/length(para_vec), accuracy = 1))
  
  # initialize a vector recording the mean trait value in the center cell of the grid
  delta_host_gens <- rep(NA, n_gen)
  delta_non_gens <- rep(NA, n_gen)
  
  # initialize a vector recording the standard deviation of trait value in the center cell of the grid
  delta_host_gens_sd <- rep(NA, n_gen)
  delta_non_gens_sd <- rep(NA, n_gen)
  
  # read in the generations file
  generations <- NULL
  file_name <- paste(output_dir, '/', sim_id, "/", rep_id, "/generations.rds", sep = '')
  while (length(generations) == 0) {
    try( # try again if it fails
      od$download_file(paste("model_rcpp_hpc_OneDrive/", file_name, sep = ''), file_name, overwrite = T),
      TRUE
    )
    try( # try again if it fails
      generations <- list.load(file_name),
      TRUE
    )    
  }
  unlink(file_name)
  
  # for each generation

    ########################################################################################
    # for hosts
    for (g in 10:n_gen){ # starting from the 10th generation since the calculation of standard variation is problematic when there is not enough standing variation in trait.
      #print(g)
      global <- rep(NA, length(generations[[g]]$host_list))
      for (i in 1:length(generations[[g]]$host_list)) {
        if(identical(generations[[g]]$host_list[[i]]$site.id, c(ceiling(n_dimension/2),ceiling(n_dimension/2)))){ # focus on the cetral cell only
          global[i] <- get_eco_trait_val(generations[[g]]$host_list[[i]])
        }
      }
      global <- global[!is.na(global)]
      delta_host_gens[g] <- mean(global)
      delta_host_gens_sd[g] <- sd(global)
      
    }
    
  ########################################################################################
  # for nons
  for (g in 10:n_gen){ # starting from the 10th generation since the calculation of standard variation is problematic when there is not enough standing variation in trait.
    #print(g)
    global <- rep(NA, length(generations[[g]]$non_list))
    for (i in 1:length(generations[[g]]$non_list)) {
      if(identical(generations[[g]]$non_list[[i]]$site.id, c(ceiling(n_dimension/2),ceiling(n_dimension/2)))){ # focus on the cetral cell only
        global[i] <- get_eco_trait_val(generations[[g]]$non_list[[i]])
      }
    }
    global <- global[!is.na(global)]
    delta_non_gens[g] <- mean(global)
    delta_non_gens_sd[g] <- sd(global)
    
  } 
  
  temp_host_mean <- abs(delta_host_gens[-1] - delta_host_gens[-length(delta_host_gens)])
  temp_host_sd <- delta_host_gens_sd[-length(delta_host_gens_sd)]
  temp_host <- temp_host_mean[temp_host_sd != 0 & is.na(temp_host_sd) == F]/temp_host_sd[temp_host_sd != 0 & is.na(temp_host_sd) == F]
  
  temp_non_mean <- abs(delta_non_gens[-1] - delta_non_gens[-length(delta_non_gens)])
  temp_non_sd <- delta_non_gens_sd[-length(delta_non_gens_sd)]
  temp_non <- temp_non_mean[temp_non_sd != 0 & is.na(temp_non_sd) == F]/temp_non_sd[temp_non_sd != 0 & is.na(temp_non_sd) == F]
  
  c(
    as.character(mean(    temp_host   , na.rm = T)),
    as.character(mean(    temp_non  , na.rm = T))
  )
  
}


try( # try again if it fails
  list.save(delta, paste(output_dir, '/delta.rds', sep = '')),
  TRUE
)
```
