## Supplementary material for "Coevolution-induced stabilizing and destabilizing selection shapes species richness in clade co-diversification": File S1: control_common.R

```
library(Microsoft365R)
print(sim_nam)

# Default settings
n_sim <- 1000 # number of simulations to run this time
n_succ_run <- 3 # number of "successful" runs to aim at

#initialization
surv <- NULL
expand <- NULL

folder_dir <- paste(output_dir, '/', sim_nam, sep = '')
dir.create(folder_dir)

od <- get_business_onedrive()

folder_list <- od$list_items(paste("model_rcpp_hpc_OneDrive/", output_dir, sep = ''))
folder_list <- folder_list$name
if(!(sim_nam %in% folder_list)){
  od$create_folder(paste("model_rcpp_hpc_OneDrive/", folder_dir, sep = ''))
}


file_name <- paste(folder_dir, '/comp_step.rds', sep = '')
if(file.exists(file_name)){
  comp <- list.load(file_name)
  #n_sim_comp <- comp$rep_comp
}else{
  comp <- as.list(rep(0, 7))# create and initialize a list recording the last step completed
  names(comp) <- c('rep_comp', 'evo', 'sp_del_host', 'sp_del_non', 'phylo_rec_host', 'phylo_rec_non', 'net')
  list.save(comp, file_name)
  #n_sim_comp <- 0
}

source('basic_functions.R')
source('geographic_dynamics.R')
source('env_optima.R')

succ_run <- NULL # create a vector for recording indices of successful runs

for(sim_id in 1:n_sim){
  
  if(comp$rep_comp < n_succ_run){
  
	  sim_output_dir <- paste(folder_dir, "/", as.character(as.numeric(comp$rep_comp) + 1), sep = '')
	  dir.create(sim_output_dir) # create a directory for storing the generated data

	  od <- get_business_onedrive()

	  folder_list <- od$list_items(paste("model_rcpp_hpc_OneDrive/", folder_dir, sep = ''))
	  folder_list <- folder_list$name
	  if(!(as.character(as.numeric(comp$rep_comp) + 1) %in% folder_list)){
	    od$create_folder(paste("model_rcpp_hpc_OneDrive/", sim_output_dir, sep = ''))
	  }
	  
	  print(
		paste("Simulation ", as.character(sim_id), ', ', as.character(length(succ_run)), " successful simulations finished", sep="")
	  )
	  
	  print(
		paste("Successful simulations: ", as.character(succ_run), sep="")
	  )
	  
	  print("Initializing...") 
 source('initialization/common.R')
 
    # 0. initialize whether there is random extinction
	  if(output_dir == "Output_no_extinction"){

	    source('initialization/cat_no.R')

	  }else if(output_dir == "Output_extinction"){

	    source('initialization/cat_ex.R')

	  }else break
    
    
	  # 1. initialize speciation mechanism for hosts

	  source(paste('initialization/host_', as.character(substr(sim_nam, 1, 1)), '.R', sep = ''))

	  if(as.character(substr(sim_nam, 1, 1)) == "a"){

	    source(paste('initialization/1_hetero_0', '.R', sep = ''))

	  }else if(as.character(substr(sim_nam, 1, 1)) == "s"){

	    source(paste('initialization/1_iso_0', '.R', sep = ''))

	  }else break

	

  	  # 2. initialize speciation mechanism for non-hosts

	  source(paste('initialization/non_', as.character(substr(sim_nam, 2, 2)), '.R', sep = ''))

	  if(as.character(substr(sim_nam, 2, 2)) == "a"){

	    source(paste('initialization/4_hetero_0', '.R', sep = ''))

	  }else if(as.character(substr(sim_nam, 2, 2)) == "s"){

	    source(paste('initialization/4_iso_0', '.R', sep = ''))

	  }else break

	

  	  # 3. initialize 1

    if(as.character(substr(sim_nam, 1, 1)) == "a"){

	    source(paste('initialization/1_iso_', as.character(substr(sim_nam, 3, 3)), '.R', sep = ''))

	  }else if(as.character(substr(sim_nam, 1, 1)) == "s"){

	    source(paste('initialization/1_hetero_', as.character(substr(sim_nam, 3, 3)), '.R', sep = ''))

	  }else break

	

  	  # 4. initialize 2

	  source(paste('initialization/2_', as.character(substr(sim_nam, 4, 4)), '.R', sep = ''))

	

  	  # 5. initialize 3

	  source(paste('initialization/3_', as.character(substr(sim_nam, 5, 5)), '.R', sep = ''))

	

  	  # 6. initialize 4

	  if(as.character(substr(sim_nam, 2, 2)) == "a"){

	    source(paste('initialization/4_iso_', as.character(substr(sim_nam, 6, 6)), '.R', sep = ''))

	  }else if(as.character(substr(sim_nam, 2, 2)) == "s"){

	    source(paste('initialization/4_hetero_', as.character(substr(sim_nam, 6, 6)), '.R', sep = ''))

	  }else break

	

  	  # 7. initialize what all treatments have in common

	  #source('initialization/common.R')
	  source('reproduction_combined_smart_det_repr.R')
	  
	  print("Evolving...")
	  if(length(succ_run) == 0 && comp$evo == 1){
	  }else{
	    source('events_generation_reordered_det_death_finite_lifespan.R')
	  }
	  
	  
		if(identical(surv, "NO") || identical(expand, "NO")){
		}else{	
		  comp$evo <- 1
		  file_name <- paste(folder_dir, '/comp_step.rds', sep = '')
		  list.save(comp, file_name)
		  
			source('reproduction_pop.R')
		  
		  print("Delineating host species...")
		  if(length(succ_run) == 0 && comp$sp_del_host == 1){
		  }else{
			  source('species_delineation_parallelization_memory_future_host.R')
		  }
		  comp$sp_del_host <- 1
		  file_name <- paste(folder_dir, '/comp_step.rds', sep = '')
		  list.save(comp, file_name)
		  
		  print("Delineating non species...")
		  if(length(succ_run) == 0 && comp$sp_del_non == 1){
		  }else{
		    source('species_delineation_parallelization_memory_future_non.R')
		  }
		  comp$sp_del_non <- 1
		  file_name <- paste(folder_dir, '/comp_step.rds', sep = '')
		  list.save(comp, file_name)
				  
			print("Phylogenetic reconstruction for hosts")
			if(length(succ_run) == 0 && comp$phylo_rec_host == 1){
			}else{
			  #source('phylogenetic_reconstruction_parallelization_host.R')
			}
			comp$phylo_rec_host <- 1
			file_name <- paste(folder_dir, '/comp_step.rds', sep = '')
			list.save(comp, file_name)
			
			print("Phylogenetic reconstruction for nons")
			if(length(succ_run) == 0 && comp$phylo_rec_non == 1){
			}else{
			  #source('phylogenetic_reconstruction_parallelization_non.R')
			}
			comp$phylo_rec_non <- 1
			file_name <- paste(folder_dir, '/comp_step.rds', sep = '')
			list.save(comp, file_name)
				  
			print("Building networks...")
			if(length(succ_run) == 0 && comp$net == 1){
			}else{
			  #source('network_construction_parallelization.R')
			  networks <- as.list(1)
			  list.save(networks, paste(sim_output_dir, "/networks.rds", sep = ''))
			}
			comp$net <- 1
			file_name <- paste(folder_dir, '/comp_step.rds', sep = '')
			list.save(comp, file_name)
			
			succ_run <- c(succ_run, sim_id)
			
			comp_temp<- comp # modify comp after a complete run
			comp_temp$rep_comp <- comp_temp$rep_comp + 1
			comp_temp$evo <- 0
			comp_temp$sp_del_host <- 0
			comp_temp$sp_del_non <- 0
			comp_temp$phylo_rec_host <- 0
			comp_temp$phylo_rec_non <- 0
			comp_temp$net <- 0
			comp <- comp_temp
			file_name <- paste(folder_dir, '/comp_step.rds', sep = '')
			list.save(comp, file_name)
			
			# do this if there is a shortage of storage on the HPC
			unlink( paste(folder_dir, "/", as.character(as.numeric(comp$rep_comp)), "/generations.rds", sep = '') )
			
		}
		
  }
  

}
```
