## Supplementary material for "Coevolution-induced stabilizing and destabilizing selection shapes species richness in clade co-diversification": File S1: env_optima.R

```
# Generate a random distribution of environmental optima (normally distributed)

env_optima <- function(n_dimension, mu, sigma){
  
  grid <- matrix(data = rnorm(n = n_dimension^2, mean = mu, sd = sigma), nrow = n_dimension, ncol = n_dimension)
  
  return(grid)
  
}
```
