## Supplementary material for "Coevolution-induced stabilizing and destabilizing selection shapes species richness in clade co-diversification": File S1: events_generation_reordered_det_death_finite_lifespan.R

```
library(Microsoft365R)

# for testing
# n_gen <- 100

#################################################################
# initialize the vector for recording the ranking for colonists
col_rank_host_mean <- rep(NA, n_gen)
col_rank_host_max <- rep(NA, n_gen)
col_rank_host_min <- rep(NA, n_gen)

col_rank_non_mean <- rep(NA, n_gen)
col_rank_non_max <- rep(NA, n_gen)
col_rank_non_min <- rep(NA, n_gen)

# initialize the vector for recording the proportion of colonists
col_prop_host <- rep(NA, n_gen)
col_prop_non <- rep(NA, n_gen)

#################################################################

#################################################################
# Record the initial states of both lists and both grids
generation <- list(host_list, non_list, grid_host, grid_non, grid.bar, NA, NA, grid.opt.host, grid.opt.non)
names(generation) <- c('host_list', 'non_list', 'grid_host', 'grid_non', 'grid.bar', 'cat.row', 'cat.col', 'grid.opt.host', 'grid.opt.non')
generations <- as.list(rep(NA, n_gen))
generations[[1]] <- generation

print(c(sim_id, 1, length(host_list), length(non_list)))

surv <- "YES" # recording whether the hosts and non-hosts survived
expand <- "NO" # recording whether the hosts and non_hosts expand

for (i in 2:n_gen){
  
  gen_id <- i
  
  # initialize the matrices for recording colonist ranking for each site/host
  col_rank_host_mean_mat <- matrix(NA, ncol = n_dimension, nrow = n_dimension)
  col_rank_host_max_mat <- matrix(NA, ncol = n_dimension, nrow = n_dimension)
  col_rank_host_min_mat <- matrix(NA, ncol = n_dimension, nrow = n_dimension)
  
  col_rank_non_mean_mat <- NULL
  col_rank_non_max_mat <- NULL
  col_rank_non_min_mat <- NULL
  
  #col_rank_non_mean_mat <- matrix(NA, ncol = n_dimension, nrow = n_dimension)
  #col_rank_non_max_mat <- matrix(NA, ncol = n_dimension, nrow = n_dimension)
  #col_rank_non_min_mat <- matrix(NA, ncol = n_dimension, nrow = n_dimension)
  
  # initialize the matrices for recording colonist proportion for each site/host
  col_prop_host_mat <- matrix(NA, ncol = n_dimension, nrow = n_dimension)
  col_prop_non_mat <- NULL
  #col_prop_non_mat <- matrix(NA, ncol = n_dimension, nrow = n_dimension)
  
  # Collect garbage before each gen if there is a shortage of memory
  gc()
    
  # record the lengths of host and non-host lists
  length_host <- length(host_list)
  length_non <- length(non_list)
  
  # testing
  #grid_non_old <- grid_non
  

  # increase the age of each host by 1
  for (life in 1:length(host_list)) {
    if(length(host_list[[life]]) == 8){
      host_list[[life]]$n_life <- host_list[[life]]$n_life + 1
    }
  }
  
  # increase the age of each non-host by 1
  for (life in 1:length(non_list)) {
    if(length(non_list[[life]]) == 8){
      non_list[[life]]$n_life <- non_list[[life]]$n_life + 1
    }
  }
  
##########################################################3333  
  # for each cell in the grid
      #host reproduction and movement
      for (j in 1:n_dimension){
        for (k in 1:n_dimension){
          
          site <- as.numeric(c(j, k))
          
          temp_list <- host_repr(grid_host, host_list, non_list, site, w_0_host, c_mat_host, n_offspr_host, host_choose_thres, host_comp_thres, host_no_comp_loci, parent_origin, host_mu, host_bar * grid.bar, n_dimension, sigma_eco_host, sigma_pre_host)
          host_list <- temp_list[[1]]
          host_no_comp_loci <- temp_list[[2]]
          parent_origin <- temp_list[[3]]
          
        }
      }
      grid_host <- count_host_grid(host_list, n_dimension)
  
      #non-host reproduction and movement
      for (j in 1:n_dimension){
        for (k in 1:n_dimension){
          
          site <- as.numeric(c(j, k))
          
          temp_list <- non_repr(grid_non, non_list, grid_host, host_list, site, w_0_non, c_mat_non, n_offspr_non, non_choose_thres, non_comp_thres, non_no_comp_loci, parent_origin, non_mu, non_bar * grid.bar, n_dimension, sigma_eco_non, sigma_pre_non)
          non_list <- temp_list[[1]]
          non_no_comp_loci <- temp_list[[2]]
          parent_origin <- temp_list[[3]]
                    
        }
      }
      grid_non <- count_non_grid(non_list, n_dimension)
      grid_host <- count_host_grid(host_list, n_dimension)
      host_list <- count_non_host(host_list, non_list)

      

      
  # death of hosts due to lifespan limit
  for(host_life in 1:length(host_list)){
    if(length(host_list[[host_life]]) == 8 && host_list[[host_life]]$n_life >= n_life_host){
      host_list[[host_life]] <- as.list("D_life")
    }
  }
  
  grid_host <- count_host_grid(host_list, n_dimension) #update grid_non
      
##############################################################
  # competitive death of hosts, with an environment capacity of K_host
  for (j in 1:n_dimension){
    for (k in 1:n_dimension){
      
      #find all host IDs in the current site
      host.on.site.comp <- NULL
      for (host_id in 1:length(host_list)){
        if(length(host_list[[host_id]]) == 8){ #ensure alive
          if(identical(as.numeric(host_list[[host_id]]$site.id), as.numeric(c(j,k)))){
            host.on.site.comp <- c(host.on.site.comp, host_id)
          }
        }
      }
      
      #create a vector containing fitness for every host individual in the host list
      host.fit.comp <- rep(w_0_host, length(host_list))
      
      # # calculate fitness for each host in this site molded by coevolution
      for (non_id in 1:length(non_list)) {
        if(length(non_list[[non_id]]) == 8 && identical(as.numeric(non_list[[non_id]]$site.id), as.numeric(c(j,k)))){
          if(length(host_list[[non_list[[non_id]]$host.id]]) == 8){
            host.fit.comp[non_list[[non_id]]$host.id] <- host.fit.comp[non_list[[non_id]]$host.id] + fit_func_host(zi = get_eco_trait_val(host_list[[non_list[[non_id]]$host.id]]), zj = get_eco_trait_val(non_list[[non_id]]), xi = xi_host, alpha = alpha_host)            
          }
        }
      }
      
      # each host's fitness is further molded by selection by the environment. Note: ladder is a parameter to help circumvent R's lower limit for numeric type
      # for (host_env in host.on.site.comp) {
      #   z_star <- grid.opt.host[host_list[[host_env]]$site.id[1], host_list[[host_env]]$site.id[2]]
      #   host.fit.comp[host_env] <- host.fit.comp[host_env] * exp(-gamma_host * (get_eco_trait_val(host_list[[host_env]]) - z_star)^2)
      # }
      host.fit.comp <- host.fit.comp[host.on.site.comp]
      
      
      ###########################################################################################################################
      # record the average percentile (ranking) of fitness for all colonists on a site
      # record whether each individual is a local or a colonist
      if(length(host.on.site.comp) > 0){
        
        loc_col <- rep("local", length(host.on.site.comp)) # each individual is a local by default
        for (loc_col_id in 1:length(host.on.site.comp)) {
          if(host_list[[host.on.site.comp[loc_col_id]]]$origin == "colonist"){
            loc_col[loc_col_id] <- "colonist"
          }
        }
        
        
        if(sum(loc_col == "colonist") > 0){ # if there are more than one colonist on the site
          
           # calculate the ranking (percentile) of each individual, with higher percentile indicating higher fitness
          ranking_all <- 1 - (rank(host.fit.comp) / length(host.fit.comp))
          
          ranking_colonist <- ranking_all[loc_col == "colonist"]
          
          # take the mean and record it
          col_rank_host_mean_mat[j,k] <- mean(ranking_colonist)
          col_rank_host_max_mat[j,k] <- max(ranking_colonist)
          col_rank_host_min_mat[j,k] <- min(ranking_colonist)         
          
        }
        
        # calculate the proportion of colonists
        prop_colonist <- sum(loc_col == "colonist") / length(host.on.site.comp)
        
        # record the proportion of colonists
        col_prop_host_mat[j,k] <- prop_colonist
        
      }
      

      ###########################################################################################################################
      
      
      while (grid_host[j,k] > K_host){
        death.id <- host.on.site.comp[host.fit.comp==min(host.fit.comp)]
        if(length(death.id) > 1){
          death.id <- sample(host.on.site.comp[host.fit.comp==min(host.fit.comp)], 1) # the individual with the lowest fitness dies
        }
        host_list[[death.id]] <- as.list("D_comp") # update host_list
        grid_host[j,k] <- grid_host[j,k] - 1 # update grid_host
        subset <- (host.on.site.comp != death.id)
        host.on.site.comp <- host.on.site.comp[subset]
        host.fit.comp <- host.fit.comp[subset]
      }
      
    }
  }
      

  col_rank_host_mean[i] <- mean(col_rank_host_mean_mat, na.rm = T)    # take the mean across the matrix and store the mean
  col_rank_host_max[i] <- mean(col_rank_host_max_mat, na.rm = T)
  col_rank_host_min[i] <- mean(col_rank_host_min_mat, na.rm = T)
  
  col_prop_host[i] <- mean(col_prop_host_mat, na.rm = T)
      
      
      
      
      
      
      
      
      
  # death of hosts due to reproduction
  # for(host_aging in 1:length(host_list)){
  #   if(length(host_list[[host_aging]]) == 8 && host_list[[host_aging]]$n_repr >= n_repr_host){
  #     host_list[[host_aging]] <- as.list("D_repr")
  #   }
  # }
  
  #kill all non-hosts on the dead hosts, too
  for (l in 1:length(non_list)) {
    if (length(non_list[[l]]) == 8){
      if (length(host_list[[non_list[[l]]$host.id]]) < 8){
        non_list[[l]] <- as.list("D_host")
      }
    }
  }
      
  grid_non <- count_non_grid(non_list, n_dimension) #update grid_non
  grid_host <- count_host_grid(host_list, n_dimension) #update grid_non
  

  
  
  
  
  
    
  # death of non-hosts due to lifespan limit
  for(non_life in 1:length(non_list)){
    if(length(non_list[[non_life]]) == 8 && non_list[[non_life]]$n_life >= n_life_non){
      non_list[[non_life]] <- as.list("D_life")
    }
  }
  
  grid_non <- count_non_grid(non_list, n_dimension) #update grid_non
  host_list <- count_non_host(host_list, non_list) # update host_list
  
  # competitive death of non-hosts, with a host capacity of K-non
  for (m in 1:length(host_list)) {
    if(length(host_list[[m]]$n_non)>0){ #only look at hosts that are still alive


      #find all non-host IDs on this host
      non.on.host <- NULL
      for (non_id in 1:length(non_list)){
        if(length(non_list[[non_id]]) == 8){ #ensure alive
          if(identical(as.numeric(non_list[[non_id]]$host.id), as.numeric(m)) == T){
            non.on.host <- c(non.on.host, non_id)
          }
        }

      }


      #create a vector containing fitness for every non-host individual in the non-host list
      non.fit.comp <- rep(w_0_non, length(non_list))

      # # calculate fitness for each non-host on this host molded by coevolution
      for (non_id in 1:length(non_list)) {
        if(length(non_list[[non_id]]) == 8 && identical(as.numeric(non_list[[non_id]]$host.id), as.numeric(m))){
          non.fit.comp[non_id] <- non.fit.comp[non_id] + fit_func_non(zi = get_eco_trait_val(non_list[[non_id]]), zj = get_eco_trait_val(host_list[[m]]), xi = xi_non, alpha = alpha_non)
        }
      }

      # each non-host's fitness is further molded by selection by the environment.
      # for (non_env in non.on.host) {
      #   z_star <- grid.opt.non[non_list[[non_env]]$site.id[1], non_list[[non_env]]$site.id[2]]
      #   non.fit.comp[non_env] <- non.fit.comp[non_env] * exp(-gamma_non * (get_eco_trait_val(non_list[[non_env]]) - z_star)^2)
      # }
      non.fit.comp <- non.fit.comp[non.on.host]


      ###########################################################################################################################
      # record the average percentile (ranking) of fitness for all colonists on a site
      # record whether each individual is a local or a colonist
      if(length(non.on.host) > 0){

        loc_col <- rep("local", length(non.on.host)) # each individual is a local by default
        for (loc_col_id in 1:length(non.on.host)) {
          if(non_list[[non.on.host[loc_col_id]]]$origin == "colonist"){
            loc_col[loc_col_id] <- "colonist"
          }
        }

        if(sum(loc_col == "colonist") > 0){ # if there are more than one colonist on the host

          # calculate the ranking (percentile) of each individual, with higher percentile indicating higher fitness
          ranking_all <- 1 - (rank(non.fit.comp) / length(non.fit.comp))

          ranking_colonist <- ranking_all[loc_col == "colonist"]

          # take the mean and record it
          col_rank_non_mean_mat <- c(col_rank_non_mean_mat, mean(ranking_colonist))
          col_rank_non_max_mat <- c(col_rank_non_max_mat, max(ranking_colonist))
          col_rank_non_min_mat <- c(col_rank_non_min_mat, min(ranking_colonist))


        }

        # calculate the proportion of colonists
        prop_colonist <- sum(loc_col == "colonist") / length(non.on.host)

        # record the proportion of colonists
        col_prop_non_mat <- c(col_prop_non_mat, prop_colonist)

      }


      ###########################################################################################################################


      while(host_list[[m]]$n_non > K_non){
        death.id <- non.on.host[non.fit.comp==min(non.fit.comp)]
        if(length(death.id) > 1){
          death.id <- sample(non.on.host[non.fit.comp==min(non.fit.comp)], 1) # the individual with the lowest fitness dies
        }
        non_list[[death.id]] <- as.list("D_comp")
        host_list[[m]]$n_non <- host_list[[m]]$n_non - 1 #update host_list
        subset <- (non.on.host != death.id)
        non.on.host <- non.on.host[subset]
        non.fit.comp <- non.fit.comp[subset]
      }
    }
  }

  col_rank_non_mean[i] <- mean(col_rank_non_mean_mat, na.rm = T)    # take the mean across the matrix and store the mean
  col_rank_non_max[i] <- mean(col_rank_non_max_mat, na.rm = T)
  col_rank_non_min[i] <- mean(col_rank_non_min_mat, na.rm = T)

  col_prop_non[i] <- mean(col_prop_non_mat, na.rm = T)
  
  ##############################################################  
  # competitive death of nons, with an environment capacity of K_non
  # for (j in 1:n_dimension){
  #   for (k in 1:n_dimension){
  #     
  #     #find all non IDs in the current site
  #     non.on.site.comp <- NULL
  #     for (non_id in 1:length(non_list)){
  #       if(length(non_list[[non_id]]) == 8){ #ensure alive
  #         if(identical(as.numeric(non_list[[non_id]]$site.id), as.numeric(c(j,k)))){
  #           non.on.site.comp <- c(non.on.site.comp, non_id)
  #         }
  #       }
  #     }
  #     
  #     #create a vector containing fitness for every non individual in the non list
  #     non.fit.comp <- rep(w_0_non, length(non_list))
  #     
  #     # # calculate fitness for each non in this site molded by coevolution
  #     for (non_id in 1:length(non_list)) {
  #       if(length(non_list[[non_id]]) == 8 && identical(as.numeric(non_list[[non_id]]$site.id), as.numeric(c(j,k)))){
  #         non.fit.comp[non_id] <- non.fit.comp[non_id] + fit_func_non(zi = get_eco_trait_val(non_list[[non_id]]), zj = get_eco_trait_val(host_list[[non_list[[non_id]]$host.id]]), xi = xi_non, alpha = alpha_non)            
  #       }
  #     }
  #     
  #     # each non's fitness is further molded by selection by the environment. Note: ladder is a parameter to help circumvent R's lower limit for numeric type
  #     # for (non_env in non.on.site.comp) {
  #     #   z_star <- grid.opt.non[non_list[[non_env]]$site.id[1], non_list[[non_env]]$site.id[2]]
  #     #   non.fit.comp[non_env] <- non.fit.comp[non_env] * exp(-gamma_non * (get_eco_trait_val(non_list[[non_env]]) - z_star)^2)
  #     # }
  #     non.fit.comp <- non.fit.comp[non.on.site.comp]
  #     
  #     
  #     ###########################################################################################################################
  #     # record the average percentile (ranking) of fitness for all colonists on a site
  #     # record whether each individual is a local or a colonist
  #     if(length(non.on.site.comp) > 0){
  #       
  #       loc_col <- rep("local", length(non.on.site.comp)) # each individual is a local by default
  #       for (loc_col_id in 1:length(non.on.site.comp)) {
  #         if(non_list[[non.on.site.comp[loc_col_id]]]$origin == "colonist"){
  #           loc_col[loc_col_id] <- "colonist"
  #         }
  #       }
  #       
  #       
  #       if(sum(loc_col == "colonist") > 0){ # if there are more than one colonist on the site
  #         
  #         # calculate the ranking (percentile) of each individual, with higher percentile indicating higher fitness
  #         ranking_all <- 1 - (rank(non.fit.comp) / length(non.fit.comp))
  #         
  #         ranking_colonist <- ranking_all[loc_col == "colonist"]
  #         
  #         # take the mean and record it
  #         col_rank_non_mean_mat[j,k] <- mean(ranking_colonist)
  #         col_rank_non_max_mat[j,k] <- max(ranking_colonist)
  #         col_rank_non_min_mat[j,k] <- min(ranking_colonist)         
  #         
  #       }
  #       
  #       # calculate the proportion of colonists
  #       prop_colonist <- sum(loc_col == "colonist") / length(non.on.site.comp)
  #       
  #       # record the proportion of colonists
  #       col_prop_non_mat[j,k] <- prop_colonist
  #       
  #     }
  #     
  #     
  #     ###########################################################################################################################
  #     
  #     
  #     while (grid_non[j,k] > K_non){
  #       death.id <- non.on.site.comp[non.fit.comp==min(non.fit.comp)]
  #       if(length(death.id) > 1){
  #         death.id <- sample(non.on.site.comp[non.fit.comp==min(non.fit.comp)], 1) # the individual with the lowest fitness dies
  #       }
  #       non_list[[death.id]] <- as.list("D_comp") # update non_list
  #       grid_non[j,k] <- grid_non[j,k] - 1 # update grid_non
  #       subset <- (non.on.site.comp != death.id)
  #       non.on.site.comp <- non.on.site.comp[subset]
  #       non.fit.comp <- non.fit.comp[subset]
  #     }
  #     
  #   }
  # }
  # 
  # 
  # col_rank_non_mean[i] <- mean(col_rank_non_mean_mat, na.rm = T)    # take the mean across the matrix and store the mean
  # col_rank_non_max[i] <- mean(col_rank_non_max_mat, na.rm = T)
  # col_rank_non_min[i] <- mean(col_rank_non_min_mat, na.rm = T)
  # 
  # col_prop_non[i] <- mean(col_prop_non_mat, na.rm = T)
  
  
  
  
  
  
  
  
  
  
  
  
  
  
  
  
  
  
  
  
  
  # death of non-hosts due to reproduction
  # for(non_aging in 1:length(non_list)){
  #   if(length(non_list[[non_aging]]) == 8 && non_list[[non_aging]]$n_repr >= n_repr_non){
  #     non_list[[non_aging]] <- as.list("D_repr")
  #   }
  # }
  

  grid_non <- count_non_grid(non_list, n_dimension) #update grid_non
  #print("yes")
  host_list <- count_non_host(host_list, non_list) # update host_list
  
  
################################################################3  
  # catastrophe that kills all individuals in that site
  cat.row <- NA
  cat.col <- NA
  
  # no_cat <- 1 # number of catastrophes is fixed
  # no_cat <- rpois(1, prob_cat) # number of catastrophes determined by a poisson distribution
  
  if(no_cat > 0){
    # randomly choose a site
    cat.cell <- as.numeric(sample(1:(n_dimension^2), size=no_cat, replace = F)) 
    
    cat.row <- floor((cat.cell-1)/n_dimension) + 1
    cat.col <- cat.cell %% n_dimension
    cat.col[cat.col == 0] <- 7
    
    for (cat in 1: no_cat) {
  
      for(host.no in 1:length(host_list)){
        if(identical(host_list[[host.no]]$site.id[1],cat.row[cat]) && identical(host_list[[host.no]]$site.id[2],cat.col[cat])){
          host_list[[host.no]] <- as.list("D_cat")
          #print("Die!")
        }
      }
      
      for(non.no in 1:length(non_list)){
        if(identical(non_list[[non.no]]$site.id[1],cat.row[cat]) && identical(non_list[[non.no]]$site.id[2], cat.col[cat])){
          non_list[[non.no]] <- as.list("D_cat")
          #print("Die!")
        }
      }
      
    }
    
    # update both grids after catastrophes
    grid_host <- count_host_grid(host_list, n_dimension)
    grid_non <- count_non_grid(non_list, n_dimension)    
  }


##################################################################3  
  # record ancestry of all living individuals (position in the list of hosts/non-hosts from the previous generation)
  #for(n in 1:length(host_list)){
  #  if(length(host_list[[n]]) == 8){
  #    host_list[[n]]$anc.id <- 0
  #  }
  #}
  for(n in 1:length(host_list)){
    if(length(host_list[[n]]) == 8){
      host_list[[n]]$anc.id <- n
    }    
  }
  #######
  #for(o in 1:length(non_list)){
  #  if(length(non_list[[o]]) == 8){
  #    non_list[[o]]$anc.id <- 0
  #  }
  #}
  
  for(o in 1:length(non_list)){
    if(length(non_list[[o]]) == 8){
      non_list[[o]]$anc.id <- o
    }    
  }
  
  # remove dead individuals  
  host_list.new <- NULL
  for (p in 1:length(host_list)) {
    if(length(host_list[[p]]) == 8){
        host_list.new[[length(host_list.new) + 1]] <- host_list[[p]]
    }
  }
  
  non_list.new <- NULL
  for (q in 1:length(non_list)) {
    if(length(non_list[[q]]) == 8){
        non_list.new[[length(non_list.new) + 1]] <- non_list[[q]]
    }
  }
  
  # terminate evolution if hosts or non-hosts go extinct
  if(length(host_list.new) < 2 || length(non_list.new) < 2){
	surv <- "NO"
	print("Hosts or non-hosts went extinct. Evolution terminated.")
	break
  }
  
  # create a vector containing the anc.id of all living hosts (their host.id from last generation)
  host_list.new.anc <- NULL
  for (r in 1:length(host_list.new)) {
    host_list.new.anc <- c(host_list.new.anc, host_list.new[[r]]$anc.id)
  }
  
  # update host.id of each non-host
  for (s in 1:length(non_list.new)){
    old.id <- non_list.new[[s]]$host.id
    new.id <- which(host_list.new.anc == old.id)
    non_list.new[[s]]$host.id <- new.id
  }
  
  #testing
  #print(non_list.new)
  
  # replace old host and non-host lists with new ones
  host_list <- host_list.new
  non_list <- non_list.new
  
  #testing
  print(c(sim_id, i, length(host_list), length(non_list)))
   
  #print(sum(grid_non - grid_non_old))
  #print(grid_non - grid_non_old)
  
  # store the states of both grids and both lists
  generation <- list(host_list, non_list, grid_host, grid_non, grid.bar, cat.row, cat.col)
  names(generation) <- c('host_list', 'non_list', 'grid_host', 'grid_non', 'grid.bar', 'cat.row', 'cat.col')
  generations[[i]] <- generation
  
  
  # store the the ranking for colonists for both the host and the non-host
  
  
   
  # geographic barriers rise and fall

  # if(i < n_gen_conn){
  #   grid.bar <- grid.bar_0
  #   #print(1)
  # }
  # else{
  #   grid.bar <- grid.bar_1
  #   #print(0)
  # }
  
  if(i < 0){
    grid.bar <- grid.bar_0
  }
  else{
      if((i-1)%%per_geo < n_gen_conn){
       grid.bar <- grid.bar_0
       #print(1)
      }
      else if((i-1)%%per_geo >= n_gen_conn && i%%per_geo < per_geo){
       grid.bar <- grid.bar_1
       #print(0)
      }      
  }
  


  
}


# save the colonist ranking and proportion to a data frame
colonist <- data.frame(col_rank_host_mean, col_rank_host_max, col_rank_host_min, col_prop_host, col_rank_non_mean, col_rank_non_max, col_rank_non_min, col_prop_non)
file_name <- paste(sim_output_dir, "/colonist.csv", sep = '')
write.csv(colonist, file_name)


# save the parent origin data frame
file_name <- paste(sim_output_dir, "/parent_origin.csv", sep = '')
write.csv(parent_origin, file_name)

# plot for intuition
# plot(1:n_gen, col_ranking0000$col_prop_host, type = "line", ylim = c(0,1))
# lines(1:n_gen, col_ranking1001$col_prop_host, col = "red")
# 
# plot(1:n_gen, col_ranking0000$col_rank_host_mean, type = "line", ylim = c(0,1))
# lines(1:n_gen, col_ranking1001$col_rank_host_mean, col = "red")
# 
# plot(1:n_gen, col_ranking0000$col_rank_host_max, type = "line", ylim = c(0,1))
# lines(1:n_gen, col_ranking1001$col_rank_host_max, col = "red")
# 
# plot(1:n_gen, col_ranking0000$col_rank_host_min, type = "line", ylim = c(0,1))
# lines(1:n_gen, col_ranking1001$col_rank_host_min, col = "red")


# plot(1:n_gen, col_ranking0000$col_prop_non, type = "line", ylim = c(0,1))
# lines(1:n_gen, col_ranking1001$col_prop_non, col = "red")
# 
# plot(1:n_gen, col_ranking0000$col_rank_non_mean, type = "line", ylim = c(0,1))
# lines(1:n_gen, col_ranking1001$col_rank_non_mean, col = "red")
# 
# plot(1:n_gen, col_ranking0000$col_rank_non_max, type = "line", ylim = c(0,1))
# lines(1:n_gen, col_ranking1001$col_rank_non_max, col = "red")
# 
# plot(1:n_gen, col_ranking0000$col_rank_non_min, type = "line", ylim = c(0,1))
# lines(1:n_gen, col_ranking1001$col_rank_non_min, col = "red")

# determine whether both hosts and non-hosts have expanded
if(!is.na(generations[[10]])){
    if(length(generations[[10]]$host_list)>10){
        expand <- "YES"
    }    
}

# if survided and expanded, save generations file
if(expand == "YES" && surv == "YES"){
  
  # save generations to a file
  file_name <- paste(sim_output_dir, "/generations.rds", sep = '')
  list.save(generations, file_name)
  
  # upload generations.rds to OneDrive and delete the local file
  od <- NULL
  while (length(od) == 0) {
    try( # try again if it fails
      od <- get_business_onedrive(),
      TRUE
    )  
  }
  
  while (!as.logical(sum(od$list_items(paste("model_rcpp_hpc_OneDrive/", sim_output_dir, sep = '')) == "generations.rds"))) { # if this file is not yet stored in OneDrive
    try( # try again if it fails
      od$upload_file(file_name, dest = paste("model_rcpp_hpc_OneDrive/", file_name, sep = '')),
      TRUE
    )  
  }
  
  unlink(file_name)
  
}


# read in the saved file
# generations <- list.load('Output/generations.rds')

if(surv == "YES"){
	# test if the hosts have become choosier in general
	choosiness_gens_host <- rep(NA, length(generations))

	for (g in 1:length(generations)){
	  #print(g)
	  choosiness <- rep(NA, length(generations[[g]]$host_list))
	  for (i in 1:length(generations[[g]]$host_list)) {
		choosiness[i] <- get_choos_trait_val(generations[[g]]$host_list[[i]])
	  }
	  choosiness_gens_host[g] <- mean(choosiness)
	  
	}  

	file_name <- paste(sim_output_dir, "/host_choosiness.pdf", sep = '')
	par(mar=c(2,2,2,2))
	pdf(file = file_name)
	plot(1:length(generations), choosiness_gens_host, main = "host choosiness", type = "S")
	dev.off()

	# test if the non_hosts have become choosier in general
	choosiness_gens_non <- rep(NA, length(generations))

	for (g in 1:length(generations)){
	  #print(g)
	  choosiness <- rep(NA, length(generations[[g]]$non_list))
	  for (i in 1:length(generations[[g]]$non_list)) {
		choosiness[i] <- get_choos_trait_val(generations[[g]]$non_list[[i]])
	  }
	  choosiness_gens_non[g] <- mean(choosiness)
	  
	}  

	file_name <- paste(sim_output_dir, "/non_choosiness.pdf", sep = '')
	par(mar=c(2,2,2,2))
	pdf(file = file_name)
	plot(1:length(generations), choosiness_gens_non, main = "non-host choosiness", type = "S")
	dev.off()

}

# testing
#for (test in 1:length(non_list)){
#  print(c(test, non_list[[test]]$host.id))
#}
```
