## Supplementary material for "Coevolution-induced stabilizing and destabilizing selection shapes species richness in clade co-diversification": File S1: figure_continuous.R

```
# create a new variable for the square of Delta_H and Delta_D
# dat_na_rm$Delta_H2 <- dat_na_rm$Delta_H ^ 2
# dat_na_rm$Delta_D2 <- dat_na_rm$Delta_D ^ 2

# logit transform the proportion data
# logitTransform <- function(p) { log(p/(1-p)) }
# dat_na_rm$Proportion_of_locals_among_host_individuals <- logitTransform(dat_na_rm$Proportion_of_locals_among_host_individuals)
# dat_na_rm$Proportion_of_locals_among_dependent_individuals <- logitTransform(dat_na_rm$Proportion_of_locals_among_dependent_individuals)

theme_set(
  theme_classic() +
    theme(legend.position = "right")
)

# remove outliers
#host_max <- max(dat_na_rm$Delta_H)
#dat_na_rm <- dat_na_rm[dat_na_rm$Delta_H < host_max,]

# calculate the 95% interval for delta values for the neutral scenario
# t_test_host <- t.test(dat_na_rm$Delta_H[dat_na_rm$Scenario=='i'], conf.level = 0.999999999999999)
# host_min <- t_test_host$conf.int[1]
# host_max <- t_test_host$conf.int[2]
host_min <- min(dat_na_rm$Delta_H[dat_na_rm$Scenario=='i'])
host_max <- max(dat_na_rm$Delta_H[dat_na_rm$Scenario=='i'])

# t_test_non <- t.test(dat_na_rm$Delta_D[dat_na_rm$Scenario=='i'], conf.level = 0.999999999999999)
# non_min <- t_test_non$conf.int[1]
# non_max <- t_test_non$conf.int[2]
non_min <- min(dat_na_rm$Delta_D[dat_na_rm$Scenario=='i'])
non_max <- max(dat_na_rm$Delta_D[dat_na_rm$Scenario=='i'])

# For hosts
# plot the proportion of local individuals against delta
p1 <- ggplot(dat_na_rm, aes(x=Delta_H, y=Proportion_of_locals_among_host_individuals)) +
  geom_text(x=host_min-0.01, y=0.228, label="Stabilizing", aes(angle=90), size=4)+
  geom_text(x=(host_min + host_max)/2, y=0.228, label="Neutral", aes(angle=90), size=4)+
  geom_text(x=host_max+0.01, y=0.228, label="Destabilizing", aes(angle=90), size=4)+
  geom_point(aes(color=Scenario, shape=Scenario))+
  geom_smooth(aes(colour = NA), colour = 'black', method=loess, se=T, fullrange=TRUE)+
  #geom_smooth(data=subset(dat_na_rm, dat_na_rm$Delta_H<host_max), aes(colour = NA), colour = 'blue', linetype='dashed', method = "lm", se=F) +
  #geom_smooth(data=subset(dat_na_rm, dat_na_rm$Delta_H>host_min & dat_na_rm$Delta_H<host_max), aes(colour = NA), colour = 'black', method = "lm", se=F) +
  #geom_smooth(data=subset(dat_na_rm, dat_na_rm$Delta_H>host_min), aes(colour = NA), colour = 'red', linetype='dashed', method = "lm", se=F)+
  #geom_smooth(aes(colour = NA), method=lm, se=T, fullrange=TRUE)+
  geom_vline(xintercept=host_min, linetype='dashed', color='black', linewidth=0.5)+
  geom_vline(xintercept=host_max, linetype='dashed', color='black', linewidth=0.5)+
  xlab(expression(bar('|'*italic(Delta * z )*'|')))+
  ylab('Degree of dispersal limitation')+
  ggtitle('(D)')+
  scale_shape_manual(values=c(0, 1, 2, 3, 4, 5, 6, 7, 8))+
  theme(text = element_text(size = 15), plot.title=element_text(size=27))+
  theme(legend.text = element_text(face = "italic"))

# linear_left <- lm(Proportion_of_locals_among_host_individuals~Delta_H, data = dat_na_rm[dat_na_rm$Delta_H<host_max,])
# summary(linear_left)
# AIC(linear_left)
# 
# qua_left <- lm(Proportion_of_locals_among_host_individuals~Delta_H+Delta_H2, data = dat_na_rm[dat_na_rm$Delta_H<host_max,])
# summary(qua_left)
# AIC(qua_left)
# 
# linear_right <- lm(Proportion_of_locals_among_host_individuals~Delta_H, data = dat_na_rm[dat_na_rm$Delta_H>host_min,])
# summary(linear_right)
# AIC(linear_right)
# 
# qua_right <- lm(Proportion_of_locals_among_host_individuals~Delta_H+Delta_H2, data = dat_na_rm[dat_na_rm$Delta_H>host_min,])
# summary(qua_right)
# AIC(qua_right)

p4 <- ggplot(dat_na_rm, aes(x=Delta_D, y=Proportion_of_locals_among_dependent_individuals)) +
  geom_text(x=non_min-0.01, y=0.2425, label="Stabilizing", aes(angle=90), size=4)+
  geom_text(x=(non_min + non_max)/2, y=0.2425, label="Neutral", aes(angle=90), size=4)+
  geom_text(x=non_max+0.01, y=0.2425, label="Destabilizing", aes(angle=90), size=4)+
  geom_point(aes(color=Scenario, shape=Scenario))+
  geom_smooth(aes(colour = NA), colour = 'black', method=loess, se=T, fullrange=TRUE)+
  #geom_smooth(data=subset(dat_na_rm, dat_na_rm$Delta_D<non_max), aes(colour = NA), colour = 'black', method = "lm", se=F) +
  #geom_smooth(data=subset(dat_na_rm, dat_na_rm$Delta_D>non_min & dat_na_rm$Delta_D<non_max), aes(colour = NA), colour = 'black', method = "lm", se=F) +
  #geom_smooth(data=subset(dat_na_rm, dat_na_rm$Delta_D>non_min), aes(colour = NA), colour = 'black', method = "lm", se=F)+
  #geom_smooth(aes(colour = NA), method=lm, se=T, fullrange=TRUE)+
  geom_vline(xintercept=non_min, linetype='dashed', color='black', linewidth=0.5)+
  geom_vline(xintercept=non_max, linetype='dashed', color='black', linewidth=0.5)+
  xlab(expression(bar('|'*italic(Delta * z )*'|')))+
  ylab('Degree of dispersal limitation')+
  ggtitle('(G)')+
  scale_shape_manual(values=c(0, 1, 2, 3, 4, 5, 6, 7, 8))+
  theme(text = element_text(size = 15), plot.title=element_text(size=27))+
  theme(legend.text = element_text(face = "italic"))

# slope_left <- lm(Proportion_of_locals_among_dependent_individuals~Delta_D, data = dat_na_rm[dat_na_rm$Delta_D<non_max,])
# slope_right <- lm(Proportion_of_locals_among_dependent_individuals~Delta_D, data = dat_na_rm[dat_na_rm$Delta_D>non_min,])
# summary(slope_left)
# summary(slope_right)

p2 <- ggplot(dat_na_rm, aes(x=Delta_H, y=Difficulty_of_hybridization_H)) +
  geom_text(x=host_min-0.01, y=3.8, label="Stabilizing", aes(angle=90), size=4)+
  geom_text(x=(host_min + host_max)/2, y=3.8, label="Neutral", aes(angle=90), size=4)+
  geom_text(x=host_max+0.01, y=3.8, label="Destabilizing", aes(angle=90), size=4)+
  geom_point(aes(color=Scenario, shape=Scenario))+
  geom_smooth(aes(colour = NA), colour = 'black', method=loess, se=T, fullrange=TRUE)+
  #geom_smooth(data=subset(dat_na_rm, dat_na_rm$Delta_H<host_max), aes(colour = NA), colour = 'black', method = "lm", se=F) +
  #geom_smooth(data=subset(dat_na_rm, dat_na_rm$Delta_H>host_min & dat_na_rm$Delta_H<host_max), aes(colour = NA), colour = 'black', method = "lm", se=F) +
  #geom_smooth(data=subset(dat_na_rm, dat_na_rm$Delta_H>host_min), aes(colour = NA), colour = 'black', method = "lm", se=F)+
  #geom_smooth(aes(colour = NA), method=lm, se=T, fullrange=TRUE)+
  geom_vline(xintercept=host_min, linetype='dashed', color='black', linewidth=0.5)+
  geom_vline(xintercept=host_max, linetype='dashed', color='black', linewidth=0.5)+
  xlab(expression(bar('|'*italic(Delta * z )*'|')))+
  ylab('Genetic distance between sites')+
  ggtitle('(E)')+
  scale_shape_manual(values=c(0, 1, 2, 3, 4, 5, 6, 7, 8))+
  theme(text = element_text(size = 15), plot.title=element_text(size=27))+
  theme(legend.text = element_text(face = "italic"))

# slope_left <- lm(Difficulty_of_hybridization_H~Delta_H, data = dat_na_rm[dat_na_rm$Delta_H<host_max,])
# slope_right <- lm(Difficulty_of_hybridization_H~Delta_H, data = dat_na_rm[dat_na_rm$Delta_H>host_min,])
# summary(slope_left)
# summary(slope_right)

p5 <- ggplot(dat_na_rm, aes(x=Delta_D, y=Difficulty_of_hybridization_D)) +
  geom_text(x=non_min-0.01, y=3, label="Stabilizing", aes(angle=90), size=4)+
  geom_text(x=(non_min + non_max)/2, y=3, label="Neutral", aes(angle=90), size=4)+
  geom_text(x=non_max+0.01, y=3, label="Destabilizing", aes(angle=90), size=4)+geom_point(aes(color=Scenario, shape=Scenario))+
  geom_point(aes(color=Scenario, shape=Scenario))+
  geom_smooth(aes(colour = NA), colour = 'black', method=loess, se=T, fullrange=TRUE)+
  #geom_smooth(data=subset(dat_na_rm, dat_na_rm$Delta_D<non_max), aes(colour = NA), colour = 'black', method = "lm", se=F) +
  #geom_smooth(data=subset(dat_na_rm, dat_na_rm$Delta_D>non_min & dat_na_rm$Delta_D<non_max), aes(colour = NA), colour = 'black', method = "lm", se=F) +
  #geom_smooth(data=subset(dat_na_rm, dat_na_rm$Delta_D>non_min), aes(colour = NA), colour = 'black', method = "lm", se=F)+
  #geom_smooth(aes(colour = NA), method=lm, se=T, fullrange=TRUE)+
  geom_vline(xintercept=non_min, linetype='dashed', color='black', linewidth=0.5)+
  geom_vline(xintercept=non_max, linetype='dashed', color='black', linewidth=0.5)+
  xlab(expression(bar('|'*italic(Delta * z )*'|')))+
  ylab('Genetic distance between sites')+
  ggtitle('(H)')+
  scale_shape_manual(values=c(0, 1, 2, 3, 4, 5, 6, 7, 8))+
  theme(text = element_text(size = 15), plot.title=element_text(size=27))+
  theme(legend.text = element_text(face = "italic"))

# slope_left <- lm(Difficulty_of_hybridization_D~Delta_D, data = dat_na_rm[dat_na_rm$Delta_D<non_max,])
# slope_right <- lm(Difficulty_of_hybridization_D~Delta_D, data = dat_na_rm[dat_na_rm$Delta_D>non_min,])
# summary(slope_left)
# summary(slope_right)

p3 <- ggplot(dat_na_rm, aes(x=Delta_H, y=Clade_diversity_H)) +
  geom_text(x=host_min-0.01, y=4, label="Stabilizing", aes(angle=90), size=4)+
  geom_text(x=(host_min + host_max)/2, y=4, label="Neutral", aes(angle=90), size=4)+
  geom_text(x=host_max+0.01, y=4, label="Destabilizing", aes(angle=90), size=4)+
  geom_point(aes(color=Scenario, shape=Scenario))+
  geom_smooth(aes(colour = NA), colour = 'black', method=loess, se=T, fullrange=TRUE)+
  #geom_smooth(data=subset(dat_na_rm, dat_na_rm$Delta_H<host_max), aes(colour = NA), colour = 'black', method = "lm", se=F) +
  #geom_smooth(data=subset(dat_na_rm, dat_na_rm$Delta_H>host_min & dat_na_rm$Delta_H<host_max), aes(colour = NA), colour = 'black', method = "lm", se=F) +
  #geom_smooth(data=subset(dat_na_rm, dat_na_rm$Delta_H>host_min), aes(colour = NA), colour = 'black', method = "lm", se=F)+
  #geom_smooth(aes(colour = NA), method=lm, se=T, fullrange=TRUE)+
  geom_vline(xintercept=host_min, linetype='dashed', color='black', linewidth=0.5)+
  geom_vline(xintercept=host_max, linetype='dashed', color='black', linewidth=0.5)+
  xlab(expression(bar('|'*italic(Delta * z )*'|')))+
  ylab('Species richness accumulated')+
  ggtitle('(F)')+
  scale_shape_manual(values=c(0, 1, 2, 3, 4, 5, 6, 7, 8))+
  theme(text = element_text(size = 15), plot.title=element_text(size=27))+
  theme(legend.text = element_text(face = "italic"))

# slope_left <- lm(Clade_diversity_H~Delta_H, data = dat_na_rm[dat_na_rm$Delta_H<host_max,])
# slope_right <- lm(Clade_diversity_H~Delta_H, data = dat_na_rm[dat_na_rm$Delta_H>host_min,])
# summary(slope_left)
# summary(slope_right)

p6 <- ggplot(dat_na_rm, aes(x=Delta_D, y=Clade_diversity_D)) +
  geom_text(x=non_min-0.01, y=22, label="Stabilizing", aes(angle=90), size=4)+
  geom_text(x=(non_min + non_max)/2, y=22, label="Neutral", aes(angle=90), size=4)+
  geom_text(x=non_max+0.01, y=22, label="Destabilizing", aes(angle=90), size=4)+geom_point(aes(color=Scenario, shape=Scenario))+
  geom_point(aes(color=Scenario, shape=Scenario))+
  geom_smooth(aes(colour = NA), colour = 'black', method=loess, se=T, fullrange=TRUE)+
  #geom_smooth(data=subset(dat_na_rm, dat_na_rm$Delta_D<non_max), aes(colour = NA), colour = 'black', method = "lm", se=F) +
  #geom_smooth(data=subset(dat_na_rm, dat_na_rm$Delta_D>non_min & dat_na_rm$Delta_D<non_max), aes(colour = NA), colour = 'black', method = "lm", se=F) +
  #geom_smooth(data=subset(dat_na_rm, dat_na_rm$Delta_D>non_min), aes(colour = NA), colour = 'black', method = "lm", se=F)+
  #geom_smooth(aes(colour = NA), method=lm, se=T, fullrange=TRUE)+
  geom_vline(xintercept=non_min, linetype='dashed', color='black', linewidth=0.5)+
  geom_vline(xintercept=non_max, linetype='dashed', color='black', linewidth=0.5)+
  xlab(expression(bar('|'*italic(Delta * z )*'|')))+
  ylab('Species richness accumulated')+
  ggtitle('(I)')+
  scale_shape_manual(values=c(0, 1, 2, 3, 4, 5, 6, 7, 8))+
  theme(text = element_text(size = 15), plot.title=element_text(size=27))+
  theme(legend.text = element_text(face = "italic"))

# slope_left <- lm(Clade_diversity_D~Delta_D, data = dat_na_rm[dat_na_rm$Delta_D<non_max,])
# slope_right <- lm(Clade_diversity_D~Delta_D, data = dat_na_rm[dat_na_rm$Delta_D>non_min,])
# summary(slope_left)
# summary(slope_right)

combined <- ggpubr::ggarrange(
  p1, p2, p3,
  p4, p5, p6,
  ncol = 3, nrow = 2
)

pdf("figure_continuous_1001.pdf", width = 14, height = 9)
combined
dev.off()
```
