## Supplementary material for "Coevolution-induced stabilizing and destabilizing selection shapes species richness in clade co-diversification": File S1: geographic_dynamics.R

```
# 1. Rise and fall of geographic barriers

# function that generate a random landscape with barriers
# matrix representing cells (geographic sites)

gen_lands <- function(n_dimension, prob_b){
  
  # create a grid representing cells (sites)
  grid = matrix(0, nrow = n_dimension, ncol = n_dimension)
  
  # add vertical barriers
  for (i in 1:(2*n_dimension - 1)){
    if (i==1)
      grid.bar.temp = as.matrix(grid[,(i+1)/2])
    if (i>1 && i%%2==0)
      grid.bar.temp = cbind(grid.bar.temp, matrix(sample(c(0.1, 99.9), size = n_dimension, prob = c(1-prob_b, prob_b), replace = T), nrow = n_dimension, ncol = 1))
    if (i>1 && i%%2!=0)
      grid.bar.temp = cbind(grid.bar.temp, as.matrix(grid[,(i+1)/2]))
  }
  
  # add horizontal barriers
  for (i in 1:(2*n_dimension - 1)){
    if (i==1)
      grid.bar = t(as.matrix(grid.bar.temp[(i+1)/2,]))
    if (i>1 && i%%2==0)
      grid.bar = rbind(grid.bar, matrix(sample(c(0.1, 99.9), size = 2*n_dimension - 1, prob = c(1-prob_b, prob_b), replace = T), nrow = 1, ncol = 2*n_dimension - 1))
    if (i>1 && i%%2!=0)
      grid.bar = rbind(grid.bar, t(as.matrix(grid.bar.temp[(i+1)/2,])))
  }

  return(grid.bar)
  
}


# 2. Find sets of connected sites, i.e., patches
connectivity.matrix <- function(grid.bar){
  
  #initialize connectivity matrix
  conn.mat <- matrix(0, nrow = n_dimension, ncol = n_dimension)
  
  #initialize the number of sets of connected sites
  no.set <- 0
  
  #initialize connectedness to above and left
  above <- "NO"
  left <- "NO"
  
  for (i in 1:nrow(conn.mat)) {
    for (j in 1:ncol(conn.mat)) {
      
      if(i==1 && j==1){
        conn.mat[i,j] <- no.set + 1
        no.set <- no.set + 1
      }
      else{
        
        #first check if the cell is connected to the cell above
        if(i==1 || identical(grid.bar[2*i-2, 2*j-1], 99.9) == T){
          above <- "NO"
        }
        else{
          above <- "YES"
        }
        
        #then check is the cell is connected to the cell to the left
        if(j==1 || identical(grid.bar[2*i-1, 2*j-2], 99.9) == T){
          left <- "NO"
        }
        else{
          left <- "YES"
        }
        
        # if connected to both above and left
        if(above=="YES" && left=="YES"){
          conn.mat[i,j] <- conn.mat[i,j-1] # same as left
          conn.mat[which(conn.mat == conn.mat[i-1,j])] <- conn.mat[i,j-1]
        }
        
        # if connected to above, but not left
        else if(above=="YES" && left=="NO"){
          conn.mat[i,j] <- conn.mat[i-1,j]
        }
        
        # if not connected to above, but to left
        else if(above=="NO" && left=="YES"){
          conn.mat[i,j] <- conn.mat[i,j-1]
        }
        
        # if not connected to either above or left
        else if(above=="NO" && left=="NO"){
          conn.mat[i,j] <- no.set + 1
          no.set <- no.set + 1
        }
        
      }
      
    }
  }
  
  # (optional) renumber patch IDs
  old.pat.ids <- unique(as.vector(conn.mat))
  new.pat.ids <- 1:length(old.pat.ids)
  
  conn.mat.vec <- as.vector(conn.mat)
  
  for (i in 1:length(old.pat.ids)) {
    conn.mat.vec[conn.mat.vec == old.pat.ids[i]] <- -new.pat.ids[i]
  }
  
  conn.mat.vec <- -conn.mat.vec
  conn.mat <- matrix(conn.mat.vec, nrow = n_dimension, ncol = n_dimension)
  
  return(conn.mat)
  
}
```
