## Supplementary material for "Coevolution-induced stabilizing and destabilizing selection shapes species richness in clade co-diversification": File S1: common.R

```
# 0. Create grid

## 0.1 Generate a random landscape
n_dimension <- 7
prob_b <- 0#1

grid.bar_1 <- gen_lands(n_dimension, prob_b) # landscape with barriers
grid.bar_0 <- gen_lands(n_dimension, 0) # landscape without barriers

grid.bar <- grid.bar_1
#plot(grid.bar)


## 0.2 Generate a random distribution of environmental optima
grid.opt.host <- env_optima(n_dimension, mu = 2, sigma = 20) # the mean can be determined by the mean trait value at initiation
grid.opt.non <- env_optima(n_dimension, mu = 2, sigma = 20) # the mean can be determined by the mean trait value at initiation

# 1. Create individuals

## 1.1 Basic genetic modules
eco_locus_unit = c(1,1)
choos_locus_unit = c(0.25, 0.25)
comp_locus_unit = c(1,1)

# Ecological trait loci
L_z = 1
loci_eco = rep(list(eco_locus_unit), L_z)

# Prezygotic isolation loci
L_a = 1
loci_pre = rep(list(choos_locus_unit), L_a)

# Postzygotic isolation loci
L_b = 5
loci_post = rep(list(comp_locus_unit), L_b)

## 1.2 Initial site
n = n_dimension
site_init = c(round((n+1)/2), round((n+1)/2))

## 1.3 Host
### 1.3.1 Individual host
host_basic = list(loci_eco,
                  loci_pre,
                  loci_post,
                  0,
                  site_init,
                  0,
                  #0,
                  0,
                  "local")
names(host_basic) = c('eco',
                      'pre',
                      'post',
                      'anc.id',
                      'site.id',
                      'n_non',
                      #'n_repr', # this is commented out because the number of successful reproductions is no longer of concern
                      'n_life',
                      'origin')

### 1.3.2 A list of hosts
n_host_init = 7
host_list = rep(list(host_basic), n_host_init)

## 1.4 Non-host
### 1.4.1 Individual non-host
host_init = 1
non_basic = list(loci_eco,
                 loci_pre,
                 loci_post,
                 0,
                 site_init,
                 host_init,
                 #0,
                 0,
                 'local')
names(non_basic) = c('eco',
                     'pre',
                     'post',
                     'anc.id',
                     'site.id',
                     'host.id',
                     #'n_repr', # this is commented out because the number of successful reproductions is no longer of concern
                     'n_life',
                     'origin')

### 1.4.2 A list of non-hosts
n_non_per_host <- 1
n_non_init = n_host_init * n_non_per_host
non_list = rep(list(non_basic), n_non_init)
for(i in 1:n_non_init){
  non_list[[i]]$host.id <- floor((i-1)/n_non_per_host) + 1
}

host_list <- count_non_host(host_list, non_list)

### 1.4.3 Update the grid
grid_host <- count_host_grid(host_list, n_dimension)
grid_non <- count_non_grid(non_list, n_dimension)


# 2. Parameters for reproduction and movement

### Initializing mutations rates for eco and pre alleles
sigma_eco_non <- 0.2
#sigma_eco_non <- 0
sigma_eco_host <- 0.2

sigma_pre_non <- 0
sigma_pre_host <- 0

### Initializing number of compatibility alleles for each loci so far
host_no_comp_loci <- rep(1, length(host_basic$post))
non_no_comp_loci <- rep(1, length(non_basic$post))

### Initializing mutation rate of incompatibility alleles
host_mu <- 0.03
non_mu <- 0.03

### Number of lifetime reproductions
n_repr_host <- Inf
n_repr_non <- Inf

### Number of maximum lifespan
#n_life_host <- Inf
n_life_host <- Inf
#n_life_host <- 15
#n_life_non <- Inf
n_life_non <- Inf
#n_life_non <- 10

# the probability of dispersal
p_dis_host <- 0.8
p_dis_non <- 0.8

# 3. Parameters for evolution

### Number of generations
n_gen <- 1500#1000

# absolute geographic barrier
n_gen_iso <- 0#6 # number of generations during which barriers are present
n_gen_conn <- 7#1 # # number of generations during which barriers are absent
per_geo <- n_gen_iso + n_gen_conn

c_am <- 10 # scaling constant in the mating probability function (Equation 3 in PDF)

# base fitness for hosts and non_hosts (fitness with species interactions)
w_0_host <- 0.01 #0.2000000000000001 #0.01
w_0_non <- 0.01 #0.0000000000000001 #0.01

# number of matings per cell
c_mat_host <- 1#3
c_mat_non <- 9

# number of offspring per successful mating
n_offspr_host <- 2
n_offspr_non <- 6

# environment and host capacity
K_host <- 10
K_non <- 2
#K_non <- 20

# 4. Species delineation

### Initializing the lists of host and non-host population and species
host_spec_list <- list(1)
host_pop_list <- list(1)#list(1:10)

non_spec_list <- list(1)
non_pop_list <- list(1)#list(1:10)

# minimum between-individual phenotypic difference for individuals to be considered from different populations
host.max.phen.dist <- Inf
non.max.phen.dist <- Inf

# initialize a data frame for recording whether the parents of offspring are locals or colonists
host_loc_loc <- rep(0, n_gen)
host_col_col <- rep(0, n_gen)
host_loc_col <- rep(0, n_gen)
non_loc_loc <- rep(0, n_gen)
non_col_col <- rep(0, n_gen)
non_loc_col <- rep(0, n_gen)

parent_origin <- data.frame(
  host_loc_loc,
  host_col_col,
  host_loc_col,
  non_loc_loc,
  non_col_col,
  non_loc_col
)
```
