## Supplementary material for "Coevolution-induced stabilizing and destabilizing selection shapes species richness in clade co-diversification": File S1: read_in_data.R

```
library(rlist)
library(tidyverse)
#install.packages("factoextra")
library(factoextra)
#install.packages("corrplot")
library(corrplot)
library(ggplot2)
library(ggpubr)
#install.packages("ggthemes")
library(ggthemes)
#install.packages("egg")
library(egg)
library(Rarity)

#install.packages("car") #An extremely useful/in-depth regression package
#install.packages("stargazer") #Produces easy to read regression results (similar to what you get in SPSS)
#install.packages("effects") #We will use this to create our interactions
#install.packages("ggplot2") #Our incredibly powerful and versatile graphing package
library(car)
library(stargazer)
library(effects)

#install.packages("interactions")
library(interactions)

# specify the number of replicates
n_rep <- 3

# generate a vector of the names of the simulations to be plotted
sim_names <- c( # all simulations
  "aa0010AMF",
  "aa0011AMF",
  "aa1010AMF",
  "aa1011AMF",
  "as0010AMF",
  "as0011AMF",
  "as1010AMF",
  "as1011AMF",
  "sa0010AMF",
  "sa0011AMF",
  "sa1010AMF",
  "sa1011AMF",
  "ss0010AMF",
  "ss0011AMF",
  "ss1010AMF",
  "ss1011AMF",
  "aa0110AMM",
  "aa0111AMM",
  "aa1110AMM",
  "aa1111AMM",
  "as0110AMM",
  "as0111AMM",
  "as1110AMM",
  "as1111AMM",
  "sa0110AMM",
  "sa0111AMM",
  "sa1110AMM",
  "sa1111AMM",
  "ss0110AMM",
  "ss0111AMM",
  "ss1110AMM",
  "ss1111AMM",
  "aa0100FFD",
  "aa0101FFD",
  "aa1100FFD",
  "aa1101FFD",
  "as0100FFD",
  "as0101FFD",
  "as1100FFD",
  "as1101FFD",
  "sa0100FFD",
  "sa0101FFD",
  "sa1100FFD",
  "sa1101FFD",
  "ss0100FFD",
  "ss0101FFD",
  "ss1100FFD",
  "ss1101FFD",
  "aa0000FFF",
  "aa0001FFF",
  "aa1000FFF",
  "aa1001FFF",
  "as0000FFF",
  "as0001FFF",
  "as1000FFF",
  "as1001FFF",
  "sa0000FFF",
  "sa0001FFF",
  "sa1000FFF",
  "sa1001FFF",
  "ss0000FFF",
  "ss0001FFF",
  "ss1000FFF",
  "ss1001FFF",
  "aa0100FFM",
  "aa0101FFM",
  "aa1100FFM",
  "aa1101FFM",
  "as0100FFM",
  "as0101FFM",
  "as1100FFM",
  "as1101FFM",
  "sa0100FFM",
  "sa0101FFM",
  "sa1100FFM",
  "sa1101FFM",
  "ss0100FFM",
  "ss0101FFM",
  "ss1100FFM",
  "ss1101FFM",
  "aa0110MDD",
  "aa0111MDD",
  "aa1110MDD",
  "aa1111MDD",
  "as0110MDD",
  "as0111MDD",
  "as1110MDD",
  "as1111MDD",
  "sa0110MDD",
  "sa0111MDD",
  "sa1110MDD",
  "sa1111MDD",
  "ss0110MDD",
  "ss0111MDD",
  "ss1110MDD",
  "ss1111MDD",
  "aa0010MDF",
  "aa0011MDF",
  "aa1010MDF",
  "aa1011MDF",
  "as0010MDF",
  "as0011MDF",
  "as1010MDF",
  "as1011MDF",
  "sa0010MDF",
  "sa0011MDF",
  "sa1010MDF",
  "sa1011MDF",
  "ss0010MDF",
  "ss0011MDF",
  "ss1010MDF",
  "ss1011MDF",
  "aa0010MMF",
  "aa0011MMF",
  "aa1010MMF",
  "aa1011MMF",
  "as0010MMF",
  "as0011MMF",
  "as1010MMF",
  "as1011MMF",
  "sa0010MMF",
  "sa0011MMF",
  "sa1010MMF",
  "sa1011MMF",
  "ss0010MMF",
  "ss0011MMF",
  "ss1010MMF",
  "ss1011MMF",
  "aa0110MMM",
  "aa0111MMM",
  "aa1110MMM",
  "aa1111MMM",
  "as0110MMM",
  "as0111MMM",
  "as1110MMM",
  "as1111MMM",
  "sa0110MMM",
  "sa0111MMM",
  "sa1110MMM",
  "sa1111MMM",
  "ss0110MMM",
  "ss0111MMM",
  "ss1110MMM",
  "ss1111MMM"
)

# create a vector recording the combination of independent variables for each similation
# note: no_extinction comes first, followed by extinction ones
nam_temp <- rep(NA, length(sim_names) * n_rep)
for ( rep_id in 1:n_rep) {
  nam_temp[(1:length(sim_names)) * n_rep - (n_rep - rep_id)] <- sim_names
}

nam <- c(
  paste(nam_temp, "no", sep = '_'),
  paste(nam_temp, "ex", sep = '_')
)


# read in the intermediate variables
# colonization
# prop_col_no <- read.csv("Output_no_extinction/prop_col.csv", header = T)
# prop_col_ex <- read.csv("Output_extinction/prop_col.csv", header = T)
# prop_col <- rbind(prop_col_no, prop_col_ex)
# 
# prop_col_host <- prop_col$prop_col_host
# prop_col_dep <- prop_col$prop_col_non

temp_list_no <- list.load("Output_no_extinction/prop_col.rds")
temp_list_ex <- list.load("Output_extinction/prop_col.rds")
temp_list <- c(temp_list_no, temp_list_ex)

# temp_list_no <- list.load("Output_no_extinction/prop_newborn.rds")
# temp_list_ex <- list.load("Output_extinction/prop_newborn.rds")
# temp_list <- c(temp_list_no, temp_list_ex)

prop_col_host <- rep(NA, length(temp_list))
prop_col_dep <- rep(NA, length(temp_list))

for (i in 1:length(prop_col_host)){
  prop_col_host[i] <- temp_list[[i]][[1]][[11]]
}

for (i in 1:length(prop_col_dep)){
  prop_col_dep[i] <- temp_list[[i]][[2]][[11]]
}


# age
# age_no <- read.csv("Output_no_extinction/age.csv", header = T)
# age_ex <- read.csv("Output_extinction/age.csv", header = T)
# age <- rbind(age_no, age_ex)
# 
# age_host <- age$age_host
# age_dep <- age$age_non

# # trait variation
# trait_var_no <- read.csv("Output_no_extinction/trait_var.csv", header = T)
# trait_var_ex <- read.csv("Output_extinction/trait_var.csv", header = T)
# trait_var <- rbind(trait_var_no, trait_var_ex)
# 
# trait_var_host <- trait_var$trait_var_host
# trait_var_dep <- trait_var$trait_var_non


#rang_siz_no <- read.csv("Output_no_extinction/rang_siz.csv", header = T)
#rang_siz_ex <- read.csv("Output_extinction/rang_siz.csv", header = T)
#rang_siz <- rbind(rang_siz_no, rang_siz_ex)

#rang_siz_mean_host <- rang_siz$rang_siz_host
#rang_siz_mean_dep <- rang_siz$rang_siz_non

#rang_siz_sum_host <- rang_siz$rang_siz_host_sum
#rang_siz_sum_dep <- rang_siz$rang_siz_non_sum

# demographic stochasticity
#dem_stoch_no <- read.csv("Output_no_extinction/demo_stoch.csv", header = T)
#dem_stoch_ex <- read.csv("Output_extinction/demo_stoch.csv", header = T)
#dem_stoch <- rbind(dem_stoch_no, dem_stoch_ex)

#dem_stoch_mean_host <- dem_stoch$demo_stoch_host
#dem_stoch_mean_dep <- dem_stoch$demo_stoch_non

# hybridization likelihood
# hybr_lik_no <- read.csv("Output_no_extinction/hybr_lik.csv", header = T)
# hybr_lik_ex <- read.csv("Output_extinction/hybr_lik.csv", header = T)
# hybr_lik <- rbind(hybr_lik_no, hybr_lik_ex)
# 
# hybr_lik_mean_host <- hybr_lik$hybr_lik_host
# hybr_lik_mean_dep <- hybr_lik$hybr_lik_non

temp_list_no <- list.load("Output_no_extinction/hybr_lik.rds")
temp_list_ex <- list.load("Output_extinction/hybr_lik.rds")
temp_list <- c(temp_list_no, temp_list_ex)

# temp_list_no <- list.load("Output_no_extinction/prop_newborn.rds")
# temp_list_ex <- list.load("Output_extinction/prop_newborn.rds")
# temp_list <- c(temp_list_no, temp_list_ex)

hybr_lik_mean_host <- rep(NA, length(temp_list))
hybr_lik_mean_dep <- rep(NA, length(temp_list))

for (i in 1:length(hybr_lik_mean_host)){
  hybr_lik_mean_host[i] <- temp_list[[i]][[1]]
}

for (i in 1:length(prop_col_dep)){
  hybr_lik_mean_dep[i] <- temp_list[[i]][[2]]
}


# trait temporl variation


temp_list_no <- list.load("Output_no_extinction/delta.rds")
temp_list_ex <- list.load("Output_extinction/delta.rds")
temp_list <- c(temp_list_no, temp_list_ex)

# temp_list_no <- list.load("Output_no_extinction/prop_newborn.rds")
# temp_list_ex <- list.load("Output_extinction/prop_newborn.rds")
# temp_list <- c(temp_list_no, temp_list_ex)

delta_host <- rep(NA, length(temp_list))
delta_dep <- rep(NA, length(temp_list))

for (i in 1:length(delta_host)){
  delta_host[i] <- temp_list[[i]][[1]]
}

for (i in 1:length(delta_dep)){
  delta_dep[i] <- temp_list[[i]][[2]]
}


# range or partner shift
# rang_part_shift_no <- read.csv("Output_no_extinction/rang_part_shift.csv", header = T)
# rang_part_shift_ex <- read.csv("Output_extinction/rang_part_shift.csv", header = T)
# rang_part_shift <- rbind(rang_part_shift_no, rang_part_shift_ex)
# 
# ana_rang_siz_bef_host <- rang_part_shift$ana_rang_siz_bef_host
# ana_rang_siz_aft_host <- rang_part_shift$ana_rang_siz_aft_host
# ana_rang_siz_prop_host <- rang_part_shift$ana_rang_siz_prop_host
# ana_rang_dist_gain_host <- rang_part_shift$ana_rang_dist_gain_host
# ana_rang_dist_loss_host <- rang_part_shift$ana_rang_dist_loss_host
# 
# clado_rang_siz_bef_host <- rang_part_shift$clado_rang_siz_bef_host
# clado_rang_siz_aft_host <- rang_part_shift$clado_rang_siz_aft_host
# clado_rang_siz_prop_host <- rang_part_shift$clado_rang_siz_prop_host
# clado_rang_dist_gain_host <- rang_part_shift$clado_rang_dist_gain_host
# clado_rang_dist_loss_host <- rang_part_shift$clado_rang_dist_loss_host
# 
# comb_rang_siz_bef_host <- rang_part_shift$comb_rang_siz_bef_host
# comb_rang_siz_aft_host <- rang_part_shift$comb_rang_siz_aft_host
# comb_rang_siz_prop_host <- rang_part_shift$comb_rang_siz_prop_host
# comb_rang_dist_gain_host <- rang_part_shift$comb_rang_dist_gain_host
# comb_rang_dist_loss_host <- rang_part_shift$comb_rang_dist_loss_host
# 
# ana_part_siz_bef_host <- rang_part_shift$ana_part_siz_bef_host
# ana_part_siz_aft_host <- rang_part_shift$ana_part_siz_aft_host
# ana_part_siz_prop_host <- rang_part_shift$ana_part_siz_prop_host
# ana_part_dist_gain_host <- rang_part_shift$ana_part_dist_gain_host
# ana_part_dist_loss_host <- rang_part_shift$ana_part_dist_loss_host
# 
# clado_part_siz_bef_host <- rang_part_shift$clado_part_siz_bef_host
# clado_part_siz_aft_host <- rang_part_shift$clado_part_siz_aft_host
# clado_part_siz_prop_host <- rang_part_shift$clado_part_siz_prop_host
# clado_part_dist_gain_host <- rang_part_shift$clado_part_dist_gain_host
# clado_part_dist_loss_host <- rang_part_shift$clado_part_dist_loss_host
# 
# comb_part_siz_bef_host <- rang_part_shift$comb_part_siz_bef_host
# comb_part_siz_aft_host <- rang_part_shift$comb_part_siz_aft_host
# comb_part_siz_prop_host <- rang_part_shift$comb_part_siz_prop_host
# comb_part_dist_gain_host <- rang_part_shift$comb_part_dist_gain_host
# comb_part_dist_loss_host <- rang_part_shift$comb_part_dist_loss_host
# 
# 
# 
# ana_rang_siz_bef_dep <- rang_part_shift$ana_rang_siz_bef_non
# ana_rang_siz_aft_dep <- rang_part_shift$ana_rang_siz_aft_non
# ana_rang_siz_prop_dep <- rang_part_shift$ana_rang_siz_prop_non
# ana_rang_dist_gain_dep <- rang_part_shift$ana_rang_dist_gain_non
# ana_rang_dist_loss_dep <- rang_part_shift$ana_rang_dist_loss_non
# 
# clado_rang_siz_bef_dep <- rang_part_shift$clado_rang_siz_bef_non
# clado_rang_siz_aft_dep <- rang_part_shift$clado_rang_siz_aft_non
# clado_rang_siz_prop_dep <- rang_part_shift$clado_rang_siz_prop_non
# clado_rang_dist_gain_dep <- rang_part_shift$clado_rang_dist_gain_non
# clado_rang_dist_loss_dep <- rang_part_shift$clado_rang_dist_loss_non
# 
# comb_rang_siz_bef_dep <- rang_part_shift$comb_rang_siz_bef_non
# comb_rang_siz_aft_dep <- rang_part_shift$comb_rang_siz_aft_non
# comb_rang_siz_prop_dep <- rang_part_shift$comb_rang_siz_prop_non
# comb_rang_dist_gain_dep <- rang_part_shift$comb_rang_dist_gain_non
# comb_rang_dist_loss_dep <- rang_part_shift$comb_rang_dist_loss_non
# 
# ana_part_siz_bef_dep <- rang_part_shift$ana_part_siz_bef_non
# ana_part_siz_aft_dep <- rang_part_shift$ana_part_siz_aft_non
# ana_part_siz_prop_dep <- rang_part_shift$ana_part_siz_prop_non
# ana_part_dist_gain_dep <- rang_part_shift$ana_part_dist_gain_non
# ana_part_dist_loss_dep <- rang_part_shift$ana_part_dist_loss_non
# 
# clado_part_siz_bef_dep <- rang_part_shift$clado_part_siz_bef_non
# clado_part_siz_aft_dep <- rang_part_shift$clado_part_siz_aft_non
# clado_part_siz_prop_dep <- rang_part_shift$clado_part_siz_prop_non
# clado_part_dist_gain_dep <- rang_part_shift$clado_part_dist_gain_non
# clado_part_dist_loss_dep <- rang_part_shift$clado_part_dist_loss_non
# 
# comb_part_siz_bef_dep <- rang_part_shift$comb_part_siz_bef_non
# comb_part_siz_aft_dep <- rang_part_shift$comb_part_siz_aft_non
# comb_part_siz_prop_dep <- rang_part_shift$comb_part_siz_prop_non
# comb_part_dist_gain_dep <- rang_part_shift$comb_part_dist_gain_non
# comb_part_dist_loss_dep <- rang_part_shift$comb_part_dist_loss_non


# read in all the calculated network-level variables
# read in quantitative modularity
#modularity_quant_no <- read.csv("Output_no_extinction/mod_quant.csv", header = T)[,2]
#modularity_quant_ex <- read.csv("Output_extinction/mod_quant.csv", header = T)[,2]
#modularity_quant <- c(modularity_quant_no, modularity_quant_ex)

# read in binary modularity
#modularity_bin_no <- read.csv("Output_no_extinction/mod_bin.csv", header = T)[,2]
#modularity_bin_ex <- read.csv("Output_extinction/mod_bin.csv", header = T)[,2]
#modularity_bin <- c(modularity_bin_no, modularity_bin_ex)

# read in quantitative nestedness
#nestedness_quant_no <- read.csv("Output_no_extinction/nest_quant.csv", header = T)[,2]
#nestedness_quant_ex <- read.csv("Output_extinction/nest_quant.csv", header = T)[,2]
#nestedness_quant <- c(nestedness_quant_no, nestedness_quant_ex)

# read in binary nestedness
#nestedness_bin_no <- read.csv("Output_no_extinction/nest_bin.csv", header = T)[,2]
#nestedness_bin_ex <- read.csv("Output_extinction/nest_bin.csv", header = T)[,2]
#nestedness_bin <- c(nestedness_bin_no, nestedness_bin_ex)

# read in interaction turnver data
# int_turn_no <- read.csv("Output_no_extinction/int_turn.csv", header = T)
# int_turn_ex <- read.csv("Output_extinction/int_turn.csv", header = T)
# int_turn <- rbind(int_turn_no, int_turn_ex)
# 
# gain_turn_both <- int_turn$gain_sp_both/int_turn$total_pairwise
# gain_turn_host <- int_turn$gain_sp_host/int_turn$total_pairwise
# gain_turn_dep <- int_turn$gain_sp_non/int_turn$total_pairwise
# 
# gain_rew <- int_turn$gain_rew/int_turn$total_pairwise
# 
# gain_total <- (gain_turn_both + gain_turn_host + gain_turn_dep + gain_rew) * int_turn$total_pairwise
# 
# loss_turn_both <- int_turn$loss_ex_both/int_turn$total_pairwise
# loss_turn_host <- int_turn$loss_ex_host/int_turn$total_pairwise
# loss_turn_dep <- int_turn$loss_ex_non/int_turn$total_pairwise
# 
# loss_rew <- int_turn$loss_rew/int_turn$total_pairwise
# 
# loss_total <- loss_turn_both + loss_turn_host + loss_turn_dep + loss_rew * int_turn$total_pairwise

# read in clade-level variables
# read in partnership conservatism
#part_cons_no <- read.csv("Output_no_extinction/part_cons.csv", header = T)
#part_cons_ex <- read.csv("Output_extinction/part_cons.csv", header = T)
#part_cons <- rbind(part_cons_no, part_cons_ex)

#part_cons_host_w <- part_cons$host_part_cons_w
#part_cons_dep_w <- part_cons$non_part_cons_w

#part_cons_host_u <- part_cons$host_part_cons_u
#part_cons_dep_u <- part_cons$non_part_cons_u

# read in trait conservatism
#trait_cons_no <- read.csv("Output_no_extinction/trait_cons.csv", header = T)
#trait_cons_ex <- read.csv("Output_extinction/trait_cons.csv", header = T)
#trait_cons <- rbind(trait_cons_no, trait_cons_ex)

#K_host <- trait_cons$host_K
#K_dep <- trait_cons$non_K

#lambda_host <- trait_cons$host_lambda
#lambda_dep <- trait_cons$non_lambda

# read in specificity data
# specificity_no <- read.csv("Output_no_extinction/specificity.csv", header = T)
# specificity_ex <- read.csv("Output_extinction/specificity.csv", header = T)
# specificity <- rbind(specificity_no, specificity_ex)
# 
# bas_spec_host <- specificity$bas_spec_host
# bas_spec_dep <- specificity$bas_spec_non
# 
# struct_spec_host <- specificity$struct_spec_host
# struct_spec_dep <- specificity$struct_spec_non
# 
# phylo_spec_host <- specificity$phylo_spec_host
# phylo_spec_dep <- specificity$phylo_spec_non
# 
# func_spec_host <- specificity$func_spec_host
# func_spec_dep <- specificity$func_spec_non

# read in species turnover and diversity
div_turn_no <- read.csv("Output_no_extinction/div_turn.csv", header = T)
div_turn_ex <- read.csv("Output_extinction/div_turn.csv", header = T)
div_turn <- rbind(div_turn_no, div_turn_ex)

# turn_host <- (div_turn$sp_host + div_turn$ex_host) / div_turn$div_host
# turn_dep <- (div_turn$sp_non + div_turn$ex_non) / div_turn$div_non
# turn_host <- div_turn$sp_host + div_turn$ex_host
# turn_dep <- div_turn$sp_non + div_turn$ex_non

div_host <- div_turn$div_host
div_dep <- div_turn$div_non

# read in mean phylogenetic distance among host lineages
#phylo_dist_host_no <- read.csv("Output_no_extinction/phylo_dist_host.csv", header = T)
#phylo_dist_host_ex <- read.csv("Output_extinction/phylo_dist_host.csv", header = T)
#phylo_dist_host <- rbind(phylo_dist_host_no, phylo_dist_host_ex)$pd_host

# store all data in a data frame
dat_raw <- data.frame(
  
  nam,
  
  # Colonization on new geographic sites
  1 - prop_col_host,
  1 - prop_col_dep,
  
  # age
  #log2(age_host),
  #log2(age_dep),
  
  # trait variation
  #trait_var_host,
  #trait_var_dep,
  
  # Range size fluctuation
  #rang_siz_mean_host , # rang_siz$rang_siz_host
  #rang_siz_mean_dep , # rang_siz$rang_siz_non
  
  #rang_siz_sum_host , # rang_siz$rang_siz_host_sum
  #rang_siz_sum_dep , # rang_siz$rang_siz_non_sum
  
  #comb_rang_dist_gain_host , # rang_part_shift$comb_rang_dist_gain_host
  #comb_rang_dist_loss_host , # rang_part_shift$comb_rang_dist_loss_host
  
  #comb_rang_dist_gain_dep , # rang_part_shift$comb_rang_dist_gain_dep
  #comb_rang_dist_loss_dep , # rang_part_shift$comb_rang_dist_loss_dep
  
  # Demographic stochasticity
  #dem_stoch_mean_host , # dem_stoch$dem_stoch_host
  #dem_stoch_mean_dep , # dem_stoch$dem_stoch_non
  
  # Hybridization
  as.numeric(hybr_lik_mean_host) , # hybr_lik$hybr_lik_host
  as.numeric(hybr_lik_mean_dep) , # hybr_lik$hybr_lik_non
  
  # trait temporal variation
  as.numeric(delta_host),
  as.numeric(delta_dep),
  
  # Speciation and extinction
  #turn_host,
  #turn_dep,
  div_host,
  div_dep#,
  
  # Partnership dynamics
  #comb_part_siz_aft_host , # rang_part_shift$comb_part_siz_aft_host
  #comb_part_siz_aft_dep , # rang_part_shift$comb_part_siz_aft_dep
  
  #comb_part_dist_gain_host , # rang_part_shift$comb_part_dist_gain_host
  #comb_part_dist_gain_dep #, #, rang_part_shift$comb_part_dist_gain_dep
  
  # phylogenetic distance among hosts
  #phylo_dist_host
  
)

# figure out which variables contain NAs
colSums(is.na(dat_raw))

# remove replicates with NAs
dat_na_rm <- dat_raw[!is.na(rowSums(dat_raw[,-1])),]
sum(!is.na(rowSums(dat_raw[,-1]))) / 4800

# add independent variables to the data frame
dat_na_rm$host_spec <- rep(NA, length(dat_na_rm$nam))
dat_na_rm$host_spec[substr(dat_na_rm$nam, 1, 1) == 'a' & substr(dat_na_rm$nam, 3, 3) == '1'] <- "neutral+"
dat_na_rm$host_spec[substr(dat_na_rm$nam, 1, 1) == 'a' & substr(dat_na_rm$nam, 3, 3) == '0'] <- "neutral-"
dat_na_rm$host_spec[substr(dat_na_rm$nam, 1, 1) == 's' & substr(dat_na_rm$nam, 3, 3) == '1'] <- "divergent+"
dat_na_rm$host_spec[substr(dat_na_rm$nam, 1, 1) == 's' & substr(dat_na_rm$nam, 3, 3) == '0'] <- "divergent-"

dat_na_rm$dep_spec <- rep(NA, length(dat_na_rm$nam))
dat_na_rm$dep_spec[substr(dat_na_rm$nam, 2, 2) == 'a' & substr(dat_na_rm$nam, 6, 6) == '1'] <- "neutral+"
dat_na_rm$dep_spec[substr(dat_na_rm$nam, 2, 2) == 'a' & substr(dat_na_rm$nam, 6, 6) == '0'] <- "neutral-"
dat_na_rm$dep_spec[substr(dat_na_rm$nam, 2, 2) == 's' & substr(dat_na_rm$nam, 6, 6) == '1'] <- "divergent+"
dat_na_rm$dep_spec[substr(dat_na_rm$nam, 2, 2) == 's' & substr(dat_na_rm$nam, 6, 6) == '0'] <- "divergent-"

dat_na_rm$host_fit_func <- rep(NA, length(dat_na_rm$nam))
dat_na_rm$host_fit_func[substr(dat_na_rm$nam, 5, 5) == 0] <- "Unaffected (0)"
dat_na_rm$host_fit_func[substr(dat_na_rm$nam, 5, 5) == '1' & substr(dat_na_rm$nam, 7, 7) == 'M' & substr(dat_na_rm$nam, 8, 8) == 'M'] <- "Trait matching (+)"
dat_na_rm$host_fit_func[substr(dat_na_rm$nam, 5, 5) == '1' & substr(dat_na_rm$nam, 7, 7) == 'M' & substr(dat_na_rm$nam, 8, 8) == 'D'] <- "Trait difference (+)"
dat_na_rm$host_fit_func[substr(dat_na_rm$nam, 5, 5) == '1' & substr(dat_na_rm$nam, 7, 7) == 'A' & substr(dat_na_rm$nam, 8, 8) == 'M'] <- "Trait matching (-)"
dat_na_rm$host_fit_func[substr(dat_na_rm$nam, 5, 5) == '1' & substr(dat_na_rm$nam, 7, 7) == 'A' & substr(dat_na_rm$nam, 8, 8) == 'D'] <- "Trait difference (-)"

dat_na_rm$dep_fit_func <- rep(NA, length(dat_na_rm$nam))
dat_na_rm$dep_fit_func[substr(dat_na_rm$nam, 4, 4) == 0] <- "Unaffected (0)"
dat_na_rm$dep_fit_func[substr(dat_na_rm$nam, 4, 4) == '1' & substr(dat_na_rm$nam, 9, 9) == 'M'] <- "Trait matching (+)"
dat_na_rm$dep_fit_func[substr(dat_na_rm$nam, 4, 4) == '1' & substr(dat_na_rm$nam, 9, 9) == 'D'] <- "Trait difference (+)"

dat_na_rm$catastrophe <- rep(NA, length(dat_na_rm$nam))
dat_na_rm$catastrophe[substr(dat_na_rm$nam, 11, 12) == "no"] <- "no"
dat_na_rm$catastrophe[substr(dat_na_rm$nam, 11, 12) == "ex"] <- "yes"


# Add independent variables based on scenario and reciprocity
dat_na_rm$scenario <- rep(NA, length(dat_na_rm$nam))

# 
dat_na_rm$scenario[dat_na_rm$host_fit_func == "Trait difference (+)" & dat_na_rm$dep_fit_func == "Trait difference (+)"] <- "a"

dat_na_rm$scenario[dat_na_rm$host_fit_func == "Trait difference (+)" & dat_na_rm$dep_fit_func == "Unaffected (0)"] <- "b"

dat_na_rm$scenario[dat_na_rm$host_fit_func == "Unaffected (0)" & dat_na_rm$dep_fit_func == "Trait difference (+)"] <- "c"

dat_na_rm$scenario[dat_na_rm$host_fit_func == "Trait matching (+)" & dat_na_rm$dep_fit_func == "Trait matching (+)"] <- "d"

dat_na_rm$scenario[dat_na_rm$host_fit_func == "Trait matching (+)" & dat_na_rm$dep_fit_func == "Unaffected (0)"] <- "e"

dat_na_rm$scenario[dat_na_rm$host_fit_func == "Unaffected (0)" & dat_na_rm$dep_fit_func == "Trait matching (+)"] <- "f"

dat_na_rm$scenario[dat_na_rm$host_fit_func == "Trait matching (-)" & dat_na_rm$dep_fit_func == "Trait matching (+)"] <- "g"

dat_na_rm$scenario[dat_na_rm$host_fit_func == "Trait matching (-)" & dat_na_rm$dep_fit_func == "Unaffected (0)"] <- "h"

dat_na_rm$scenario[dat_na_rm$host_fit_func == "Unaffected (0)" & dat_na_rm$dep_fit_func == "Unaffected (0)"] <- "i"

# add clade specific scenario order
dat_na_rm$scenario_host <- factor(dat_na_rm$scenario, levels=c('a','g','d','e','b','c','f','h','i'))
dat_na_rm$scenario_dep <- factor(dat_na_rm$scenario, levels=c('a','g','d','e','f','b','c','h','i'))

# Add highlight colors
dat_na_rm$mode_host <- rep('No change predicted', length(dat_na_rm$nam))

dat_na_rm$mode_host[dat_na_rm$scenario == "a" | dat_na_rm$scenario == "g"] <- "Destabilizing selection hypothesis\npredicts a decrease"
dat_na_rm$mode_host[dat_na_rm$scenario == "d" | dat_na_rm$scenario == "e"] <- "Stabilizing selection hypothesis\npredicts an increase"

dat_na_rm$mode_host <- factor(dat_na_rm$mode_host,levels=c("Destabilizing selection hypothesis\npredicts a decrease",  "Stabilizing selection hypothesis\npredicts an increase", "No change predicted"))

dat_na_rm$mode_dep <- rep('No change predicted', length(dat_na_rm$nam))

dat_na_rm$mode_dep[dat_na_rm$scenario == "a" | dat_na_rm$scenario == "g"] <- "Destabilizing selection hypothesis\npredicts a decrease"
dat_na_rm$mode_dep[dat_na_rm$scenario == "d" | dat_na_rm$scenario == "e" | dat_na_rm$scenario == "f"] <- "Stabilizing selection hypothesis\npredicts an increase"

dat_na_rm$mode_dep <- factor(dat_na_rm$mode_dep,levels=c("Destabilizing selection hypothesis\npredicts a decrease",  "Stabilizing selection hypothesis\npredicts an increase", "No change predicted"))

# subset the original dataset to neutral speciation entries only
#dat_na_rm <- dat_na_rm[dat_na_rm$host_spec=='neutral-' | dat_na_rm$host_spec=='neutral+',]
#dat_na_rm <- dat_na_rm[dat_na_rm$dep_spec=='neutral-' | dat_na_rm$dep_spec=='neutral+',]

# rename columns
colnames(dat_na_rm) <- c(
  'nam',
  'Proportion_of_locals_among_host_individuals',
  'Proportion_of_locals_among_dependent_individuals',
  #'Age_of_host_individuals',
  #'Age_of_dependent_individuals',
  #'Trait_variation_between_host_individuals_in_adjacent_sites',
  #'Trait_variation_between_dependent_individuals_in_adjacent_sites',
  #'Range_size_H',
  #'Range_size_D',
  #'Distance_of_geographic_range_change_H',
  #'Distance_of_geographic_range_change_D',
  #'Demographic_stochasticity_H',
  #'Demographic_stochasticity_D',
  'Difficulty_of_hybridization_H',
  'Difficulty_of_hybridization_D',
  'Delta_H',
  'Delta_D',
  #'Clade_turnover_H',
  #'Clade_turnover_D',
  'Clade_diversity_H',
  'Clade_diversity_D',
  #'Specificity_H',
  #'Specificity_D',
  #'Distance_of_partner_shifts_H',
  #'Distance_of_partner_shifts_D',
  #'Phylogenetic_distance_among_host_lineages',
  'Speciation_condition_H',
  'Speciation_condition_D',
  'Fitness outcome for the host',
  'Fitness outcome for the dependent',
  'Local_extinction_events',
  'Scenario',
  'Scenario_host',
  'Scenario_dep',
  'Mode of evolution for hosts',
  'Mode of evolution for dependents'
)

#(optional depending on what is needed)
# logit transfrom the proportional variables
# dat_na_rm$Proportion_of_colonists_H <- log(dat_na_rm$Proportion_of_colonists_H / (1 - dat_na_rm$Proportion_of_colonists_H))
# dat_na_rm$Proportion_of_colonists_D <- log(dat_na_rm$Proportion_of_colonists_D / (1 - dat_na_rm$Proportion_of_colonists_D))

# save the original dataset
dat_na_rm_original <- dat_na_rm
```
