## Supplementary material for "Coevolution-induced stabilizing and destabilizing selection shapes species richness in clade co-diversification": File S1: README.md

# Coevolution-induced stabilizing and destabilizing selection shapes species richness in clade codiversification

## Description
This project is for simulating eco-evolutionary dynamics described in "Coevolution-induced stabilizing and destabilizing selection shapes species richness in clade codiversification" authored by Yichao Zeng, David H. Hembry, and John J. Wiens. It is written in the R language (4.0.0) and to be run on computing clusters running on Slurm Workload Manager.

## Table of Contents
- [Setup](#setup)
- [Simulation](#simulation)
- [Analyses](#analyses)

## Setup
1. Copy all files to your working directory on the computing cluster.
2. Unzip 'slurm.zip' and 'control.zip'.
3. Create two folders under the working directory, one named "Output_extinction" and the other named "Output_no_extinction". Extinction vs. no extinction are obsolete features of the model that have been nullified, but the folder names need to be keep intact to generate the desired number of replicates.
4. Replace the default working directory "Desktop/model_rcpp_hpc_OneDrive" with your own working directory in the code.
5. Depending on the user's R library, packages will need to be installed before they can be used.

## Simulation
To simulate the described eco-evolutionary dynamics:

1. Run the code in batch_script/simulation.text by copying it to your command line, and the simulation output will be stored in the two folders that you created in Step 2 in Setup.
2. (Optional) Run the code in batch_script/cancel_all_jobs.text by copying it to your command line if you need to cancel all jobs.

## Analyses
1. To quantify trait temporal variation as mean absolute step difference (the mean absolute value of delta z): Run 'sbatch Desktop/model_rcpp_hpc_OneDrive/slurm/delta_ex.slurm' and
'sbatch Desktop/model_rcpp_hpc_OneDrive/slurm/delta_noex.slurm'. The output will be saved to the two folders that you created Step 2 in Setup.

2. To quantify the degree of dispersal limitation as the proportion of native individuals among all individuals: Run 'sbatch Desktop/model_rcpp_hpc_OneDrive/slurm/prop_col_ex.slurm' and
'sbatch Desktop/model_rcpp_hpc_OneDrive/slurm/prop_col_noex.slurm'. The output will be saved to the two folders that you created Step 2 in Setup.

3. To quantify genetic distance between geographic sites: Run 'sbatch Desktop/model_rcpp_hpc_OneDrive/slurm/hybr_lik_ex.slurm' and
'sbatch Desktop/model_rcpp_hpc_OneDrive/slurm/hybr_lik_noex.slurm'. The output will be saved to the two folders that you created Step 2 in Setup.

4. To quantify species richness accomulated: Run 'calc_div_turn.R' twice, first time with 'output_dir <- "Output_extinction"' and second time with 'output_dir <- "Output_no_extinction"'. This step does not necessarily need to be run on a computing cluster. 

5. To read in the quantified variables: Run 'read_in_data.R'.

6. To visualize the relationshipts between selective regime, degree of dispersal limitation, genetic differentiation, and species richness accumulated: Run 'figure_continuous.R'. The plot will be saved as a PDF to your work directory.

7. To visualize, for each simulated scenario, a typical time series where trait value changes over time: Run 'trait_single_cell_zoom_in_out.R'. The plot will be saved as a PDF to your work directory.

## Contact information
- Any questions or comments are welcome and should be sent to Yichao Zeng (yichaozeng.weebly.com).
