## Supplementary material for "Coevolution-induced stabilizing and destabilizing selection shapes species richness in clade co-diversification": File S1: reproduction_combined_smart_det_repr.R

```
# reproduction function that updates the host list
host_repr <- function(grid_host, host_list, non_list, repro.site.id, w_0, c_mat_host, no_offspring_max, choos_thres, comp_thres, host_no_comp_loci, parent_origin, host_mu, grid_bar, n_dimension, sigma_eco, sigma_pre){
  
  #reproduction cannot occur if there is only one individual
  if(grid_host[repro.site.id[1], repro.site.id[2]] < 2){
    return(list(host_list, host_no_comp_loci, parent_origin))
  }
  
  else{
    
    #no_mating_max <- c_mat_host * (grid_host[repro.site.id[1], repro.site.id[2]])^2
    #no_mating_max <- floor(no_mating_max) + 1
    #no_mating_max <- 1
    no_mating_max <- c_mat_host
    
    #find all host IDs in the reproduction site
    host.on.site <- NULL
    for (i in 1:length(host_list)){
      if(length(host_list[[i]]) == 8){ #ensure alive
        if(identical(as.numeric(host_list[[i]]$site.id), as.numeric(repro.site.id)) == T){
          host.on.site <- c(host.on.site, i)
        }
      }
      
    }
    
    #if(length(host.on.site)<2){ #reproduction cannot occur if fewer than 2 individuals
    #  return(host_list)
    #}
    
    #else{
      
      #create a vector containing fitness for every host individual in the host list
      host.fit <- rep(w_0, length(host_list))

      # calculate fitness for each host in this site molded by coevolution
      for (i in 1:length(non_list)) {
        if(length(non_list[[i]]) == 8 && identical(as.numeric(non_list[[i]]$site.id), as.numeric(repro.site.id))){
          host.fit[non_list[[i]]$host.id] <- host.fit[non_list[[i]]$host.id] + fit_func_host(zi = get_eco_trait_val(host_list[[non_list[[i]]$host.id]]), zj = get_eco_trait_val(non_list[[i]]), xi = xi_host, alpha = alpha_host)            
        }
      }
      
      host.fit.check <- host.fit[host.on.site]

      #if(length(host.fit.check) < 2){ # reproduction cannot happen if there are fewer than 2 individuals
      #  return(list(host_list = host_list, host_no_comp_loci = host_no_comp_loci))
      #} 
      #else{
        
        # each host's fitness is further molded by selection by the environment. Note: ladder is a parameter to help circumvent R's lower limit for numeric type
        #ladder <- 0
        #host.fit.temp <- host.fit
        #while(max(host.fit.temp) < Inf){ # until one of the values is Inf
        #  host.fit.temp <- host.fit
        #  for (host_env in host.on.site) {
        #    z_star <- grid.opt.host[host_list[[host_env]]$site.id[1], host_list[[host_env]]$site.id[2]]
        #    host.fit.temp[host_env] <- host.fit.temp[host_env] * exp(-gamma_host * (get_eco_trait_val(host_list[[host_env]]) - z_star)^2 + ladder)
        #  }
        #  host.fit.temp <- host.fit.temp[host.on.site]
        #  
        #  if(length(host.fit.temp[host.fit.temp>0]) >= 2){
        #    break
        #  }
        #  ladder <- ladder + 100
        #}
        
        host.fit <- host.fit.check
		
		# probability of each host individual being drawn as a parent (Note that if any individual has a fitness of Inf, there must not be enough individuals of positive mating probability)
		#host.par.prob <- host.fit/sum(host.fit)
		host.par.prob <- host.fit
		
		#if(length(host.par.prob[host.par.prob > 0]) >= 2){
  		    # repeat until a fixed number of mating attempts have been made
          no.mat.attempt <- 0
          while(no.mat.attempt < no_mating_max){
            # choose 1st and 2nd parents
            # choose 1st and 2nd parents
            ids <- c(1:length(host.par.prob))
            id.1 <- ids
            if(length(id.1) > 1){
              id.1 <- sample(x = ids[host.par.prob == max(host.par.prob)], size = 1, replace = F)                
            }
            ids.temp <- ids[ids != id.1]
            id.2 <- ids.temp
            if(length(id.2) > 1){
              id.2 <- sample(x = ids.temp[host.par.prob[ids.temp] == max(host.par.prob[ids.temp])], size = 1, replace = F)                
            }
            
            #ids <- sample(x = 1:length(host.par.prob), size = 2, prob = host.par.prob)
            #id.1 <- ids[1]
            #id.2 <- ids[2]
            
            par.id.1 <- host.on.site[id.1]
            par.id.2 <- host.on.site[id.2]
            
            par_1 <- host_list[[par.id.1]]
            par_2 <- host_list[[par.id.2]]
            
            #determine whether the mating can result in successful reproduction
            ## prezygotic isolation  
            choos.prob <- mat_prob(par_1, par_2, c_am)
            pre.iso <- (choos.prob < choos_thres)
            
            ## postzygotic isolation
            post.iso <- F#(incomp_count(par_1, par_2) > comp_thres)
            
            #one offspring is produced and added to host list
            if(pre.iso==F && post.iso==F){
              # increase n_repr for both parents
              #host_list[[par.id.1]]$n_repr <- host_list[[par.id.1]]$n_repr + 1
              #host_list[[par.id.2]]$n_repr <- host_list[[par.id.2]]$n_repr + 1
              # initialize the number of offspring already produced
              no.offspring <- 0
              while (no.offspring < no_offspring_max) {
    
    #############################################################################################
                  
                  # inherit host.id (for non-hosts only) & site.id
                  offspring <- par_1
                  
                  # erase the number of successful reproductions
                  #offspring$n_repr <- 0
                  
                  # erase age
                  offspring$n_life <- 0
                  
                  # erase the number of non-hosts (for hosts only)
                  offspring$n_non <- 0
                  
                  # initialize origin
                  offspring$origin <- "local"
                  
                  # gamete from 1st parent
                  gam1 <- par_1
                  for(i in 1:L_z){
                    gam1$eco[[i]] <- rnorm(n=1, mean = sample(par_1$eco[[i]], size=1), sd = sigma_eco)
                  }
                  for(i in 1:L_a){
                    gam1$pre[[i]] <- rnorm(n=1, mean = sample(par_1$pre[[i]], size=1), sd = sigma_pre)
                  }
                  for(i in 1:L_b){
                    
                    allele_val <- sample(par_1$post[[i]], 1)
                    if(rbern(1, host_mu)==1){
                      allele_val <- host_no_comp_loci[i] + 1 # because the mutations must be a new one, following the infinite sites model
                      host_no_comp_loci[i] <- allele_val
                    }
                    
                    gam1$post[[i]] <- allele_val
                  }
                  
                  # gamete from 2nd parent
                  gam2 <- par_2
                  for(i in 1:L_z){
                    gam2$eco[[i]] <- rnorm(n=1, mean = sample(par_2$eco[[i]], size=1), sd = sigma_eco)
                  }
                  for(i in 1:L_a){
                    gam2$pre[[i]] <- rnorm(n=1, mean = sample(par_2$pre[[i]], size=1), sd = sigma_pre)
                  }
                  for(i in 1:L_b){
                    
                    allele_val <- sample(par_2$post[[i]], 1)
                    if(rbern(1, host_mu)==1){
                      allele_val <- host_no_comp_loci[i] + 1 # because the mutations must be a new one, following the infinite sites model
                      host_no_comp_loci[i] <- allele_val
                    }
                    
                    gam2$post[[i]] <- allele_val
                  }
                  
                  # combine into a list
                  gams <- list(gam1, gam2)
                  
                  #generate genotype
                  for (j in 1:2) {
                    
                    for(i in 1:L_z){
                      offspring$eco[[i]][j] <- gams[[j]]$eco[[i]]
                    }
                    for(i in 1:L_a){
                      offspring$pre[[i]][j] <- gams[[j]]$pre[[i]]
                    }
                    for(i in 1:L_b){
                      offspring$post[[i]][j] <- gams[[j]]$post[[i]]
                    }
                    
                  }
                  
    
                  
                
    ###########################################################################            
                
                host_list[[length(host_list)+1]] <- host_ind_mov(offspring, grid_bar, n_dimension)
                
                no.offspring <- no.offspring + 1
                
                # update the parent origin data frame
                count_col <- (par_1$origin == "colonist") + (par_2$origin == "colonist")
                if(count_col==0)
                  parent_origin$host_loc_loc[gen_id] <- parent_origin$host_loc_loc[gen_id] + 1
                else if(count_col==1)
                  parent_origin$host_loc_col[gen_id] <- parent_origin$host_loc_col[gen_id] + 1
                else if(count_col==2)
                  parent_origin$host_col_col[gen_id] <- parent_origin$host_col_col[gen_id] + 1
                  
                
              }
              
            }
            
            no.mat.attempt <- no.mat.attempt + 1
            
          }
          
          #if(min(host.fit) <= 0){
          #  return(list(host_list = NULL, host_no_comp_loci = NULL, parent_origin = NULL))
          #}else{
            return(list(host_list = host_list, host_no_comp_loci = host_no_comp_loci, parent_origin = parent_origin))
          
          #}

        
        #}        		      
		    #}
		    #else{
		    #  return(list(host_list = host_list, host_no_comp_loci = host_no_comp_loci))
		    #}

      #}

    
    
  }
  
  
}


# reproduction function that updates the non-host list
non_repr <- function(grid_non, non_list, grid_host, host_list, repro.site.id, w_0, c_mat_non, no_offspring_max, choos_thres, comp_thres, non_no_comp_loci, parent_origin, non_mu, grid_bar, n_dimension, sigma_eco, sigma_pre){
  
  #reproduction cannot occur if there is only one individual
  if(grid_non[repro.site.id[1], repro.site.id[2]] < 2){
    return(list(non_list, non_no_comp_loci, parent_origin))
  }
  
  else{
    
    #no_mating_max <- c_mat_non * (grid_non[repro.site.id[1], repro.site.id[2]])^2
    #no_mating_max <- floor(no_mating_max) + 1
    #no_mating_max <- 1
    no_mating_max <- c_mat_non
    
    #find all non-host IDs in the reproduction site
    non.on.site <- NULL
    for (i in 1:length(non_list)){
      if(length(non_list[[i]]) == 8){ #ensure alive
        if(identical(as.numeric(non_list[[i]]$site.id), as.numeric(repro.site.id)) == T){
          non.on.site <- c(non.on.site, i)
        }
      }
      
    }
    
    #if(length(host.on.site)<2){ #reproduction cannot occur if fewer than 2 individuals
    #  return(host_list)
    #}
    
    #else{
      
      #create a vector containing fitness for every non-host individual in the non-host list
      non.fit <- rep(w_0, length(non_list))
      
      # calculate fitness for each non-host in this site molded by coevolution
      for (i in 1:length(non_list)) {
        if(length(non_list[[i]]) == 8 && identical(as.numeric(non_list[[i]]$site.id), as.numeric(repro.site.id))){
          non.fit[i] <- non.fit[i] + fit_func_non(zi = get_eco_trait_val(non_list[[i]]), zj = get_eco_trait_val(host_list[[non_list[[i]]$host.id]]), xi = xi_non, alpha = alpha_non)
        }
      }
      
      non.fit.check <- non.fit[non.on.site]
      
      #if(length(non.fit.check[non.fit.check!=0]) < 2){ # reproduction cannot happen if all individuals have zero fitness (including low fitness rounded by R to 0) or fewer than two individuals have non-zerofitness
      #  return(list(non_list = non_list, non_no_comp_loci = non_no_comp_loci))
      #} 
	  #  else{
        
		# each non-host's fitness is further molded by selection by the environment. Note: ladder is a parameter to help circumvent R's lower limit for numeric type
        #ladder <- 0
        #non.fit.temp <- non.fit
        #while(max(non.fit.temp) < Inf){ # until one of the values is Inf
        #  non.fit.temp <- non.fit
        #  for (non_env in non.on.site) {
        #    z_star <- grid.opt.non[non_list[[non_env]]$site.id[1], non_list[[non_env]]$site.id[2]]
        #    non.fit.temp[non_env] <- non.fit.temp[non_env] * exp(-gamma_non * (get_eco_trait_val(non_list[[non_env]]) - z_star)^2 + ladder)
        #  }
        #  non.fit.temp <- non.fit.temp[non.on.site]
        #  
        #  if(length(non.fit.temp[non.fit.temp>0]) >= 2){
        #    break
        #  }
        #  ladder <- ladder + 100
        #}
		
		non.fit <- non.fit.check
		
		# probability of each _non-host individual being drawn as a parent
        #non.par.prob <- non.fit/sum(non.fit)
        non.par.prob <- non.fit
        
        if(min(non.par.prob) <= 0){
          break
        }
        
        #if(length(non.par.prob[non.par.prob > 0]) >= 2){
            # repeat until a fixed number of mating attempts have been made
            no.mat.attempt <- 0
            while(no.mat.attempt < no_mating_max){
              # choose 1st and 2nd parents
              ids <- c(1:length(non.par.prob))
              id.1 <- ids
              if(length(id.1) > 1){
                id.1 <- sample(x = ids[non.par.prob == max(non.par.prob)], size = 1, replace = F)                
              }
              ids.temp <- ids[ids != id.1]
              id.2 <- ids.temp
              if(length(id.2) > 1){
                id.2 <- sample(x = ids.temp[non.par.prob[ids.temp] == max(non.par.prob[ids.temp])], size = 1, replace = F)                
              }
              
              #ids <- sample(x = 1:length(non.par.prob), size = 2, prob = non.par.prob)
              #id.1 <- ids[1]
              #id.2 <- ids[2]
              
              par.id.1 <- non.on.site[id.1]
              par.id.2 <- non.on.site[id.2]
              
              par_1 <- non_list[[par.id.1]]
              par_2 <- non_list[[par.id.2]]
              
              #determine whether the mating can result in successful reproduction
              ## prezygotic isolation  
              choos.prob <- mat_prob(par_1, par_2, c_am)
              pre.iso <- (choos.prob < choos_thres)
              
              ## postzygotic isolation
              post.iso <- F#(incomp_count(par_1, par_2) > comp_thres)
              
              #one offspring is produced and added to non-host list
              if(pre.iso==F && post.iso==F){
                # increase n_repr for both parents
                #non_list[[par.id.1]]$n_repr <- non_list[[par.id.1]]$n_repr + 1
                #non_list[[par.id.2]]$n_repr <- non_list[[par.id.2]]$n_repr + 1
                
                # initialize the number of offspring already produced
                no.offspring <- 0
                while (no.offspring < no_offspring_max) {
                  
                  #############################################################################################
                  
                  # inherit host.id (for non-hosts only) & site.id
                  offspring <- par_1
                  
                  # erase the number of successful reproductions
                  #offspring$n_repr <- 0
                  
                  # erase age
                  offspring$n_life <- 0
                  
                  # initialize origin
                  offspring$origin <- "local"
                  
                  # gamete from 1st parent
                  gam1 <- par_1
                  for(i in 1:L_z){
                    gam1$eco[[i]] <- rnorm(n=1, mean = sample(par_1$eco[[i]], size=1), sd = sigma_eco)
                  }
                  for(i in 1:L_a){
                    gam1$pre[[i]] <- rnorm(n=1, mean = sample(par_1$pre[[i]], size=1), sd = sigma_pre)
                  }
                  for(i in 1:L_b){
                    
                    allele_val <- sample(par_1$post[[i]], 1)
                    if(rbern(1, non_mu)==1){
                      allele_val <- non_no_comp_loci[i] + 1 # because the mutations must be a new one, following the infinite sites model
                      non_no_comp_loci[i] <- allele_val
                    }
                    
                    gam1$post[[i]] <- allele_val
                  }
                  
                  # gamete from 2nd parent
                  gam2 <- par_2
                  for(i in 1:L_z){
                    gam2$eco[[i]] <- rnorm(n=1, mean = sample(par_2$eco[[i]], size=1), sd = sigma_eco)
                  }
                  for(i in 1:L_a){
                    gam2$pre[[i]] <- rnorm(n=1, mean = sample(par_2$pre[[i]], size=1), sd = sigma_pre)
                  }
                  for(i in 1:L_b){
                    
                    allele_val <- sample(par_2$post[[i]], 1)
                    if(rbern(1, non_mu)==1){
                      allele_val <- non_no_comp_loci[i] + 1 # because the mutations must be a new one, following the infinite sites model
                      non_no_comp_loci[i] <- allele_val
                    }
                    
                    gam2$post[[i]] <- allele_val
                  }
                  
                  # combine into a list
                  gams <- list(gam1, gam2)
                  
                  #generate genotype
                  for (j in 1:2) {
                    
                    for(i in 1:L_z){
                      offspring$eco[[i]][j] <- gams[[j]]$eco[[i]]
                    }
                    for(i in 1:L_a){
                      offspring$pre[[i]][j] <- gams[[j]]$pre[[i]]
                    }
                    for(i in 1:L_b){
                      offspring$post[[i]][j] <- gams[[j]]$post[[i]]
                    }
                    
                  }
                  
                  
                  
                  
                  ###########################################################################            
                  
                  non_list[[length(non_list)+1]] <- non_ind_mov(offspring, grid_bar, n_dimension, grid_host, host_list)
                  
                  no.offspring <- no.offspring + 1
                  
                  # update the parent origin data frame
                  count_col <- (par_1$origin == "colonist") + (par_2$origin == "colonist")
                  if(count_col==0)
                    parent_origin$non_loc_loc[gen_id] <- parent_origin$non_loc_loc[gen_id] + 1
                  else if(count_col==1)
                    parent_origin$non_loc_col[gen_id] <- parent_origin$non_loc_col[gen_id] + 1
                  else if(count_col==2)
                    parent_origin$non_col_col[gen_id] <- parent_origin$non_col_col[gen_id] + 1
                  
                }
                
              }
              
              no.mat.attempt <- no.mat.attempt + 1
              
            }
            
            #if(min(non.fit) <= 0){
            #  return(list(non_list = NULL, non_no_comp_loci = NULL, parent_origin = NULL))
            #}else{
              return(list(non_list = non_list, non_no_comp_loci = non_no_comp_loci, parent_origin = parent_origin))
            #}
            
        #}
        #else{
        #    return(list(non_list = non_list, non_no_comp_loci = non_no_comp_loci))
        #}
      
      #}
      
      	      
	    #}

  }
  
  
}
```
