## Supplementary material for "Coevolution-induced stabilizing and destabilizing selection shapes species richness in clade co-diversification": File S1: reproduction_pop.R

```
##########################################################################
# for species delineation

# a function that calculates the mating probability between two populations
mat_prob_pop <- function(pop1, pop2, ind_list1, ind_list2, c_am){
  
  #get ecological trait value for both parents
  z1 <- mean(unlist(lapply(ind_list1[pop1], get_eco_trait_val)))
  z2 <- mean(unlist(lapply(ind_list2[pop2], get_eco_trait_val)))
  
  #get choosiness trait value for both parents
  a1 <- mean(unlist(lapply(ind_list1[pop1], get_choos_trait_val)))
  a2 <- mean(unlist(lapply(ind_list2[pop2], get_choos_trait_val)))
  
  #use the choosier parent for probability calculation
  a <- a1
  if(abs(a1 < abs(a2))){
    a <- a2
  }
  
  temp <- ( 1-0.5*exp(-a^2) ) * exp( - (z1 -z2)^2 / (2/(a^2 * c_am))^2 )
  if(a < 0){
    mat_prob <- 1 - temp
  }
  else if(a ==0){
    mat_prob <- 0.5
  }
  else if(a > 0){
    mat_prob <- temp
  }
  
  return(mat_prob)
  
}


# a function that calculates the mean genetic distance between two populations
#sourceCpp("Rcpp/incomp_count_pop.cpp")

incomp_count_pop <- function(pop1, pop2, ind_list1, ind_list2){
    
# create a genetic distance matrix
gen.dist <- matrix(NA, nrow = length(pop1), ncol = length(pop2))

for(i in 1:length(pop1)){
  for(j in 1:length(pop2)){
    
  gen.dist[i,j] <- incomp_count(ind_list1[[pop1[i]]], ind_list2[[pop2[j]]])
    
  }
}

return(mean(gen.dist))
 
}
```
