## Supplementary material for "Coevolution-induced stabilizing and destabilizing selection shapes species richness in clade co-diversification": File S1: simulation_validation.R

```
library(rlist)
library(Microsoft365R)

# specify the folder containing simulated data
output_dir <- "Output_no_extinction"

# specify the number of replicates
n_rep <- 3

# check if all simulations are complete
folder_names <- list.dirs(output_dir, recursive = F)
folder_names <- substr(folder_names, nchar(output_dir) + 2, nchar(output_dir) + 10)

# check whether all simulations are complete
complete_check <- matrix(NA, nrow = length(folder_names), ncol = n_rep)
rownames(complete_check) <- folder_names

for (row.id in rownames(complete_check)) {
  for (col.id in 1:ncol(complete_check)) {
    file_name <- paste(output_dir, "/", row.id, '/', as.character(col.id), '/', 'networks.rds', sep = '')
    complete_check[row.id, col.id] <- file.exists(file_name)
  }
}

mean(complete_check)

complete_check[1:100,]
complete_check[101:144,]
# complete_check[101:200,]
# complete_check[201:240,]


# check whether the "generations" file is properly stored in OneDrive
od <- NULL
od <- get_business_onedrive()

generations_check <- matrix(NA, nrow = length(folder_names), ncol = n_rep)
rownames(generations_check) <- folder_names

for (row.id in rownames(generations_check)) {
  print(row.id)
  for (col.id in 1:ncol(generations_check)) {
    sim_rep_name <- paste(output_dir, "/", row.id, '/', as.character(col.id), sep = '')
    file_list <- od$list_items(paste("model_rcpp_hpc_OneDrive/", sim_rep_name, sep = '')    )
    
    generations_check[row.id, col.id] <- as.logical(sum(file_list == "generations.rds"))
  }
}

mean(generations_check)

generations_check[1:100,]
generations_check[101:144,]
# generations_check[101:200,]
# generations_check[201:240,]


# change the save file if needed
save <- list.load("Output_extinction/aa1011AMF/comp_step.rds")
save$
list.save(save, "Output_extinction/aa1011AMF/comp_step.rds")


#############################################################################################
# check whether enough species have evolved in each simulation
host_sp_count <- matrix(NA, nrow = length(folder_names), ncol = n_rep)
rownames(host_sp_count) <- folder_names

for (row.id in rownames(host_sp_count)) {
  print(row.id)
  for (col.id in 1:ncol(host_sp_count)) {
    sp_count <- NA # find the minimum number of species in the last 100 generations
    file_name <- paste(output_dir, "/", row.id, '/', as.character(col.id), '/', 'pop_spec_host.rds', sep = '')
    pop_spec_host <- list.load(file_name)
    for(gen.id in (length(pop_spec_host)-99) : length(pop_spec_host)){
      sp_count[gen.id] <- length(pop_spec_host[[gen.id]]$host_spec_list)
    }
    host_sp_count[row.id, col.id] <- min(sp_count, na.rm = T)
  }
}
host_sp_count[1:100,]
host_sp_count[101:144,]
#host_sp_count[101:200,]
#host_sp_count[201:240,]


non_sp_count <- matrix(NA, nrow = length(folder_names), ncol = n_rep)
rownames(non_sp_count) <- folder_names

for (row.id in rownames(non_sp_count)) {
  print(row.id)
  for (col.id in 1:ncol(non_sp_count)) {
    sp_count <- NA # find the minimum number of species in the last 100 generations
    file_name <- paste(output_dir, "/", row.id, '/', as.character(col.id), '/', 'pop_spec_non.rds', sep = '')
    pop_spec_non <- list.load(file_name)
    for(gen.id in (length(pop_spec_non)-99) : length(pop_spec_non)){
      sp_count[gen.id] <- length(pop_spec_non[[gen.id]]$non_spec_list)
    }
    non_sp_count[row.id, col.id] <- min(sp_count, na.rm = T)
  }
}
non_sp_count[1:100,]
non_sp_count[101:144,]
#non_sp_count[101:200,]
#non_sp_count[201:240,]
```
