## Supplementary material for "Coevolution-induced stabilizing and destabilizing selection shapes species richness in clade co-diversification": File S1: trait_single_cell_zoom_in_out.R

```
library(ggplot2)
library(rlist)
library(Microsoft365R)

library(tidyverse)
#library(tidyr)
#install.packages("factoextra")
library(factoextra)
#install.packages("corrplot")
library(corrplot)
library(ggpubr)
#install.packages("ggthemes")
library(ggthemes)
#install.packages("egg")
library(egg)
library(Rarity)

library(plot.matrix)

theme_set(
  theme_classic() +
    theme(legend.position = "none")
)

#output_dir <- "Output_extinction"

# specify the color palette
greyscale <- c("gray90", "gray45", "gray0")

sim_nam <- "aa0000FFF" # this can be anything
source("basic_functions.R")

n_rep <- 3
n_gen <- 2500
n_dimension <- 7

# specify the name of the output folder
#output_dir <- "Output_no_extinction"

# initialize gen ids
no_generations <- 1:n_gen
#rep_id <- 1:n_rep

# initialize lists for storing the panels 
panel_list <- NULL
panel_list_host <- NULL
panel_list_non <- NULL

# initialize lists for storing the matrices
matrix_list_host <- NULL
matrix_list_non <- NULL

# create a vector storing all simulation ids
rep_id <- NULL
for (pos1 in c('a', 's')[1]) {
  for (pos2 in c('a', 's')[1]) {
    for (pos3 in c('0', '1')[1]) {
      for (pos6 in c('0', '1')[1]) {
        for (pos_ex in c('Output_no_extinction', 'Output_extinction')[1]) {
          for (pos_sim in 1:n_rep) {
            rep_id <- c(rep_id, paste(pos1, pos2, pos3, pos6, pos_ex, pos_sim, sep = ''))
          }
        }
      }
    }
  }
}

# initialize a vector containing all possible scenarios
# scenarios <- c(
#   "aa0110ADD", # Trait difference (+/-) Trait difference (+)
#   "aa0110ADM", # Trait difference (+/-) Trait matching (+)
#   "aa0010ADF", # Trait difference (+/-) Unaffected (0)
#   "aa0110MMD", # Trait matching (+) Trait difference (+)
#   "aa0110MMM", # Trait matching (+) Trait matching (+)
#   "aa0010MMF", # Trait matching (+) Unaffected (0)
#   "aa0110AMD", # Trait matching (-) Trait difference (+)
#   "aa0110AMM", # Trait matching (-) Trait matching (+)
#   "aa0010AMF", # Trait matching (-) Unaffected (0)
#   "aa0100FFD", # Unaffected (0) Trait difference (+)
#   "aa0100FFM", # Unaffected (0) Trait matching (+)
#   "aa0000FFF" # Unaffected (0) Unaffected (0)
# )

 scenarios <- c(
#   "01MDF", # Trait difference (+/-) Trait difference (+)
#   "01ADF"#, # Trait difference (+/-) Trait difference (+)
   "11MDD", # Trait difference (+/-) Trait difference (+)
   "01MDF", # Trait difference (+/-) Unaffected (0)  
   "10FFD", # Unaffected (0) Trait difference (+)  
   
   "11MMM", # Trait matching (+) Trait matching (+)  
   "01MMF", # Trait matching (+) Unaffected (0)
   "10FFM", # Unaffected (0) Trait matching (+)  
   
   "11AMM", # Trait matching (-) Trait matching (+) 
   "01AMF", # Trait matching (-) Unaffected (0)  
   #"10FFM", # Unaffected (0) Trait matching (+)   
   
   "00FFF" # Unaffected (0) Unaffected (0)
  
  #"11ADM", # Trait difference (+/-) Trait matching (+)
  
  #"11MMD", # Trait matching (+) Trait difference (+)
  
  
  #"11AMD", # Trait matching (-) Trait difference (+)
)

for (scen in scenarios) {
  
  print(length(panel_list))
  
  rep <- rep_id[2]
  
  output_dir <- substr(rep, 5, (nchar(rep)-1))
  alt_id <- paste(substr(rep, 1, 3), substr(scen, 1, 2), substr(rep, 4, 4), substr(scen, 3, 5), sep = '')
  sim_dir <- paste(output_dir, "/", alt_id, sep = '')
  sim <- substr(rep, nchar(rep), nchar(rep))
  
  od <- NULL
  while (length(od) == 0) {
    try( # try again if it fails
      od <- get_business_onedrive(),
      TRUE
    )  
  }
  
  generations <- NULL
  file_name <- paste(sim_dir, "/", as.character(sim), "/generations.rds", sep = "")
  while (length(generations) == 0) {
    try( # try again if it fails
      od$download_file(paste("model_rcpp_hpc_OneDrive/", file_name, sep = ''), file_name, overwrite = T),
      TRUE
    )
    try( # try again if it fails
      generations <- list.load(file_name),
      TRUE
    )    
  }
  print('1')
  unlink(file_name)
  
  ######################################################################################3
  # the evolution of host trait
  print("calculating host trait")
  global_host_trait_list <- NULL
  
  #for (rep in rep_id) {
    
    # initialize the mean, lower 95%, and upper 95%
    host_trait_mean_global <- rep(0, n_gen)
    host_trait_mean_global_low <- rep(0, n_gen)
    host_trait_mean_global_high <- rep(0, n_gen)
    
    for (g in 1:n_gen){
      #print(g)
      global <- rep(NA, length(generations[[g]]$host_list))
      for (i in 1:length(generations[[g]]$host_list)) {
        if(identical(generations[[g]]$host_list[[i]]$site.id, c(ceiling(n_dimension/2),ceiling(n_dimension/2)))){ # focus on the cetral cell only
          global[i] <- get_eco_trait_val(generations[[g]]$host_list[[i]])
        }
      }
      global <- global[!is.na(global)]
      host_trait_mean_global[g] <- mean(global)
      if(length(unique(global))==1){
        host_trait_mean_global_low[g] <- host_trait_mean_global[g]
        host_trait_mean_global_high[g] <- host_trait_mean_global[g]
      }
      else{
        host_trait_mean_global_low[g] <- t.test(global)$conf.int[1]
        host_trait_mean_global_high[g] <- t.test(global)$conf.int[2]   
      }

    }
    
  #}
  
  ######################################################################################
  # the evolution of non-host trait
    print("calculating non trait")
    global_non_trait_list <- NULL
    
    #for (rep in rep_id) {
    
    # initialize the mean, lower 95%, and upper 95%
    non_trait_mean_global <- rep(0, n_gen)
    non_trait_mean_global_low <- rep(0, n_gen)
    non_trait_mean_global_high <- rep(0, n_gen)
    
    for (g in 1:n_gen){
      #print(g)
      global <- rep(NA, length(generations[[g]]$non_list))
      for (i in 1:length(generations[[g]]$non_list)) {
        if(identical(generations[[g]]$non_list[[i]]$site.id, c(ceiling(n_dimension/2),ceiling(n_dimension/2)))){ # focus on the cetral cell only
          global[i] <- get_eco_trait_val(generations[[g]]$non_list[[i]])
        }
      }
      global <- global[!is.na(global)]
      non_trait_mean_global[g] <- mean(global)
      if(length(unique(global))==1){
        non_trait_mean_global_low[g] <- non_trait_mean_global[g]
        non_trait_mean_global_high[g] <- non_trait_mean_global[g]
      }
      else{
        non_trait_mean_global_low[g] <- t.test(global)$conf.int[1]
        non_trait_mean_global_high[g] <- t.test(global)$conf.int[2]   
      }
      
    }
    
    #}
  
  
  #############################################################################################
  # Save all Trait values in a dataset
  data <- data.frame(no_generations, host_trait_mean_global, host_trait_mean_global_low, host_trait_mean_global_high, non_trait_mean_global, non_trait_mean_global_low, non_trait_mean_global_high)
  
  #############################################################################################
  # Plotting
  trait_global <- ggplot(data, aes(no_generations, host_trait_mean_global)) +    # ggplot2 plot without confidence band
    geom_line(color = "blue", alpha = 0.4, linewidth=1)
  
  panel <- trait_global +
    #ggtitle("all simulations") +
    xlab("") + ylab("") +                              # Add confidence intervals
    #ylim(-30,150) +
    #geom_ribbon(aes(ymin = host_trait_mean_global_low, ymax = host_trait_mean_global_high), fill = "blue", alpha = 0.2) +
    geom_line(aes(y = non_trait_mean_global), color = "orange", alpha = 0.4, linewidth=1)+
    #geom_ribbon(aes(ymin = non_trait_mean_global_low, ymax = non_trait_mean_global_high), fill = "red", alpha = 0.2) +
    theme(legend.position="none")
  
  panel_list[[length(panel_list) + 1]] <- panel
  
  # specify what range to use for plotting
  if(length(panel_list_host) == 6){
    rang_host <- 40
    rang_non <- 40
  }else{
    rang_host <- 8
    rang_non <- 10
  }
  
  panel_host <- ggplot(data[(nrow(data) - 200) : nrow(data),], aes(no_generations, host_trait_mean_global)) +    # ggplot2 plot without confidence band
    xlab("") + ylab("") + 
    ylim(min(data[(nrow(data) - 200) : nrow(data),]$host_trait_mean_global), min(data[(nrow(data) - 200) : nrow(data),]$host_trait_mean_global) + rang_host) +
    theme(axis.text.x = element_text(angle = 90)) +
    geom_line(color = "blue", alpha = 0.4, linewidth=1)
    
  panel_list_host[[length(panel_list_host) + 1]] <- panel_host
  
  panel_non <- ggplot(data[(nrow(data) - 200) : nrow(data),], aes(no_generations, non_trait_mean_global)) +    # ggplot2 plot without confidence band
    xlab("") + ylab("") + 
    ylim(min(data[(nrow(data) - 200) : nrow(data),]$non_trait_mean_global), min(data[(nrow(data) - 200) : nrow(data),]$non_trait_mean_global) + rang_non) +
    theme(axis.text.x = element_text(angle = 90)) +
    geom_line(color = "orange", alpha = 0.4, linewidth=1)
    
  panel_list_non[[length(panel_list_non) + 1]] <- panel_non

}

list.save(panel_list, paste(output_dir, '/panel_list_20230614.rds', sep = ''))


########################################################################
# Plot host trait deviation across geographic space
# for (scen in scenarios) {
#   rep <- rep_id[1]
#   output_dir <- substr(rep, 5, (nchar(rep)-1))
#   alt_id <- paste(substr(rep, 1, 3), substr(scen, 1, 2), substr(rep, 4, 4), substr(scen, 3, 5), sep = '')
#   sim_dir <- paste(output_dir, "/", alt_id, sep = '')
#   sim <- substr(rep, nchar(rep), nchar(rep))
# 
#   od <- NULL
#   while (length(od) == 0) {
#     try( # try again if it fails
#       od <- get_business_onedrive(),
#       TRUE
#     )
#   }
# 
#   generations <- NULL
#   file_name <- paste(sim_dir, "/", as.character(sim), "/generations.rds", sep = "")
#   while (length(generations) == 0) {
#     try( # try again if it fails
#       od$download_file(paste("model_rcpp_hpc_OneDrive/", file_name, sep = ''), file_name, overwrite = T),
#       TRUE
#     )
#     try( # try again if it fails
#       generations <- list.load(file_name),
#       TRUE
#     )
#   }
#   print('3')
#   unlink(file_name)
# 
#   generation <- generations[[n_gen]]
#   host_list <- generation$host_list
# 
#   # initialize a list recording individual Trait deviations for each site
#   trait_list_host <- as.list(rep(NA, n_dimension^2))
# 
#   # record Trait deviations of hosts in the list
#   for (host_id in 1:length(host_list)) {
#     site_row <- host_list[[host_id]]$site.id[1]
#     site_col <- host_list[[host_id]]$site.id[2]
# 
#     trait_list_host[[(site_row-1) * n_dimension + site_col]][length(trait_list_host[[(site_row-1) * n_dimension + site_col]]) + 1]<- get_eco_trait_val(host_list[[host_id]])
#   }
# 
#   # fill in a matrix using the list
#   matrix_host <- matrix(NA, nrow = n_dimension, ncol = n_dimension)
#   for (site_id in 1:length(trait_list_host)) {
#     col_id <- site_id%%n_dimension
#     if(col_id == 0) col_id <- n_dimension
#     row_id <- ceiling(site_id/n_dimension)
#     matrix_host[row_id, col_id] <- mean(trait_list_host[[site_id]], na.rm=T)
#   }
# 
#   matrix_host_sd <- matrix(NA, nrow = n_dimension, ncol = n_dimension)
#   for (site_id in 1:length(trait_list_host)) {
#     col_id <- site_id%%n_dimension
#     if(col_id == 0) col_id <- n_dimension
#     row_id <- ceiling(site_id/n_dimension)
#     matrix_host_sd[row_id, col_id] <- sd(trait_list_host[[site_id]], na.rm=T)
#   }
# 
#   matrix_host_scaled_original <- (matrix_host - mean(matrix_host)) / mean(matrix_host_sd)
#   print(range(matrix_host_scaled_original))
# 
#   matrix_list_host[[length(matrix_list_host) + 1]] <- matrix_host_scaled_original
# 
# }
# 
# 
# # plot and save
# #panel_list_host[[length(panel_list_host) + 1]] <- plot(matrix_host_scaled, key = NULL, axis.col = NULL, axis.row = NULL, main = '', xlab = '', ylab = '', breaks = c(-2, -1.5, -1, -0.5, 0, 0.5, 1, 1.5, 2))
# for (plot_id in 1:length(scenarios)) {
# 
#   matrix_host_scaled_original <- matrix_list_host[[plot_id]]
# 
#   matrix_host_scaled <- as.data.frame(matrix_host_scaled_original)
#   matrix_host_scaled <- rownames_to_column(matrix_host_scaled, "f_id")
#   matrix_host_scaled <- pivot_longer(matrix_host_scaled, -c(f_id), names_to = "samples", values_to = "Trait deviation")
# 
#   matrix_host_scaled$`Trait deviation`[matrix_host_scaled$`Trait deviation` > -1.65 & matrix_host_scaled$`Trait deviation` < -0.55] <- -1.5
#   matrix_host_scaled$`Trait deviation`[matrix_host_scaled$`Trait deviation` > -0.55 & matrix_host_scaled$`Trait deviation` < 0.55] <- -0.5
#   matrix_host_scaled$`Trait deviation`[matrix_host_scaled$`Trait deviation` > 0.55 & matrix_host_scaled$`Trait deviation` < 1.65] <- 0.5
#   #matrix_host_scaled$`Trait deviation`[matrix_host_scaled$`Trait deviation` > 1 & matrix_host_scaled$`Trait deviation` < 2] <- 1.5
# 
#   matrix_host_scaled$`Trait deviation` <- as.character(matrix_host_scaled$`Trait deviation`)
# 
#   matrix_host_scaled$`Trait deviation`[matrix_host_scaled$`Trait deviation` == '-1.5'] <- '-1.65 ~ -0.55'
#   matrix_host_scaled$`Trait deviation`[matrix_host_scaled$`Trait deviation` == '-0.5'] <- '-0.55 ~ 0.55'
#   matrix_host_scaled$`Trait deviation`[matrix_host_scaled$`Trait deviation` == '0.5'] <- '0.55 ~ 1.65'
#   #matrix_host_scaled$`Trait deviation`[matrix_host_scaled$`Trait deviation` == '1.5'] <- '1 ~ -2'
# 
#   matrix_host_scaled$`Trait deviation` <- factor(matrix_host_scaled$`Trait deviation`, levels = c('-1.65 ~ -0.55', '-0.55 ~ 0.55', '0.55 ~ 1.65'))
# 
#   greyscale_sub <- NULL
#   for (color_id in 1:length(sort(unique(matrix_host_scaled$`Trait deviation`)))) {
#     greyscale_sub <- c(greyscale_sub, greyscale[which(c('-1.65 ~ -0.55', '-0.55 ~ 0.55', '0.55 ~ 1.65') == sort(unique(matrix_host_scaled$`Trait deviation`))[color_id])])
#   }
# 
#   panel_host <- ggplot(matrix_host_scaled, aes(x=samples, y=f_id, fill=`Trait deviation`))+
#     geom_tile()+
#     theme_void()+
#     theme(legend.position = "none")+
#     xlab('')+
#     ylab('')+
#     scale_fill_manual(values=greyscale_sub)
#     #continuous_scale("Highway mpg", breaks = c(-20, 0, 20))+
#     #scale_fill_viridis_c()
#     #scale_fill_continuous(breaks = c(4, 5, 6), labels = c(10000, 100000, 1000000))
#     #scale_fill_viridis_c(begin = 0.5 - 0.5 * (min(matrix_host_scaled_original) * (-1.39) - 1.21), end = 0.5 + 0.5 * (max(matrix_host_scaled_original) * 1.17 -0.7))
#     #scale_fill_viridis_d(option = "C", )
# 
#   panel_list_host[[length(panel_list_host) + 1]] <- panel_host
# 
# }

list.save(panel_list_host, 'panel_list_host_20230614.rds')


########################################################################
# Plot example non trait distribution across landscape
# for (scen in scenarios) {
#   rep <- rep_id[1]
#   output_dir <- substr(rep, 5, (nchar(rep)-1))
#   alt_id <- paste(substr(rep, 1, 3), substr(scen, 1, 2), substr(rep, 4, 4), substr(scen, 3, 5), sep = '')
#   sim_dir <- paste(output_dir, "/", alt_id, sep = '')
#   sim <- substr(rep, nchar(rep), nchar(rep))
# 
#   od <- NULL
#   while (length(od) == 0) {
#     try( # try again if it fails
#       od <- get_business_onedrive(),
#       TRUE
#     )
#   }
# 
#   generations <- NULL
#   file_name <- paste(sim_dir, "/", as.character(sim), "/generations.rds", sep = "")
#   while (length(generations) == 0) {
#     try( # try again if it fails
#       od$download_file(paste("model_rcpp_hpc_OneDrive/", file_name, sep = ''), file_name, overwrite = T),
#       TRUE
#     )
#     try( # try again if it fails
#       generations <- list.load(file_name),
#       TRUE
#     )
#   }
#   print('4')
#   unlink(file_name)
# 
#   generation <- generations[[n_gen]]
#   non_list <- generation$non_list
# 
#   # initialize a list recording individual Trait deviations for each site
#   trait_list_non <- as.list(rep(NA, n_dimension^2))
# 
#   # record Trait deviations of nons in the list
#   for (non_id in 1:length(non_list)) {
#     site_row <- non_list[[non_id]]$site.id[1]
#     site_col <- non_list[[non_id]]$site.id[2]
# 
#     trait_list_non[[(site_row-1) * n_dimension + site_col]][length(trait_list_non[[(site_row-1) * n_dimension + site_col]]) + 1]<- get_eco_trait_val(non_list[[non_id]])
#   }
# 
#   # fill in a matrix using the list
#   matrix_non <- matrix(NA, nrow = n_dimension, ncol = n_dimension)
#   for (site_id in 1:length(trait_list_non)) {
#     col_id <- site_id%%n_dimension
#     if(col_id == 0) col_id <- n_dimension
#     row_id <- ceiling(site_id/n_dimension)
#     matrix_non[row_id, col_id] <- mean(trait_list_non[[site_id]], na.rm=T)
#   }
# 
#   matrix_non_sd <- matrix(NA, nrow = n_dimension, ncol = n_dimension)
#   for (site_id in 1:length(trait_list_non)) {
#     col_id <- site_id%%n_dimension
#     if(col_id == 0) col_id <- n_dimension
#     row_id <- ceiling(site_id/n_dimension)
#     matrix_non_sd[row_id, col_id] <- sd(trait_list_non[[site_id]], na.rm=T)
#   }
# 
#   matrix_non_scaled_original <- (matrix_non - mean(matrix_non)) / mean(matrix_non_sd)
#   print(range(matrix_non_scaled_original))
# 
#   matrix_list_non[[length(matrix_list_non) + 1]] <- matrix_non_scaled_original
# 
# }
# 
# 
# # plot and save
# #panel_list_non[[length(panel_list_non) + 1]] <- plot(matrix_non_scaled, key = NULL, axis.col = NULL, axis.row = NULL, main = '', xlab = '', ylab = '', breaks = c(-2, -1.5, -1, -0.5, 0, 0.5, 1, 1.5, 2))
# for (plot_id in 1:length(scenarios)) {
# 
#   matrix_non_scaled_original <- matrix_list_non[[plot_id]]
# 
#   matrix_non_scaled <- as.data.frame(matrix_non_scaled_original)
#   matrix_non_scaled <- rownames_to_column(matrix_non_scaled, "f_id")
#   matrix_non_scaled <- pivot_longer(matrix_non_scaled, -c(f_id), names_to = "samples", values_to = "Trait deviation")
# 
#   matrix_non_scaled$`Trait deviation`[matrix_non_scaled$`Trait deviation` > -1.65 & matrix_non_scaled$`Trait deviation` < -0.55] <- -1.5
#   matrix_non_scaled$`Trait deviation`[matrix_non_scaled$`Trait deviation` > -0.55 & matrix_non_scaled$`Trait deviation` < 0.55] <- -0.5
#   matrix_non_scaled$`Trait deviation`[matrix_non_scaled$`Trait deviation` > 0.55 & matrix_non_scaled$`Trait deviation` < 1.65] <- 0.5
#   #matrix_non_scaled$`Trait deviation`[matrix_non_scaled$`Trait deviation` > 1 & matrix_non_scaled$`Trait deviation` < 2] <- 1.5
# 
#   matrix_non_scaled$`Trait deviation` <- as.character(matrix_non_scaled$`Trait deviation`)
# 
#   matrix_non_scaled$`Trait deviation`[matrix_non_scaled$`Trait deviation` == '-1.5'] <- '-1.65 ~ -0.55'
#   matrix_non_scaled$`Trait deviation`[matrix_non_scaled$`Trait deviation` == '-0.5'] <- '-0.55 ~ 0.55'
#   matrix_non_scaled$`Trait deviation`[matrix_non_scaled$`Trait deviation` == '0.5'] <- '0.55 ~ 1.65'
#   #matrix_non_scaled$`Trait deviation`[matrix_non_scaled$`Trait deviation` == '1.5'] <- '1 ~ -2'
# 
#   matrix_non_scaled$`Trait deviation` <- factor(matrix_non_scaled$`Trait deviation`, levels = c('-1.65 ~ -0.55', '-0.55 ~ 0.55', '0.55 ~ 1.65'))
# 
#   greyscale_sub <- NULL
#   for (color_id in 1:length(sort(unique(matrix_non_scaled$`Trait deviation`)))) {
#     greyscale_sub <- c(greyscale_sub, greyscale[which(c('-1.65 ~ -0.55', '-0.55 ~ 0.55', '0.55 ~ 1.65') == sort(unique(matrix_non_scaled$`Trait deviation`))[color_id])])
#   }
# 
#   panel_non <- ggplot(matrix_non_scaled, aes(x=samples, y=f_id, fill=`Trait deviation`))+
#     geom_tile()+
#     theme_void()+
#     theme(legend.position = "none")+
#     xlab('')+
#     ylab('')+
#     scale_fill_manual(values=greyscale_sub)
#   #continuous_scale("Highway mpg", breaks = c(-20, 0, 20))+
#   #scale_fill_viridis_c()
#   #scale_fill_continuous(breaks = c(4, 5, 6), labels = c(10000, 100000, 1000000))
#   #scale_fill_viridis_c(begin = 0.5 - 0.5 * (min(matrix_non_scaled_original) * (-1.39) - 1.21), end = 0.5 + 0.5 * (max(matrix_non_scaled_original) * 1.17 -0.7))
#   #scale_fill_viridis_d(option = "C", )
# 
#   panel_list_non[[length(panel_list_non) + 1]] <- panel_non
# 
# }

list.save(panel_list_non, 'panel_list_non_20230614.rds')


#########################################################################
# read in the list
#########################################################################
# read in the list
panel_list <- list.load(paste('Output_no_extinction/panel_list_20230614.rds', sep = ''))
panel_list_host <- list.load('panel_list_host_20230614.rds')
panel_list_non <- list.load('panel_list_non_20230614.rds')

# create a blank panel
blank <- panel_list[[1]] + lims(x = c(0,0), y = c(0,0)) + theme_void()

# add titles and axes to panels
panel_list[[1]] <- panel_list[[1]] + ggtitle('(a) Trait difference\n(mutualistic or antagonistic)\nSymmetric') + ylab('Trait value') + ylim(-25, 100)
panel_list_host[[1]] <- panel_list_host[[1]] + ggtitle('Host') + theme(plot.title=element_text(size=10))
panel_list_non[[1]] <- panel_list_non[[1]] + ggtitle('Dependent') + theme(plot.title=element_text(size=10))

host_non_1 <- ggpubr::ggarrange(
  blank,
  panel_list_host[[1]], blank,
  panel_list_non[[1]], blank,
  ncol = 1, nrow = 5, heights = c(1, 2, 0, 2, 0)
)

panel_1 <- ggpubr::ggarrange(
  panel_list[[1]], host_non_1,
  ncol = 2, nrow = 1, widths = c(3,2)
)

#panel_1 <- panel_list[[1]]


panel_list[[2]] <- panel_list[[2]] + ggtitle('(b) Trait difference\n(mutualistic or antagonistic)\nWeak for dependents') + ylim(-25, 100)
panel_list_host[[2]] <- panel_list_host[[2]] + ggtitle('Host') + theme(plot.title=element_text(size=10))
panel_list_non[[2]] <- panel_list_non[[2]] + ggtitle('Dependent') + theme(plot.title=element_text(size=10))

host_non_2 <- ggpubr::ggarrange(
  blank,
  panel_list_host[[2]], blank,
  panel_list_non[[2]], blank,
  ncol = 1, nrow = 5, heights = c(1, 2, 0, 2, 0)
)

panel_2 <- ggpubr::ggarrange(
  panel_list[[2]], host_non_2,
  ncol = 2, nrow = 1, widths = c(3,2)
)
#panel_2 <- panel_list[[2]]

panel_list[[3]] <- panel_list[[3]] + ggtitle('(c) Trait difference\n(mutualistic or antagonistic)\nWeak for hosts') + ylim(-25, 100)
panel_list_host[[3]] <- panel_list_host[[3]] + ggtitle('Host') + theme(plot.title=element_text(size=10))
panel_list_non[[3]] <- panel_list_non[[3]] + ggtitle('Dependent') + theme(plot.title=element_text(size=10))

host_non_3 <- ggpubr::ggarrange(
  blank,
  panel_list_host[[3]], blank,
  panel_list_non[[3]], blank,
  ncol = 1, nrow = 5, heights = c(1, 2, 0, 2, 0)
)

panel_3 <- ggpubr::ggarrange(
  panel_list[[3]], host_non_3,
  ncol = 2, nrow = 1, widths = c(3,2)
)
#panel_3 <- panel_list[[3]]

panel_list[[4]] <- panel_list[[4]] + ggtitle('(d) Trait matching\n(mutualistic)\nSymmetric') + ylab('Trait value') + ylim(-25, 100)
panel_list_host[[4]] <- panel_list_host[[4]] + ggtitle('Host') + theme(plot.title=element_text(size=10))
panel_list_non[[4]] <- panel_list_non[[4]] + ggtitle('Dependent') + theme(plot.title=element_text(size=10))

host_non_4 <- ggpubr::ggarrange(
  blank,
  panel_list_host[[4]], blank,
  panel_list_non[[4]], blank,
  ncol = 1, nrow = 5, heights = c(1, 2, 0, 2, 0)
)

panel_4 <- ggpubr::ggarrange(
  panel_list[[4]], host_non_4,
  ncol = 2, nrow = 1, widths = c(3,2)
)
#panel_4 <- panel_list[[4]]

panel_list[[5]] <- panel_list[[5]] + ggtitle('(e) Trait matching\n(mutualistic)\nWeak for dependents') + ylim(-25, 100)
panel_list_host[[5]] <- panel_list_host[[5]] + ggtitle('Host') + theme(plot.title=element_text(size=10))
panel_list_non[[5]] <- panel_list_non[[5]] + ggtitle('Dependent') + theme(plot.title=element_text(size=10))

host_non_5 <- ggpubr::ggarrange(
  blank,
  panel_list_host[[5]], blank,
  panel_list_non[[5]], blank,
  ncol = 1, nrow = 5, heights = c(1, 2, 0, 2, 0)
)

panel_5 <- ggpubr::ggarrange(
  panel_list[[5]], host_non_5,
  ncol = 2, nrow = 1, widths = c(3,2)
)
#panel_5 <- panel_list[[5]]

panel_list[[6]] <- panel_list[[6]] + ggtitle('(f) Trait matching\n(mutualistic or antagonistic)\nWeak for hosts') + ylim(-25, 100)
panel_list_host[[6]] <- panel_list_host[[6]] + ggtitle('Host') + theme(plot.title=element_text(size=10))
panel_list_non[[6]] <- panel_list_non[[6]] + ggtitle('Dependent') + theme(plot.title=element_text(size=10))

host_non_6 <- ggpubr::ggarrange(
  blank,
  panel_list_host[[6]], blank,
  panel_list_non[[6]], blank,
  ncol = 1, nrow = 5, heights = c(1, 2, 0, 2, 0)
)

panel_6 <- ggpubr::ggarrange(
  panel_list[[6]], host_non_6,
  ncol = 2, nrow = 1, widths = c(3,2)
)
#panel_6 <- panel_list[[6]]

panel_list[[7]] <- panel_list[[7]] + ggtitle('(g) Trait matching\n(antagonistic)\nSymmetric') + xlab('Generation') + ylab('Trait value') + ylim(-25, 100)
panel_list_host[[7]] <- panel_list_host[[7]] + ggtitle('Host') + theme(plot.title=element_text(size=10))
panel_list_non[[7]] <- panel_list_non[[7]] + ggtitle('Dependent') + theme(plot.title=element_text(size=10))

host_non_7 <- ggpubr::ggarrange(
  blank,
  panel_list_host[[7]], blank,
  panel_list_non[[7]], blank,
  ncol = 1, nrow = 5, heights = c(1, 2, 0, 2, 0)
)

panel_7 <- ggpubr::ggarrange(
  panel_list[[7]], host_non_7,
  ncol = 2, nrow = 1, widths = c(3,2)
)
#panel_7 <- panel_list[[7]]

panel_list[[8]] <- panel_list[[8]] + ggtitle('(h) Trait matching\n(antagonistic)\nWeak for dependents') + xlab('Generation') + ylim(-25, 100)
panel_list_host[[8]] <- panel_list_host[[8]] + ggtitle('Host') + theme(plot.title=element_text(size=10))
panel_list_non[[8]] <- panel_list_non[[8]] + ggtitle('Dependent') + theme(plot.title=element_text(size=10))

host_non_8 <- ggpubr::ggarrange(
  blank,
  panel_list_host[[8]], blank,
  panel_list_non[[8]], blank,
  ncol = 1, nrow = 5, heights = c(1, 2, 0, 2, 0)
)

panel_8 <- ggpubr::ggarrange(
  panel_list[[8]], host_non_8,
  ncol = 2, nrow = 1, widths = c(3,2)
)
#panel_8 <- panel_list[[8]]

panel_list[[9]] <- panel_list[[9]] + ggtitle('\n\n(i) No coevolution') + xlab('Generation') + xlab('Generation') + ylim(-25, 100)
panel_list_host[[9]] <- panel_list_host[[9]] + ggtitle('Host') + theme(plot.title=element_text(size=10))
panel_list_non[[9]] <- panel_list_non[[9]] + ggtitle('Dependent') + theme(plot.title=element_text(size=10))

host_non_9 <- ggpubr::ggarrange(
  blank,
  panel_list_host[[9]], blank,
  panel_list_non[[9]], blank,
  ncol = 1, nrow = 5, heights = c(1, 2, 0, 2, 0)
)

panel_9 <- ggpubr::ggarrange(
  panel_list[[9]], host_non_9,
  ncol = 2, nrow = 1, widths = c(3,2)
)
#panel_9 <- panel_list[[9]]

# create a legend panel
# legend <- panel_list_host[[1]] + lims(x = c(0,0), y = c(0,0))+ #scale_color_manual(values=c('royalblue4', 'skyblue'))+
#   theme_void()+
#   theme(legend.position = c(0.5,0.5),
#         legend.key.size = unit(0.8, "cm"),
#         legend.text = element_text(size = 13),
#         legend.title = element_text(size = 15, face = "bold")
#   )+
#   ggtitle('')+
#   xlim("Fair","Ideal", "Good")+
#   ylim("Fair","Ideal", "Good")
#guides(colour = guide_legend(override.aes = list(size=5)))#+
#scale_fill_manual(greyscale)

# arrange and plot
# trait_evo <- egg::ggarrange(
#   panel_1, panel_2, panel_3, blank,
#   panel_4, panel_5, panel_6, legend,
#   panel_7, panel_8, panel_9, blank,
#   ncol = 4, nrow = 3, widths = c(2,2,2,1.5), heights = c(1,1,1)
# )

# arrange and plot
trait_evo <- egg::ggarrange(
  panel_1, panel_2, panel_3,# blank,
  panel_4, panel_5, panel_6,# legend,
  panel_7, panel_8, panel_9,# blank,
  ncol = 3, nrow = 3, widths = c(2,2,2), heights = c(1,1,1)
)

pdf("trait_evo_single_24_20230706.pdf", width = 11, height = 12)
trait_evo
dev.off()
```
